# Prolonged heat stress in *Brassica napus* during flowering negatively impacts yield and alters glucosinolate and sugars metabolism

**DOI:** 10.1101/2024.04.01.587615

**Authors:** Mariam Kourani, Maria Anastasiadi, John P. Hammond, Fady Mohareb

## Abstract

Oilseed rape (*Brassica napus*), one of the most important sources of vegetable oil worldwide, is adversely impacted by heat wave-induced temperature stress especially during its yield determining reproductive stages. However, the underlying molecular and biochemical mechanisms are still poorly understood.

In this study, we investigated the transcriptomic and metabolomic responses to heat stress in *B. napus* plants exposed to a gradual increase of temperature reaching 30 °C in the day and 24 °C at night for a period of 6 days. High Performance Liquid Chromatography (HPLC) and Liquid Chromatography-Mass Spectrometry (LC-MS) was used to quantify the content of carbohydrates and glucosinolates respectively. RNA-Seq analysis of flower buds showed a total of 1892, 3253, 4553, 4165 and 1713 genes were differentially expressed at days 0, 1, 2, 6 and 12 after treatment respectively.

Results showed that heat stress reduced yield and altered oil composition. Heat stress also increased the content of carbohydrate (Glucose, Fructose and Sucrose) and aliphatic glucosinolates (Gluconapin, Progoitrin) in the leaves but decreased the content of the indolic glucosinolate (Glucobrassicin). Heat treatment resulted in down regulation of genes involved in respiratory metabolism namely Glycolysis, Pentose Phosphate Pathway, Citrate Cycle and Oxidative Phosphorylation especially after 48 hrs of heat stress. Other down regulated genes mapped to sugars transporters, nitrogen transport and storage, cell wall modification and methylation. In contrast, up regulated genes mapped to small heat shock proteins (sHSP20) and other heat shock factors that play important roles in thermotolerance. Furthermore, two genes were chosen from the pathways involved in the heat stress response to further examine their expression using real time RT-qPCR.

Taken together, the results of this study demonstrated that the changes in carbohydrates and glucosinolates metabolism and transport under heat stress is one of the response mechanisms employed by the plant under heat stress conditions. Moreover, the transcriptomic data will be useful for further understanding the molecular mechanisms of *B. napus* plant tolerance to heat stress and provide a basis for breeding heat-tolerant varieties.

## 1. INTRODUCTION

Over the last decade, climate change has led to more extreme climatic events, impacting crop productivity and threatening global food security (Ismaili et al., 2015; Lamaoui et al., 2018). These extreme events include periods of high temperature stress in the form of heatwaves (Goel et al., 2023), broadly defined as periods of excessively high temperature as compared to the local climate (Dikšaitytė et al., 2020). Previous studies have shown that heat stress can be detrimental during reproduction development due to the damage caused to plant organs and cellular structures (Hedhly 2011; Hinojosa et al., 2019, Goel et al., 2023). During reproduction, high temperature can induce irreversible structural and physiological changes in both male and female floral organs, leading to premature senescence (Lohani et al., 2020; Jagadish et al., 2021). High temperature stress can also decrease chlorophyll synthesis and disrupt photosynthesis and respiration, further reducing the yield potential of crop plants (Hasanuzzaman et al., 2014; Zandalinas et al., 2016). Thus, understanding the impact of heatwaves on crops and particularly during reproductive stages is essential to the development of varieties that can withstand these periods of elevated temperatures.

Traditionally, most heat stress experiments involve a heat shock, where plants are subjected to a high temperature within a very short time (10–15 °C above their optimal temperature, from several minutes to a few hours (Wahid et al, 2007). As a result, plant responses to heat shock have been well studied (Huang et al., 2019; Wang et al., 2020), but relatively little information exists on the responses to heatwaves, especially at the transcriptomic and metabolomic levels (Jin et al., 2011; Glaubitz et al., 2017).

In plants, heat shock reduces photosynthesis and respiratory metabolism, and increases antioxidant activity (Huang et al., 2019; Wang et al., 2020). Heat shock also negatively affects plant growth and productivity (Ismaili et al., 2015). While these experiments revealed the regulatory mechanisms in response to sudden heat stress (Seth et al, 2021; Ikram et al., 2022; Li et al., 2022), they do not fully represent the impact of temperature changes in the field during heatwave conditions. As a result, several studies employed prolonged warming experiments to study heat stress (Jin et al., 2011; Way & Yamori 2014; Glaubitz et al., 2017). In Arabidopsis, studies showed that plants exhibit different response patterns to prolonged warming (7 days) as compared to heat shocks (Jin et al., 2011; Wang et al., 2020). While prolonged warming led to a reduction in stomatal conductance, heat shock increased transpiration. Under both heat shock and prolonged heat stress there was an induction of antioxidant enzymes in Arabidopsis, however, these were significantly higher under the heat shock treatment compared to the prolonged heat treatment (Wang et al., 2020). Arabidopsis plants grown at 28/23°C (elevated by 5°C from ambient temperature) suffered from significant reduction in life span, total biomass, total weight of seeds, and leaf concentration of starch, chlorophyll, and proline (Jin et al., 2011). In rice, under prolonged elevated night temperature, transcriptomic analysis showed changes in cell wall, ABA signalling and secondary metabolism related genes, while metabolite profiles revealed a highly activated Citrate Cycle and enzyme activities (Glaubitz et al., 2017).

Oilseed rape (*Brassica napus* L.) is the second-largest source of oilseed after soybean and the third-largest source of vegetable oil worldwide (Lee et al., 2020). Cultivation and breeding practices have resulted in numerous genetically diverse lines with strong agronomic and adaptation traits (Sun et al., 2017). As a result, *B. napus* is widely cultivated around the world for food, biofuel and animal feed (Namazkar et al., 2016). Like other major temperate field crops, *B. napus* can be extremely sensitive to high temperature stress, especially if it occurs during flowering or seed development, threatening yields and quality (Angadi et al., 2000; Koscielny et al., 2018; Kourani et al., 2022). Studies in different Brassica species have found negative relationships between heat stress and seed yield and quality (Yu et al., 2014). For example, under elevated temperature scenarios, four *B. napus* cultivars experienced a significant reduction in seed biomass, resulting in a 58% decrease in the oil yield, and 77% decrease in the essential fatty acid C18:3-ω3 (Namazkar et al., 2016). Heat stress also reduced oil seed content and impaired carbohydrate incorporation into triacylglycerols in *B. napus* (Huang et al., 2019). Additionally, impairment of chlorophyll biosynthesis and disruption of the biochemical reactions of photosystems were exhibited in 10-day-old *B. napus* seedlings subjected to 38 °C (Hasanuzzaman et al., 2014). This was manifested through a significant reduction in chlorophyll and leaf relative water content as well as through an inefficient antioxidant defence system (Hasanuzzaman et al., 2014). Despite current advances in studying the effect of heat on different Brassica species, the impact of heatwaves on *B. napus,* especially during its yield determining reproductive stages, is still poorly understood (Kourani et al., 2022).

To address this, we investigated the transcriptomic and metabolomic responses to heat stress in *B. napus* under simulated field conditions. To achieve this, a heat treatment experiment simulating heatwave episode during the flowering stage of *B. napus* cv. Westar was conducted in a controlled environment. Global transcriptome profiling using RNA-Seq was employed to identify differentially expressed genes in flower buds of heat stressed plants (30 °C/24 °C day/night for a period of 6 days). This was complemented with HPLC and LC-MS analyses to detect changes in sugars and glucosinolates (GLS) concentration in leaves in response to the heat treatment.

Therefore, this study aims to provide insights into the impact of a future warmer climate on the important oil crop species *B. napus* during its reproductive stages.

## 2. MATERIALS AND METHODS

### 2.1 Plant materials and physiological parameters

Seeds of spring oilseed rape (*Brassica napus* L., cv. Westar) were surface-sterilized and sown into seed trays filled with a seed potting mix (Clover Peat, Dungannon, Northern Ireland). Trays were then placed in polyethylene tunnels for four weeks at the Crops and Environment Laboratory, University of Reading, to germinate. Twenty-eight days after sowing (DAS), the plants were transplanted into 3L pots containing a peat based potting mix (Clover Peat).

At green bud stage (38 DAS), the plants were moved to two controlled environment chambers (Fitotron, Weiss Technik (UK) Ltd), for a period of one week to adjust to the new environment before the start of the heat experiment. Each cabinet contained 16 plants and was maintained under control conditions of 20 °C/14 °C day/night and a photoperiod of 16/8-hr light/dark.

### 2.2 Heatwave experimental design

Since the flowering stage of winter sown *B. napus* occurs during May in the UK, an analysis of daily weather temperatures during this developmentally crucial period was carried out using the UK Meteorological (Met) Office weather data obtained from the Met office (Met office n.d.) and from WeatherOnline (WeatherOnline n.d.) platforms. The data showed an increase in the frequency of high temperature fluctuations during May, ranging between 26 °C and 28 °C over several days (Supplementary Fig. S1). Given the continuous rise in global temperature, it is expected that future heatwaves would increase in severity and duration (Seneviratne et al., 2012). Based on these data, the current study temperature simulates a heat stress event through a temperature increase to 30°C/24 °C day/night for a period of 6 days and 20 °C/14 °C day/night for control.

Forty-five DAS, the temperature in one cabinet was kept under control conditions (20 °C/14 °C) and denoted as control cabinet, while the second cabinet was set to a gradual increase in temperature and denoted as heat treatment (HT) cabinet. To mimic a heatwave, the temperature in the HT cabinet was increased gradually from 20 °C to 30 °C between 9:00 h and 12:00 h in 3 steps: 20 °C at 9:00 h, 24 °C at 10:00 h, 28 °C at 11:00 h, and 30 °C at 12:00 h. The temperature was held at 30 °C until 22:00 h (Supplementary Fig. S2B), before it was dropped gradually and maintained at 24 °C until 8:00 h of the next day (Supplementary Fig. S2B). This cycle of gradual increase (day) and decrease (night) of temperature was held for five days (Supplementary Fig. S2C). On the 6^th^ day of treatment, the temperature was gradually decreased to 20 °C/14 °C day/night cycle (Supplementary Fig. S2D) and held for a recovery period of 7 days (Supplementary Fig. S2E). During heat treatment, the plants were frequently irrigated to avoid drought.

### 2.3 Sample preparation for transcriptomic analysis

Five biological replicates from each treatment condition were collected at days 0, 1, 2, 6, & 12 of the experiment. Each replicate was made up of three individual plants. Three flower buds were collected per plant (nine buds per replicate). All samples were immediately snap-frozen in liquid nitrogen before they were stored at -80°C. Bud samples were ground into fine powder using pestle and mortar, with the frequent addition of liquid nitrogen to stop enzymatic reactions. The ground material was then stored at -80°C until RNA extraction.

### 2.4 RNA extraction and sequencing

Total RNA from bud samples was extracted using the Spectrum™ Plant Total RNA Kit (Sigma-Aldrich) in accordance with the manufacturer’s protocol and treated with genomic DNAse using DNASE 70-On Column DNase I Digestion set (Sigma-Aldrich) to eliminate DNA contamination. RNA quantification was estimated on both NanoDrop 2000 (Thermo Scientific) and Qubit 2.0 fluorometer (Invitrogen, USA) and its quality was evaluated on 1% (w/v) denaturing formaldehyde agarose gel (MOPS). Samples were shipped over dry-ice to Novogene Europe, where sequencing was performed on the Illumina Novaseq™ 6,000 (PE150) platform.

### 2.5 Differential expression and cluster analysis

Raw sequence reads were assessed for quality using FastQC tools (v0.11.5) (Andrews 2012). The mean sequence lengths were 150 bp and the mean sequence GC content was 43%. The mean quality scores in each base position were higher than 36 and the mean quality scores per sequence were 36. As a result, sequence trimming was not necessary.

STAR software (v2.7.10a) was used to map the clean reads to the *B. napus* cv. Westar v0 reference genome (Song et al., 2020), and the percentage of aligned reads was calculated by using the flagstat command from samtools on each alignment file generated by STAR aligner (Supplementary Figure S3). To estimate transcript abundance, RSEM (RNA-Seq by Expectation-Maximization) software (v1.2.18) was used in three steps to prepare the reference, calculate expression and generate the count matrices. Firstly, rsem-prepare-reference script was used with ‘–gff3’ option to extract reference sequences from the genome and ‘—star’ option to build STAR indices. Next, rsem-calculate-expression script was used with ‘--paired-end’ option to align input reads against the reference transcriptome with STAR and calculated expression values using the alignments. Finally, rsem-generate-data-matrix script was used to generate the count matrices from expression results files.

For differential expression analysis, DESeq2 (v1.41.10) was used in R environment (Love et al., 2014) with default parameters. A pre-filtering step was performed to keep transcripts that have a minimum count of 10 reads in a minimum of four samples. To identify transcripts that were significantly differentially expressed, the two conditions (Heat stress versus Control) were contrasted in pairwise comparisons. Thresholds of |log2(foldchange)| ≥ 0.5 and adjusted P-value < 0.05 (using the Benjamini and Hochberg method) were considered to identify significantly differentially expressed transcripts between the two treatment conditions.

Significant DEGs in each contrast were further analysed using K-mean clustering based on the kmeans() function in R (distance: Euclidean), with log2FC as the input.

### 2.6 Gene Ontology and KEGG pathway enrichment analysis of DEGs

Gene Ontology (GO) and Kyoto Encyclopedia of Genes and Genomes (KEGG) enrichment analyses were conducted on the DEGs using OmicsBox (BioBam Bioinformatics, 2019). The significantly enriched GO terms and biological pathways in HT samples as compared to control at different timepoints were identified to determine heat stress-related functions and pathways. The analysis was carried out using Fisher exact statistical test with FDR adjustment cut-off <0.05. The background dataset consisted of all *B. napus* cv. Westar identifiers present in the assembly’s annotation file (Song et al., 2020).

### 2.7 Validation of RNA-seq results by real-time quantitative PCR (RT-qPCR)

Sucrose Synthase 5 (SS5) and Heat Shock Protein 20 (HSP20) were chosen from the pathways that were involved in the heat stress response to further examine their expression in real time using RT-qPCR. Five timepoints were chosen for each gene to represent the time course of the experiment. SS5 and HSP20 were not analysed at 1 day after treatment (DAT) as they exhibited similar expression profile at 0 & 1 DAT, therefore one timepoint was selected (Supplementary Table S2).

Total RNA used for transcriptomic sequencing was used for cDNA synthesis. First, cDNA was synthesized from 1 µg total RNA by Invitrogen SuperScript IV VILO Master Mix kit according to the manufacturer’s instructions (ThermoFisher Scientific). The RT-qPCR reaction volume was 20 µL consisting of 3 µL cDNA, 10 µL SYBR Premix Ex Taq II (Takara, China), 1 µL (200 nM final concentration) of 4 µM of each primer (forward and reverse, Supplementary Table S3) and 5 µL RNAse-free water. Three technical and three biological replicates for each sample were measured with the following protocol: 50 °C for 2 min, 95 °C for 2 min, followed by 40 cycles of 95 °C for 3 s, 60 °C for 30 s, followed by dissociation curve analysis ramping from 60°C to 95 °C with a ramp rate of 0.3 °C s^-1^ on an ABI StepOne Plus RT-qPCR platform (Applied Biosystems, USA). BnaActin was used as housekeeping gene to normalize the data. The relative expression level of all selected genes at each time point was calculated using the 2^−ΔΔCT^ method of the StepOne software (v2.2.2).

### 2.8 Sample preparation for metabolomic analysis

Five biological replicates from each treatment condition were collected at days 0, 1, 2, 5, 6, 8 & 12 of the heat treatment experiment. Each replicate was made up of three youngest fully expanded leaves collected from three plants. All leaf samples were immediately snap frozen in liquid nitrogen before being stored at -80°C until further analysis. Frozen leaf samples were freeze-dried for five days, then ground into a fine powder using a Precellys 24 lysis & homogenization (Stretton Scientific, Alfreton, UK) and stored at -40 °C in sealed sample bags until extraction and analysis.

### 2.9 Glucosinolates extraction and analysis

GLS extraction was carried out as per the protocol of Bell et al. (2015) as follows: 40 mg of ground leaf powder was heated in a dry block at 75 °C for 2 min. This step was done as a precautionary measure to inactivate as much myrosinase enzyme as possible before extraction (Pasini et al., 2012). Afterwards, 1 mL of preheated (70 % v/v) methanol (70 °C) was added to each sample and placed in a water bath for 20 min at 70 °C. Samples were then centrifuged for 5 min (6000 rpm at 18 °C) to collect the loose material into a pellet. The supernatant was then transferred into fresh labelled Eppendorf tube and stored at -80 °C until further analysis. Before LC-MS analysis, the samples were filtered using 0.25 µm filter discs, diluted in 50% methanol at a 1:4 ratio (dilution factor =5) and spiked with a 100 µL of the internal standard Sinigrin (100 ng mL^-1^). The GLS content were analysed by SCIEX QTRAP 6500+ LC-MS.

For LC separation, a Waters Acquity BEH C18 column (particle size 1.7µ, 2.1 x 50 mm) with a Security VanGuard system from Waters (UK) was used. The mobile phase consisted of water with 0.1 % formic acid (A) and methanol with 0.1 % formic acid (B). The gradient started at 5 % B and was raised to 90 % B in 3.5 min, held at 90 % B for 0.5 min and re-equilibrated at 5 % B for 1 min. The total time of the gradient program was 5 min. The flow rate was 0.4 mL min^-1^ with a column temperature of 60 °C. A 1 µL aliquot of sample was injected for analysis.

The LC system was interfaced with a SCIEX QTRAP 6500+ mass spectrometer equipped with an IonDrive Turbo V ion source. Multiquant software was used to control sample acquisition and data analysis. The QTRAP 6500+ mass spectrometer was tuned and calibrated according to the manufacturer’s recommendations. For quantification, an external standard calibration curve was prepared using a series of standard samples containing the following GLS: Gluconapin (GNA), Progoitrin (PRO) & Glucobrassicin (GBS), with concentrations: 1000 ng mL^-1^, 500 ng mL^-1^, 250 ng mL^-1^, 100 ng mL^-1^, 50 ng mL^-1^, 10 ng mL^-1^, and 1 ng mL^-1^. Two-way mixed ANOVA in R was performed at 0.05 significance level to calculate the significance variation in concentration under heat treatment as compared to control.

### 2.10 Sugars content extraction and analysis

The extraction of soluble sugars (Glucose, Fructose, and Sucrose) was performed as follows: 50 mg of ground leaf powder was extracted with 1 mL of 62.5 % methanol: 37.5% HPLC-grade water (v/v) at 55 °C over a period of 15 min in a shaking water bath and vortexed every 5 min. Afterwards, the samples were allowed to cool for 2 min and then centrifuged at 13000 rpm for 10 min and filtered using 0.25 µm filter discs. The supernatant was then transferred to a clean labelled Eppendorf tube and stored at -80 °C until further analysis. Before HPLC analysis, the samples were diluted in HPLC-grade water in a 1:2 ratio (dilution factor = 3). Chromatographic separation of sugars content was performed using HPLC (Agilent 1260, Infinity Series) equipped with an Evaporative Light Scattering Detector (ELSD) system and an Asahipak NH2P-50 4E column (250 × 4.6 mm; Shodex, Tokyo, Japan) in an isocratic elution mode. The mobile phase consisted of 75 % acetonitrile: 25 % water at a flow rate of 1 mL min^-1^, a column temperature of 40 °C and an injection volume of 20 µL. The analysis had a total run time of 25 min. For quantification, an external standard calibration curve was prepared using a series of standard samples containing Glucose, Fructose and Sucrose with concentrations: 0.025 mg mL^-1^, 0.05 mg mL^-1^, 0.1 mg mL^-1^, 0.5 mg mL^-1^ and 1 mg mL^-1^.

### 2.11 Near-Infra Red Spectroscopy (NIRS) scanning

Following the treatments, the plants were allowed to complete their lifecycle and total aboveground biomass and seed yields were collected from individual plants. Seed samples were scanned with DA 7250 NIR analyzer (PerkinElmer, Beaconsfield, UK). The seed samples were scanned for moisture, oil and protein content as well as for fatty acid composition of the oil.

### 2.12 Statistical analysis

Two-way mixed ANOVA was performed at 0.05 significance level to calculate the statistically significant differences between the means of sugars and glucosinolate concentration under heat treatment as compared to control. In addition, NIRS data was analysed for the investigated components using a t-test. All statistical analysis was performed using R.

## 3. RESULTS

### 3.1 Prolonged heat exposure reduced yield and altered B. napus oil composition

Heat treatment significantly altered plant growth and appearance, with visible physiological changes on the flowers and buds (Fig. 1). At 5 DAT, flowers displayed smaller and paler petals compared to plants growing under control conditions. At 3 days of recovery (DOR), many buds suffered from abscission, appeared brown and shrunken in size on plants subjected to the heat treatment. At maturity, heat-treated plants exhibited less total biomass and total seed weight relative to control plants (Supplementary Fig. S4). In terms of oil quality, heat treatment significantly decreased Linoleic Acid, Linolenic Acid and Palmitic Acid concentration, but increased Oleic Acid and Stearic Acid concentration in seeds compared to control plants (Supplementary Fig. S5)

**Figure 1.**
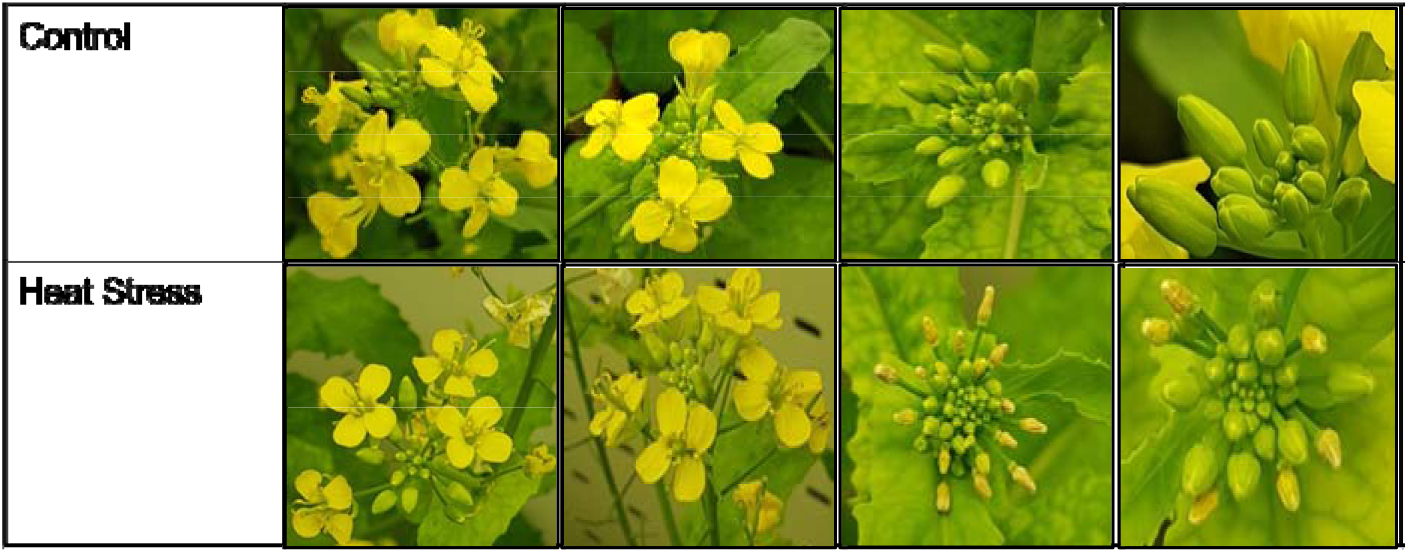
B. napus flowers and buds at 5 DAT & 3 DOR respectively.

### 3.2 Differential gene expression identifies distinct patterns of expression in B. napus under heat stress and recovery

Libraries were constructed and sequenced on the Illumina NovaSeq platform. An average of 47 million reads were generated per sample after low quality reads were filtered out (Supplementary Table S1). Following this cleaning step, around 90%-95.49% total reads per sample were mapped to the B. napus cv. Westar v0 reference genome (Supplementary Fig. S3) (Song et al., 2020). Principle component analysis (PCA) was used to analyse the samples which drove the group separation (Supplementary Fig. S7). PC1 and PC2 captured 48.9% and 8.2% of the total variance of the samples respectively. The analysis showed that, both control and treatment samples clustered near each other at 7 DOR which demonstrated that heat treated plants might have started to return to normal after 7 days of recovery.

To understand the effect of heat stress on gene expression in *B. napus*, differentially expressed genes (DEGs) were identified in a pairwise comparison between heat treatment and control conditions at five timepoints during and after heat treatment. A total of 1892, 3253, 4553, 4165 and 1713 genes were differentially expressed at days 0, 1, 2, 6 and 12 respectively (Supplementary Fig. S8). Analysis of the top differentially expressed genes after 24 hrs of heat treatment revealed that eight transcripts with logFC ranging between 4.4-10.8 mapped to small heat shock proteins (sHSP20), in addition to transcripts mapped to other heat shock factors such as Elongation Factor 1-beta 1-like and Chaperone Protein ClpB1 (Supplementary Table S7). On the other hand, top down regulated genes featured transcripts mapped to sugars transporters, nitrogen transport and storage, cell wall modification and methylation (Supplementary Table S8).

To identify trends in the expression of genes across the time course of the experiment, K-means clustering was applied to 9933 DE genes with logFC ≥ 0.5. The algorithm randomly assigned each gene into one of the clusters based on the Euclidean distance between the gene and the cluster mean. Six distinctive clusters based on expression changes across the five timepoints were identified (Fig. 2). Interestingly, clusters 1 & 3 (475 & 607 genes), showed opposite expression patterns towards the end of the heat treatment and the start of the recovery phase. In cluster 1, the expression profile of genes decreased at 1 DOR (HS6vsC6), before returning to normal levels at 7 DOR (HS12vsC12), while in cluster 3, a sharp increase in expression was observed at 1 DOR (HS6vsC6), which returned to normal levels at 7 DOR (HS12vsC12).

**Figure 2.**
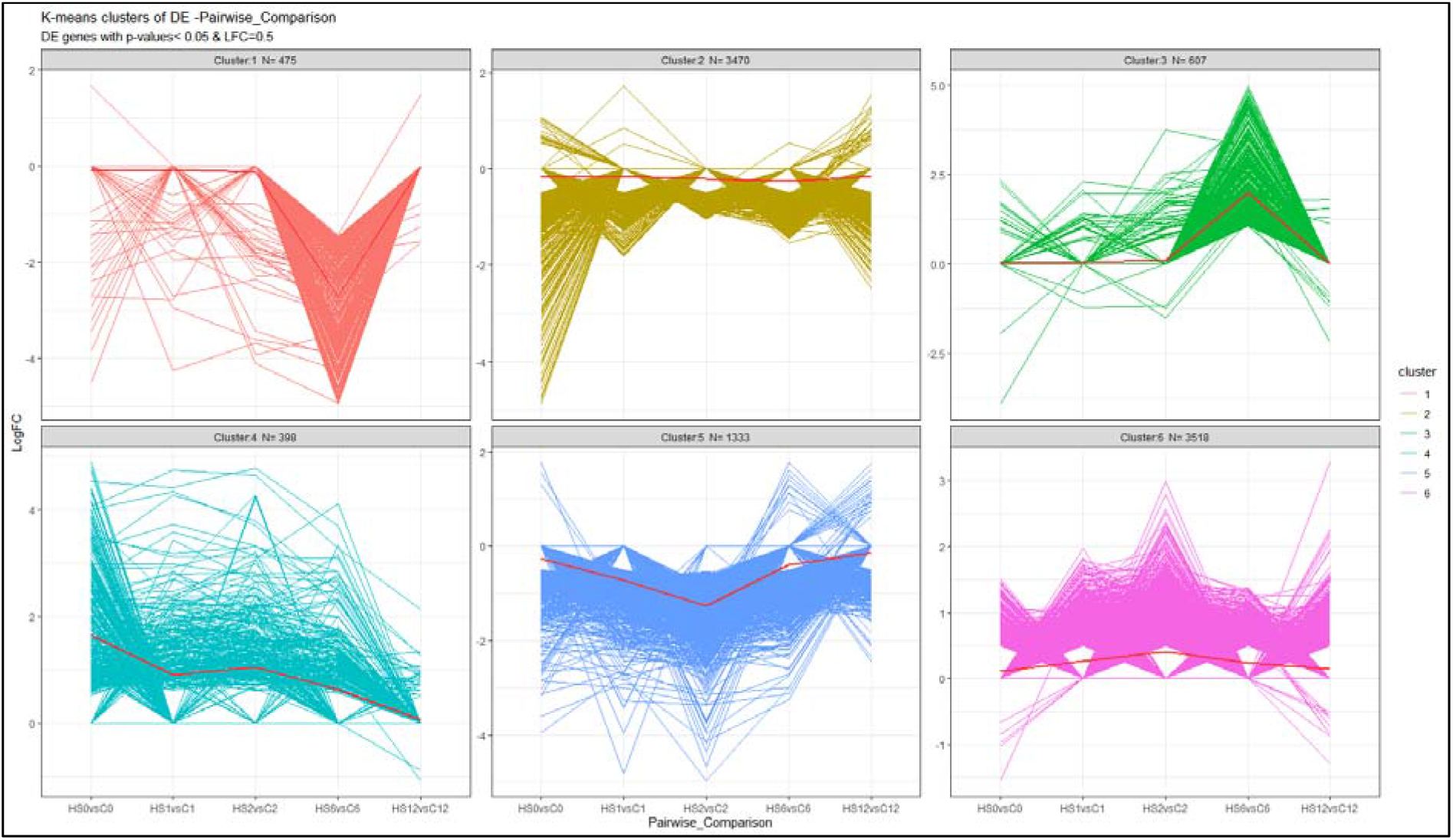
K-means cluster analysis of differential expressed genes in B. napus during and after heat treatment. K-means clustering was applied to 9933 DE genes with logFC ≥ 0.5 using the “kmeans” function in R package “stats” (v. 3.2.2), where k represents a pre-specified number of clusters. Six groups of genes were classified, with N the number of genes in each cluster, and the solid red lines in each cluster indicate mean changes in DEG expression.

In clusters 2 & 5 (3470 & 1333 genes), genes were down regulated along the course of the experiment, but more pronounced during the heat treatment phase especially in cluster 5 which exhibited a greater decease in expression. Both clusters 4 & 6 (398 & 3518 genes) showed increased expression during the treatment before it started to decline towards the end, reaching normal levels at 7 DOR (HS12vsC12) (Fig. 2).

### 3.3 Validation of RNA-seq data by quantitative real-time PCR

To verify the reliability of RNA sequencing results, two genes with diverse expression profiles, including up regulated or down regulated at different timepoints of the experiment, were selected for real-time qPCR to measure expression levels. As a result, HSP20 (Heat Shock Protein 20) and SS5 (Sucrose Synthase 5), exhibited similar expression profiles between RT-qPCR and RNA-seq data (Supplementary Fig. S9 & Table S4), with correlation coefficients of r=0.86, and r=0.68 respectively.

### 3.4 Functional Annotation of DEGs identifies key pathways responding to heat treatment and recovery

Gene ontology (GO) enrichment analysis of the DEGs was performed to identify enriched GO terms. Terms such as “binding”, “catalytic activity”, “metabolic process”, “hydrolase activity” and “ion binding” were among the predominant enriched terms at all timepoints with more transcripts mapped to these terms during heat treatment than during recovery (Supplementary Figs. S10-S14). Oxidoreductase, Hydrolase and Transferase were among the top identified enzymes (Supplementary Fig. S15). These results suggest that high temperature activates several metabolic processes through the expression of numerous enzymes involved in alleviating the impact of heat stress not only during the stress period but also during recovery.

Based on KEGG (Kyoto Encyclopedia of Genes and Genomes) enrichment analysis (Kanehisa & Goto, 2000), pathways in the respiratory metabolism namely Glycolysis and Pentose Phosphate Pathway were enriched in all heat treatment comparisons at all timepoints, while Citrate Cycle and Oxidative Phosphorylation were enriched at 2 DAT and 1 DOR respectively (Supplementary Figs. S16-S19). Other related pathways were also enriched along the course of the treatment and during recovery period, and these include one Carbon Pool by Folate (enriched at 1 & 2 DAT and 1 DOR), Carbon Fixation in Photosynthetic Organisms, Cysteine and Methionine metabolism (enriched at 1 & 2 DAT), Glycerolipid and Tryptophan metabolism (enriched at 2 DAT), Fatty Acid Biosynthesis and Pyruvate metabolism (enriched at 1 DOR) and Fructose and Mannose metabolism (enriched at 7 DOR). In the present study, most of the genes involved in these pathways were responsive to high temperature stress, with their expression declining with the onset of heat treatment, especially after 48 hours (Supplementary Figs. S20-S27).

#### 3.4.1 Heat treatment down regulated the transcript levels of most aliphatic and indolic GLSs synthetic genes

Analysis of the aliphatic GLS synthetic and regulatory genes showed that heat treatment had a significant effect on the expression of 18 genes at 2 DAT (Fig. 3). Thirteen genes encoding different GLS synthetic genes including different UDP Glycosyltransferases and Glutathione S-transferases were down regulated while 5 genes encoding GLS related transcription factors were up regulated. At 1 DOR which marks 24 hours of recovery, nine genes that were not differentially expressed under heat treatment were found up regulated. These genes include different Cytochrome P450 genes, transcription factors, Branched-Chain Aminotransferase 4.2 and Isopropylmalate Dehydrogenase 2.

Analysis of the indolic GLS synthetic and regulatory genes showed that heat treatment altered the expression of 16 and 23 out of 44 genes at 1 & 2 DAT respectively (Fig. 4). These genes include Sulfotransferases, Cytochrome P450 genes, UDP Glycosyltransferases, Glutathione S-transferases and transcription factors. After one week recovery (7 DOR), most of the differentially expressed genes identified during heat treatment were no longer differentially expressed. This is similar to what was seen with the aliphatic GLS encoding genes. Interestingly, one transcript encoding UDP Glycosyltransferase 74E2-like remained down regulated after one week of recovery.

**Figure 3.**
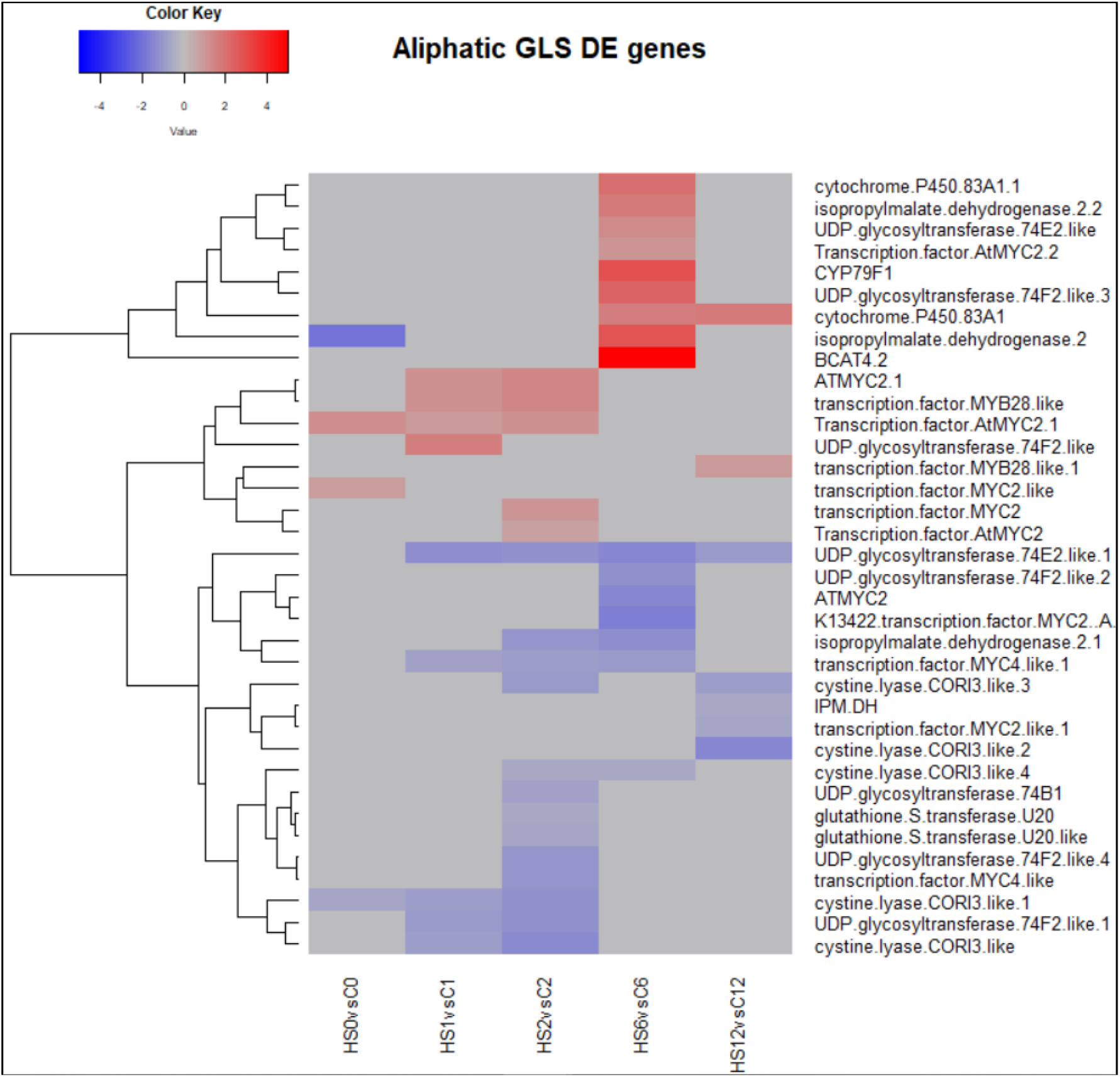
Heatmap showing differential expression of genes involved in the regulation or biosynthesis of aliphatic glucosinolates (GLS) in Brassica napus. Thirteen genes encoding different GLS synthetic genes including different UDP Glycosyltransferases and Glutathione S-transferases were down regulated while 5 genes encoding GLS related transcription factors were up regulated at 2 DAT.

**Figure 4.**
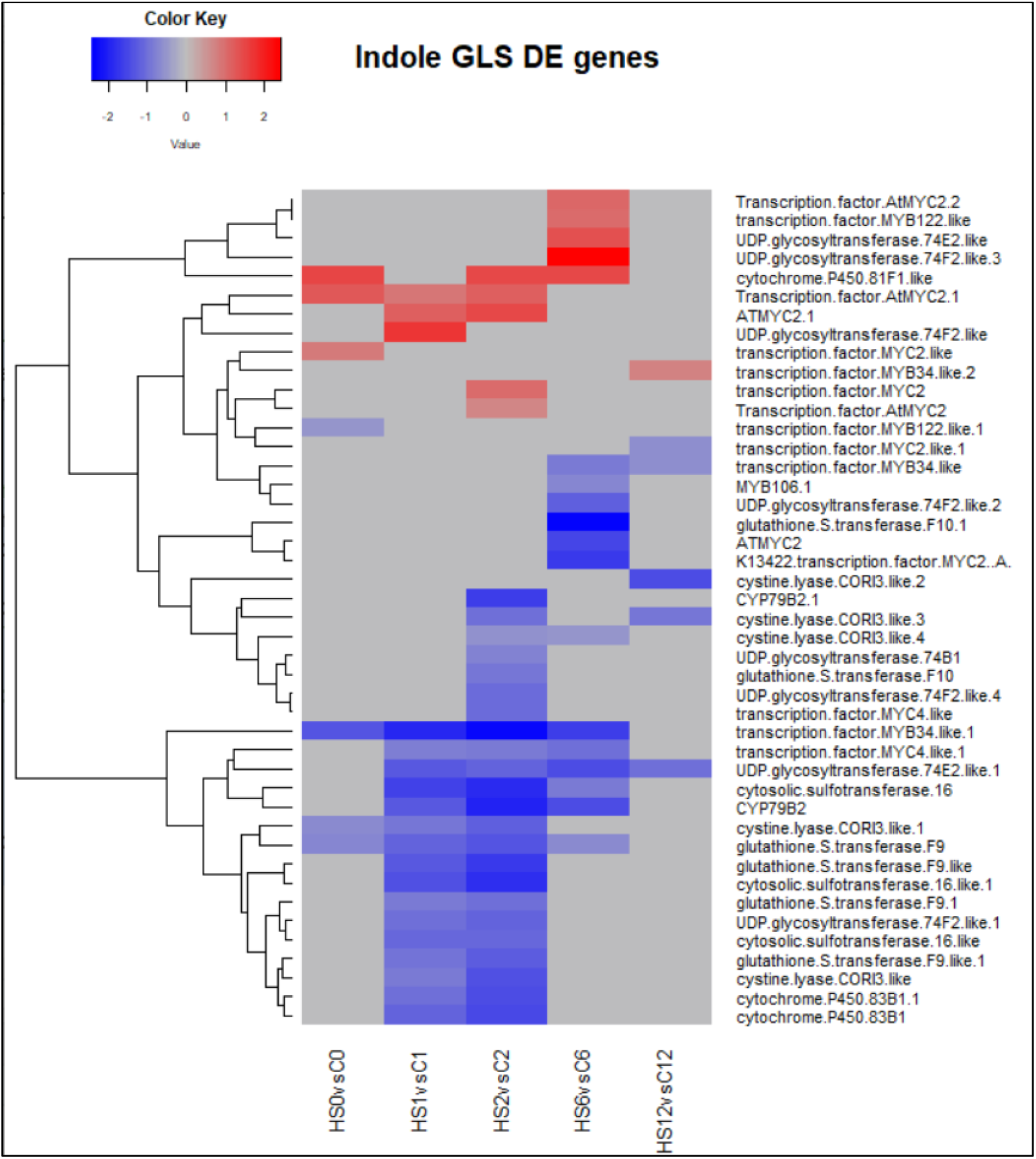
Heatmap showing differential expression of genes involved in the regulation or biosynthesis of indolic glucosinolates (GLS) in Brassica napus. Sixteen and twenty-three out of 44 genes encoding different Sulfotransferases, Cytochrome P450 genes, UDP Glycosyltransferases, Glutathione S-transferases and transcription factors were down regulated at 1 & 2 DAT respectively.

#### 3.4.2 Heat treatment down regulated the transcript levels of most Sulphur assimilation and transport genes

Analysis of the genes encoding Sulphur assimilation and transport during heat treatment showed that 20 transcripts were down regulated in response to heat at either 1 or 2 DAT or both (Fig. 5). One transcript encoding Sulphate transporter was up regulated at 1 & 2 DAT, while two transcripts encoding Adenylyl Sulphate Kinase was up regulated at 2 DAT only. Interestingly, after the end of heat treatment, only 5 transcripts remained down regulated at 1 DOR, and 2 transcripts encoding Adenylyl Sulphate Kinase and ATP sulfurylase were up regulated at 7 DOR.

**Figure 5.**
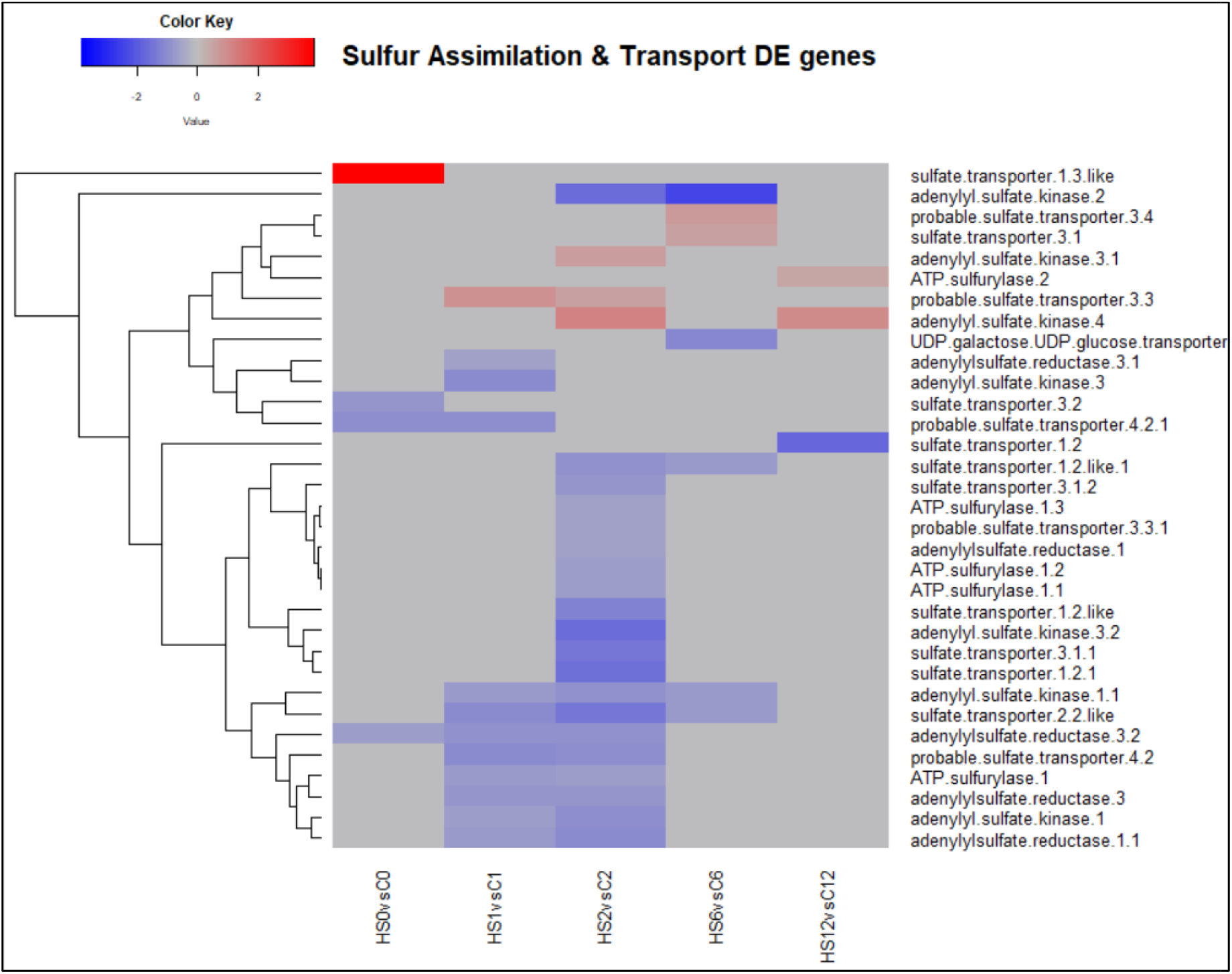
Heatmap showing differential expression of genes involved in the assimilation and transport of Sulphur in Brassica napus. Twenty transcripts were down regulated in response to heat at either 1 or 2 DAT or both.

#### 3.4.3 Metabolic analysis revealed altered leaf concentration of aliphatic and indolic GLS in response to heat treatment

Given the changes observed in the abundance of transcripts associated with GLS metabolism and Sulphur assimilation and transport, the concentration of two aliphatic GLS Progoitrin (PRO) and Gluconapin (GNA), and one indole GLS Glucobrassicin (GBS) were quantified using LC-MS. Their role in response to different abiotic stress factors has been previously reported (Ljubej et al, 2021; Jasper et al, 2020). Two-way mixed ANOVA pairwise comparisons showed a gradual increase in both PRO and GNA concentration in response to heat (Figs. 6 & 7). This increase was significant at 2 DAT. After 5 days of the heat treatment, the concentration of GLS started to decline but remained higher in heat treated plants than in the leaves of control plants. In contrast, the concentration GBS decreased in response to heat treatment (Fig. 8). During recovery, the concentration of GBS continued to decrease with the difference being significant at 1 & 3 DOR. After one week of recovery, GBS concentration in heat treated plants remained lower than that of the control.

**Figure 6.**
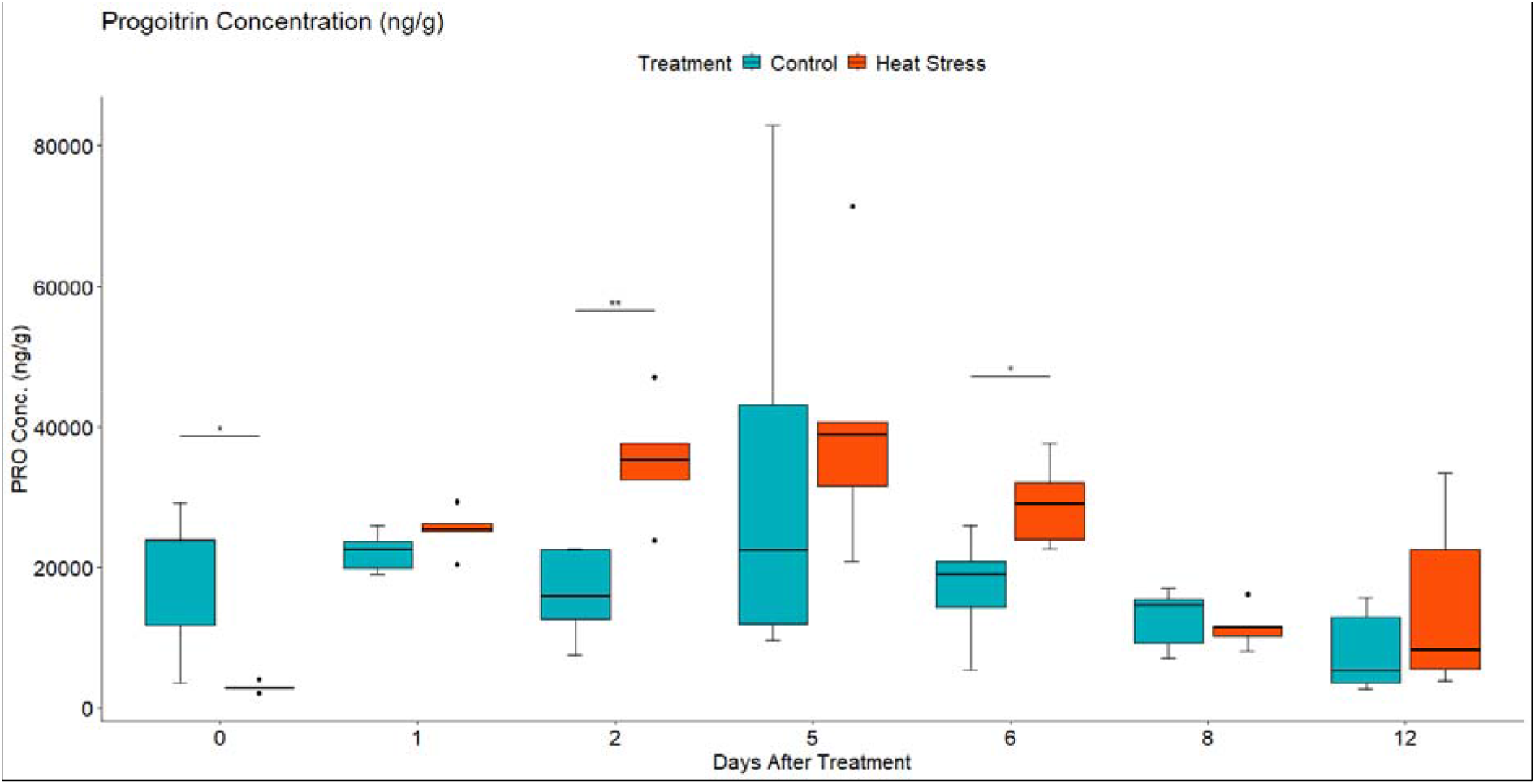
Leaf Progroitin (PRO) concentration in ng/g of dry weight (DW) in Brassica napus plants subjected to heat treatment (30 °C/24 °C day/night) at 0-5 DAT (red) or maintained under control temperatures (20 °C/14 °C day/night) at all timepoints (blue). GLS concentration were quantified using LC-MS. Data were analysed using two-way mixed ANOVA pairwise comparisons. Each box plot represents data from 5 replicates, showing the median (horizontal bar), 25^th^ and 75^th^ percentiles (the box) and the min and max values (whiskers), with outliers as dots. Significant differences between heat treated and control plants at each timepoint are denoted by (*) for p-value ≤ 0.05 or (**) for p-value ≤ 0.01. Heat treatment significantly increased leaf concentration of aliphatic GLS Progoitrin.

**Figure 7.**
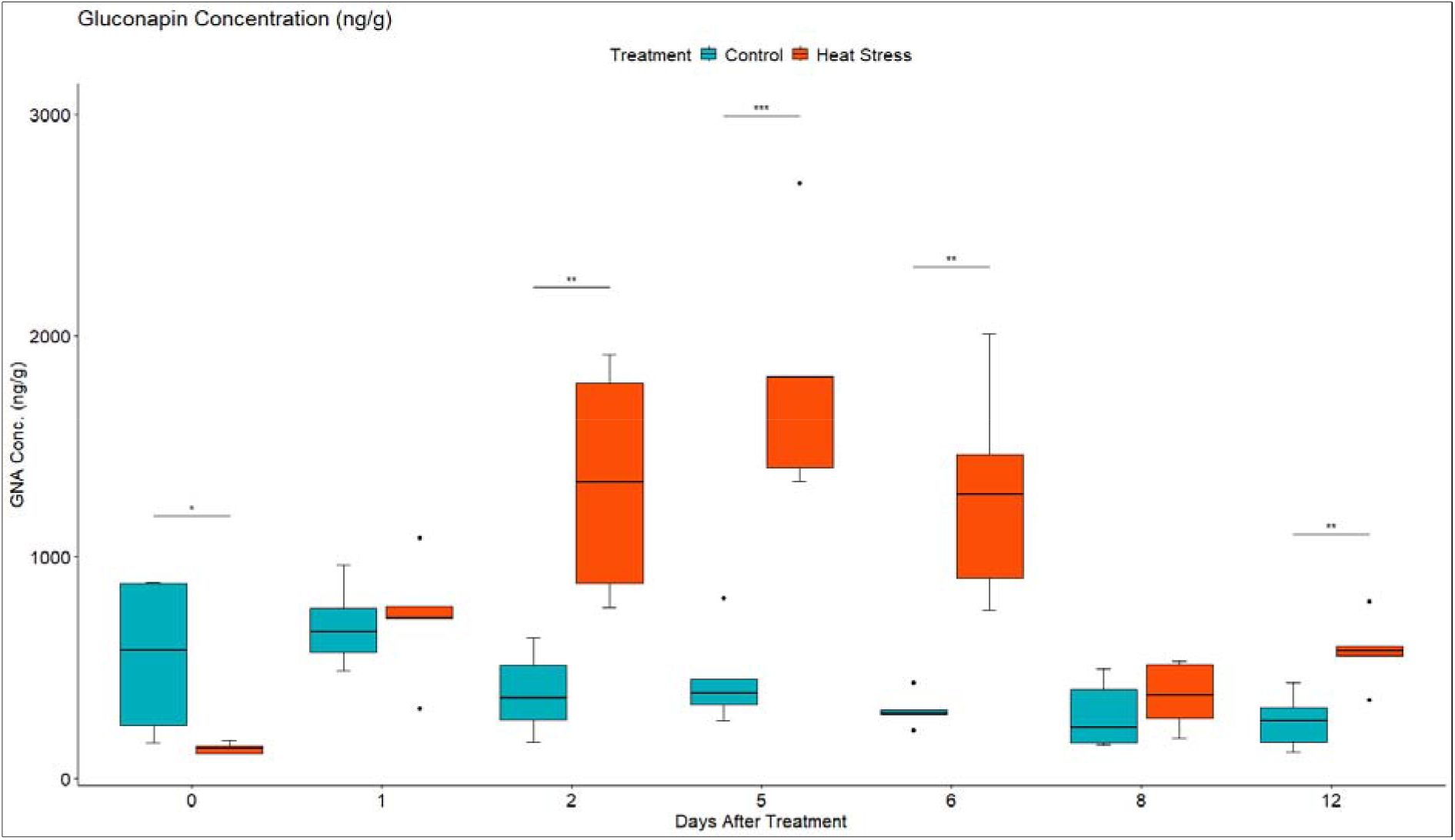
Leaf Gluconapin (GNA) concentration in ng/g of dry weight (DW) in Brassica napus plants subjected to heat treatment (30 °C/24 °C day/night) at 0-5 DAT (red) or maintained under control temperatures (20 °C/14 °C day/night) at all timepoints (blue). GLS concentration were quantified using LC-MS. Data were analysed using two-way mixed ANOVA pairwise comparisons. Each box plot represents data from 5 replicates, showing the median (horizontal bar), 25^th^ and 75^th^ percentiles (the box) and the min and max values (whiskers), with outliers as dots. Significant differences between heat treated and control plants at each timepoint are denoted by (*) for p-value ≤ 0.05 or (**) for p-value ≤ 0.01 or (***) for p-value ≤ 0.001. Heat treatment significantly increased leaf concentration of aliphatic GLS Gluconapin.

**Figure 8.**
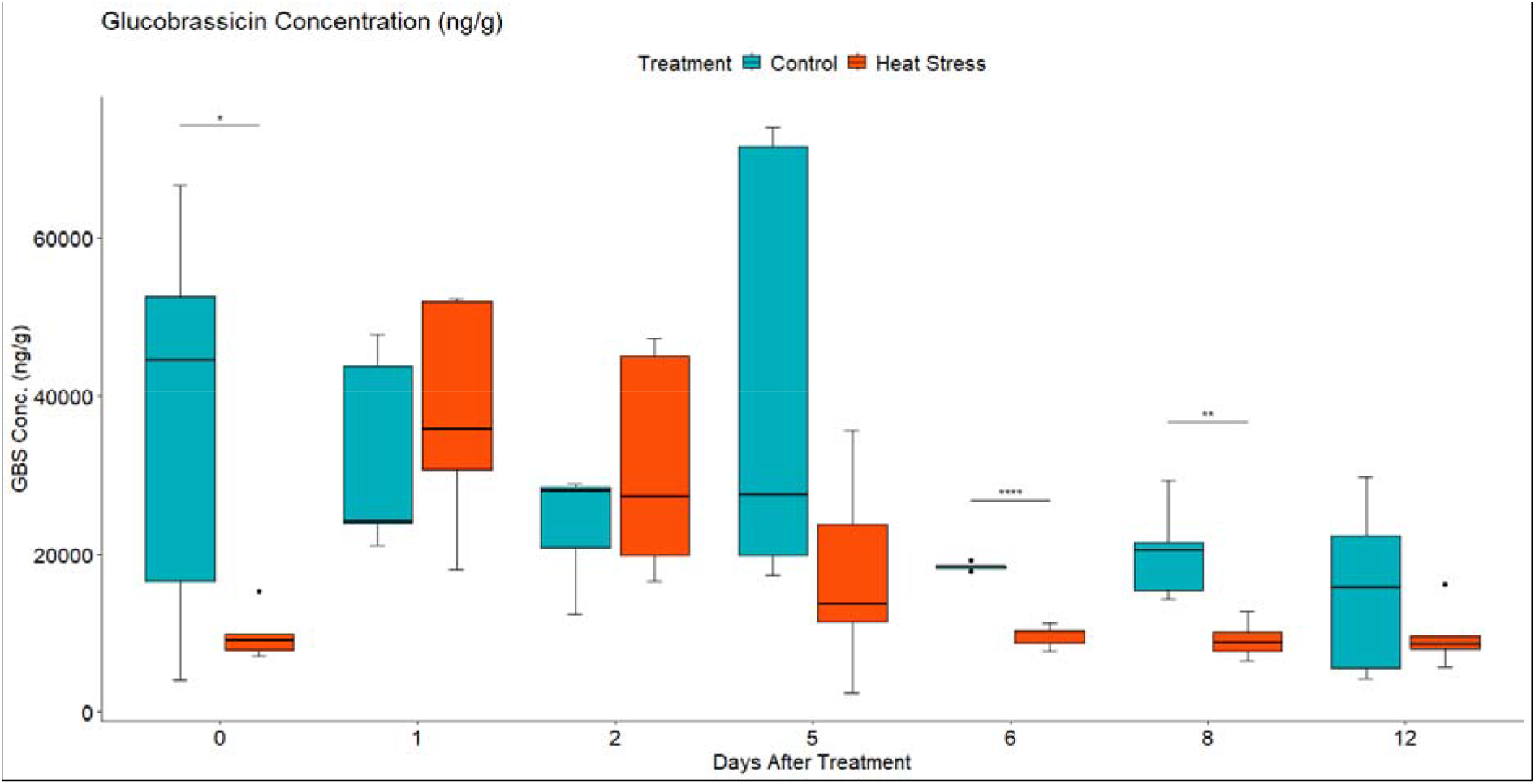
Leaf Glucobrassicin (GBS) concentration in ng/g of dry weight (DW) in Brassica napus plants subjected to heat treatment (30 °C/24 °C day/night) at 0-5 DAT (red) or maintained under control temperatures (20 °C/14 °C day/night) at all timepoints (blue). GLS concentration were quantified using LC-MS. Data were analysed using two-way mixed ANOVA pairwise comparisons. Each box plot represents data from 5 replicates, showing the median (horizontal bar), 25 and 75 percentiles (the box) and the min and max values (whiskers), with outliers as dots. Significant differences between heat treated and control plants at each timepoint are denoted by (*) for p-value ≤ 0.05 or (**) for p-value ≤ 0.01 or (****) for p-value ≤ 0.0001. Heat treatment significantly decreased leaf concentration of the indolic GLS Glucobrassicin.

#### 3.4.4 Heat treatment down regulated the transcript levels of most sugars transporter genes

To investigate the effect of heat stress on sugars transport and metabolism, analyses of genes encoding the SWEET and ERD6-like sugars transporters were undertaken. Compared to the control plants, heat stress significantly altered the expression of 21 SWEET genes with logFC ranged between -5 and 5 (Fig. 9). These genes belonged to the phylogenetic clades II and III. Members of these clades predominantly transport Hexose and Sucrose respectively. Nine of these transcripts were down regulated at 2 DAT, while two were up regulated. Interestingly, out of the 13 genes that were DE at 1 DOR, five up regulated genes were not DE during heat stress (were not affected by heat treatment) and became up regulated after the removal of stress (Fig. 9).

**Figure 9.**
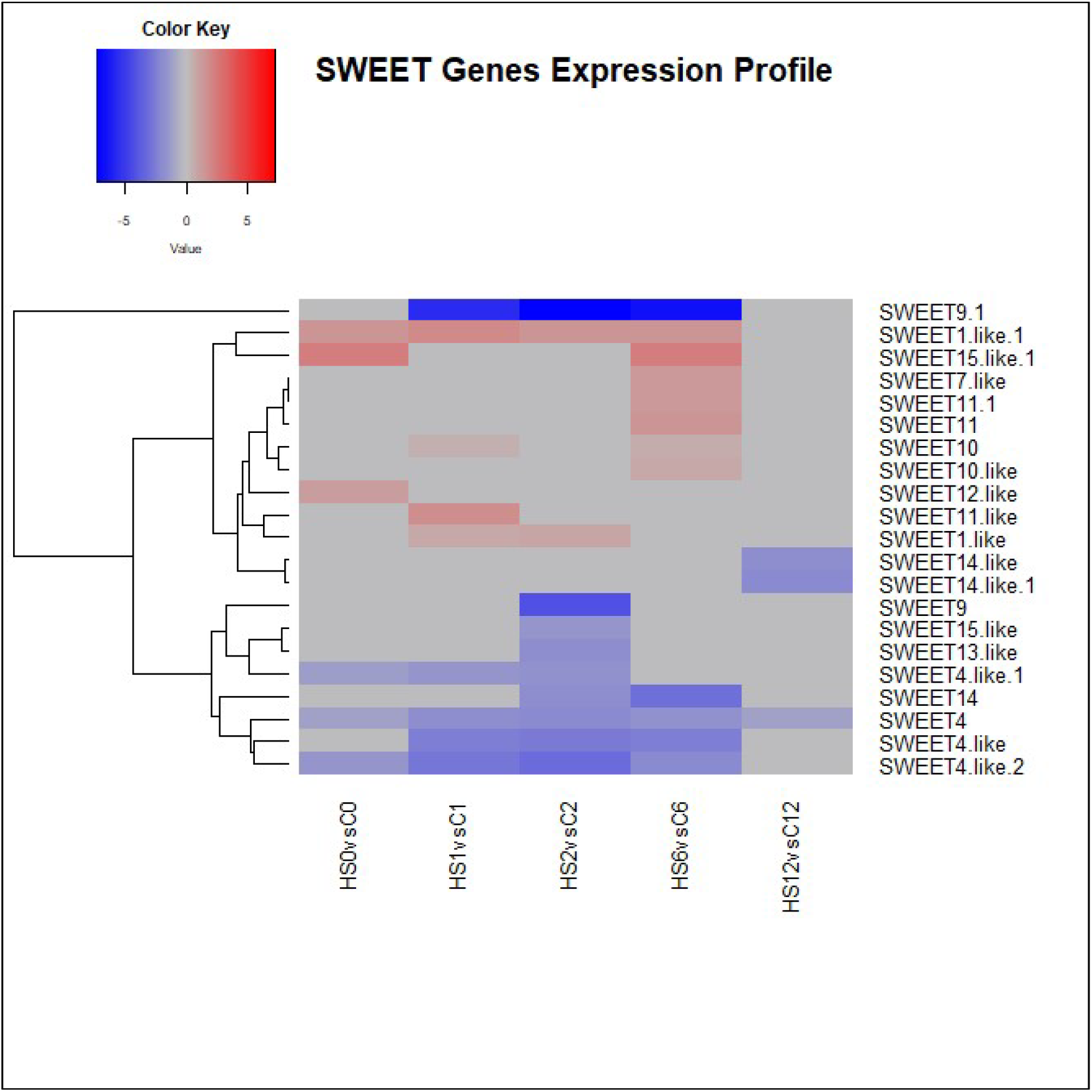
Heatmap showing differential expression of SWEET genes involved in sugars transport in Brassica napus. Heat stress significantly altered the expression of 21 SWEET genes with logFC ranged between -5 and 5.

Analysis of genes encoding the tonoplastic Glucose symporters showed that 11 transcripts mapped to different ERD6-like genes (Fig. 10). At 2 DAT, five were down regulated and two were up regulated.

**Figure 10.**
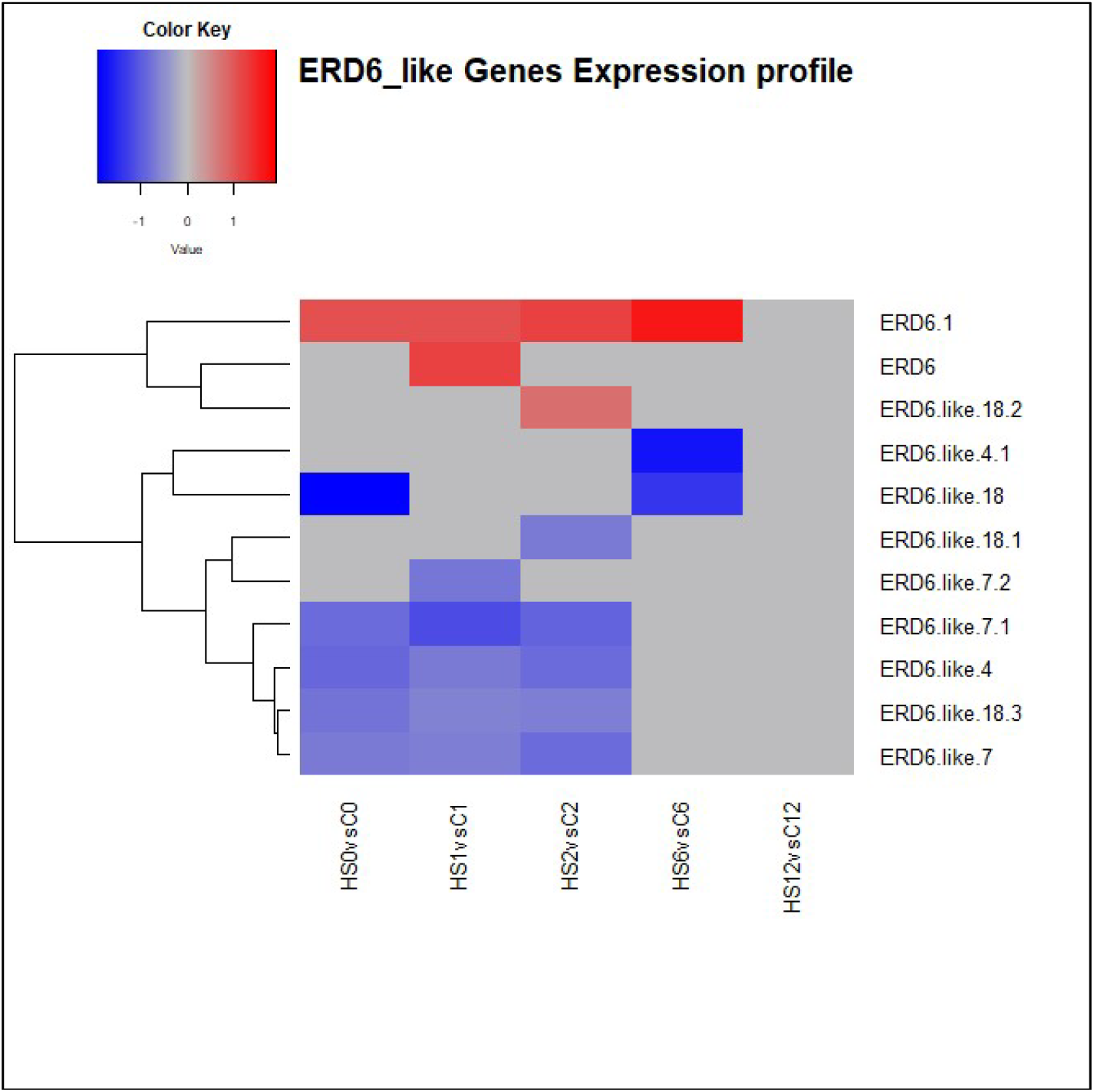
Heatmap showing differential expression of ERD-6 like genes involved in sugars transport in Brassica napus. At 2 DAT, five transcripts were down regulated and two transcripts were up regulated.

In addition, one transcript mapping to a Facilitated Glucose transporter member 8-like showed a LogFC of -9.2 at 1 DAT, and transcripts encoding Sucrose transporters (SUCs) were found up regulated at 1 DAT & 1 DOR (Supplementary Fig. S28). In addition to their significant role in phloem loading and unloading, these transporters are also involved in sugars influx into the cytosol.

Furthermore, the present results showed that the expression of transcripts mapping to the Sucrose synthesizing enzyme, Sucrose Phosphate Synthase (SPS) increased during heat treatment (Supplementary Table S5), while the expression of Sucrose catalysing enzymes, namely Sucrose Synthase and Cell Wall Invertase significantly decreased (Supplementary Table S6).

#### 3.4.5 Metabolic analysis revealed an increase in leaf concentration of sugars in response to heat treatment

To investigate the effect of heat stress on the sugars concentration in the leaves of *B. napus*, the concentration of Fructose, Glucose and Sucrose were quantified using High Performance Liquid Chromatography. Under control growth conditions, leaf Fructose, Glucose and Sucrose concentration increased slowly as the plants grew (Figs. 11-13). In contrast, in the heat-treated plants, the leaf concentration of individual sugars increased. Two-way mixed ANOVA pairwise comparisons showed that the change in sugars concentration was significant during both heat treatment and recovery for Glucose and Fructose.

After the removal of heat stress, the leaf concentration of Sucrose in the treated plants decreased gradually until it reached a lower concentration than that of the control plants at 3 DOR. In contrast, Fructose and Glucose concentration showed a slight decrease initially before they increased again significantly at 3 DOR. After 7 days of recovery, the content of Fructose, Glucose and Sucrose dropped substantially, reaching lower concentration than that of the control plants.

**Figure 11.**
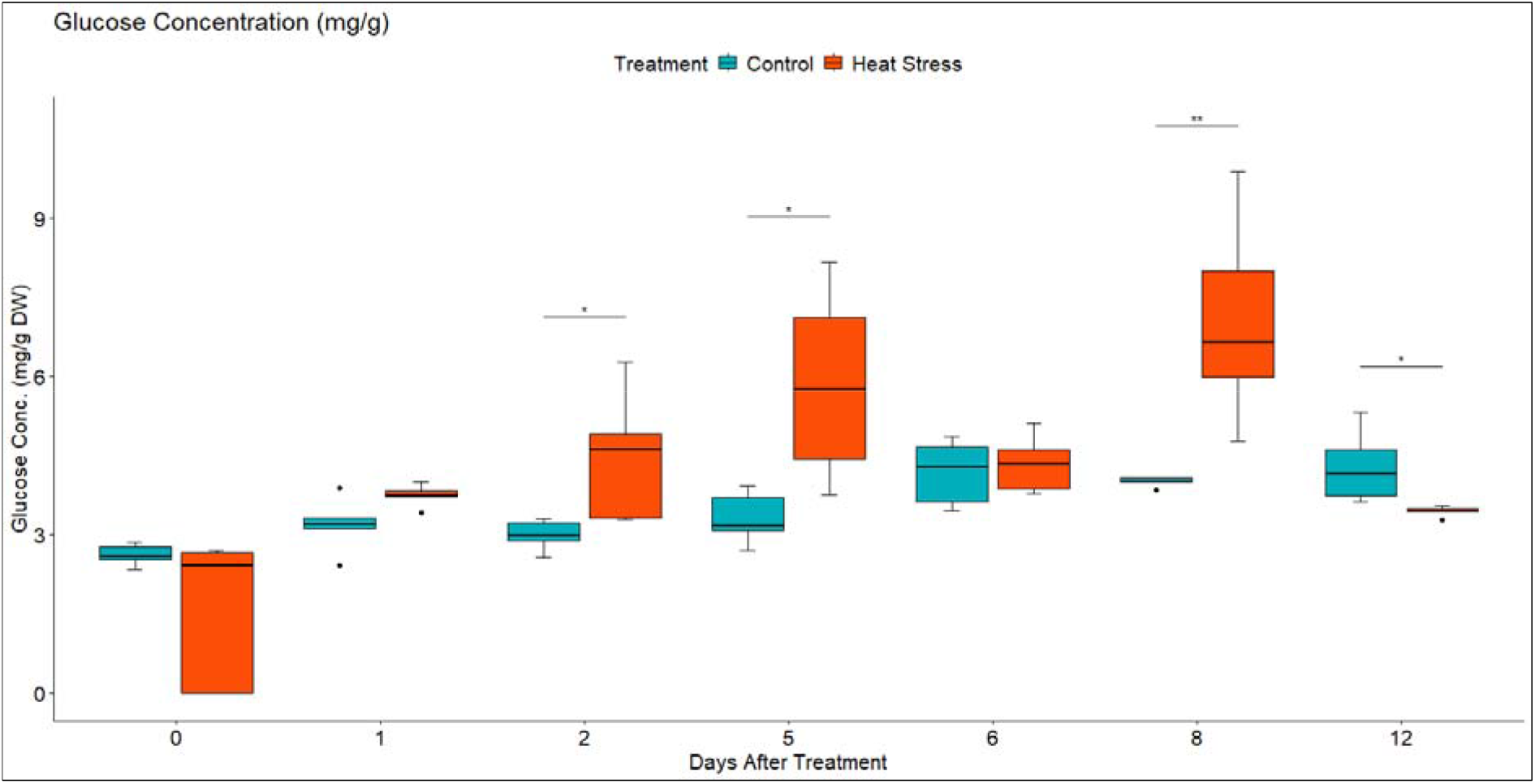
Leaf Glucose concentration in mg/g of dry weight (DW) in Brassica napus plants subjected to heat treatment (30 °C/24 °C day/night) at 0-5 DAT (red) or maintained under control temperatures (20 °C/14 °C day/night) at all timepoints (blue). Concentration was quantified using HPLC. Data were analysed using two-way mixed ANOVA pairwise comparisons. Each box plot represents data from 5 replicates, showing the median (horizontal bar), 25^th^ and 75^th^ percentiles (the box) and the min and max values (whiskers), with outliers as dots. Significant differences between heat treated and control plants at each timepoint are denoted by (*) for p-value ≤ 0.05 or (**) for p-value ≤ 0.01. Heat treatment significantly increased leaf concentration of Glucose.

**Figure 12.**
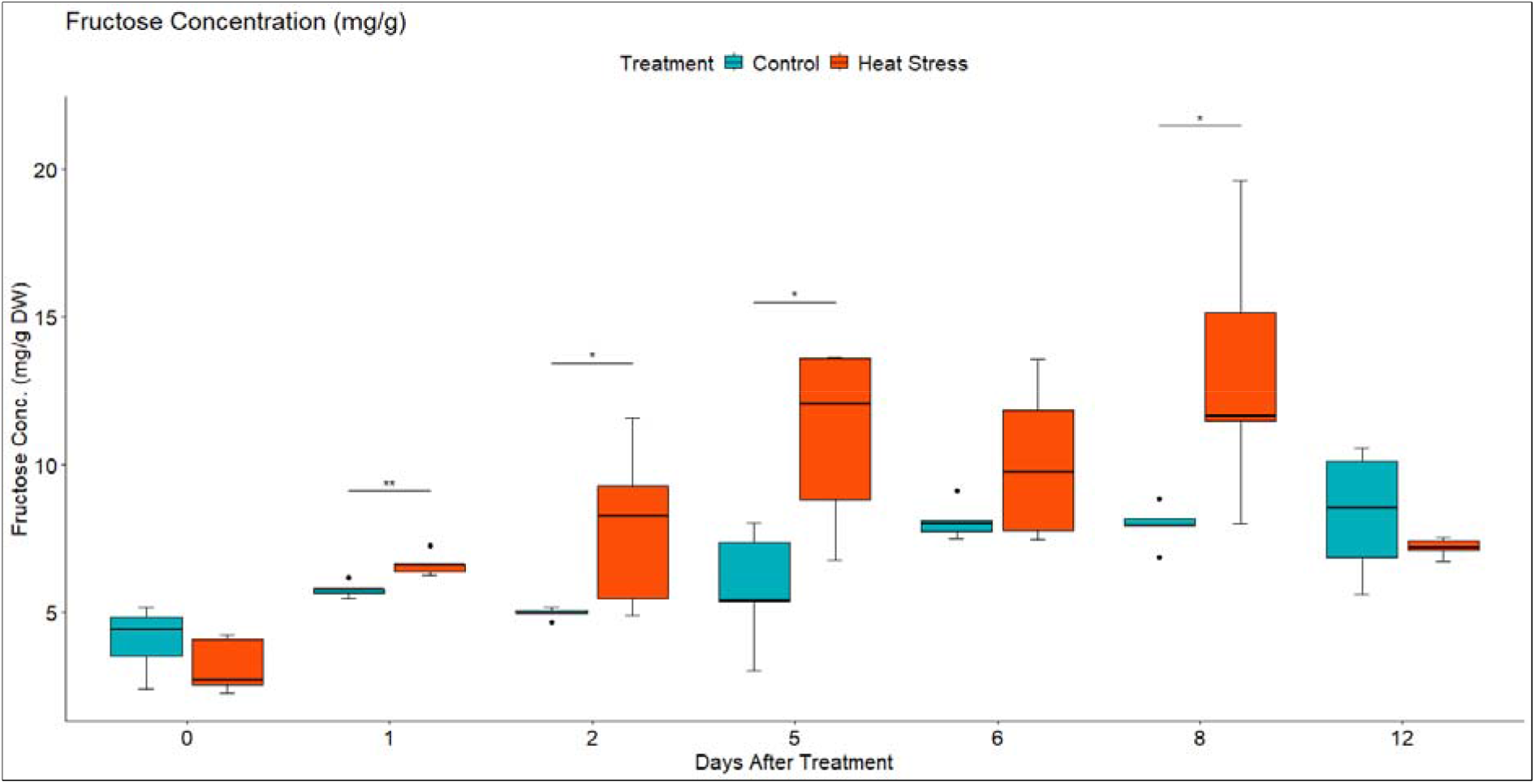
Leaf Fructose concentration in mg/g of dry weight (DW) in Brassica napus plants subjected to heat treatment (30 °C/24 °C day/night) at 0-5 DAT (red) or maintained under control temperatures (20 °C/14 °C day/night) at all timepoints (blue). Concentration was quantified using HPLC. Data were analysed using two-way mixed ANOVA pairwise comparisons. Each box plot represents data from 5 replicates, showing the median (horizontal bar), 25 and 75 percentiles (the box) and the min and max values (whiskers), with outliers as dots. Significant differences between heat treated and control plants at each timepoint are denoted by (*) for p-value ≤ 0.05 or (**) for p-value ≤ 0.01. Heat treatment significantly increased leaf concentration of Fructose.

**Figure 13.**
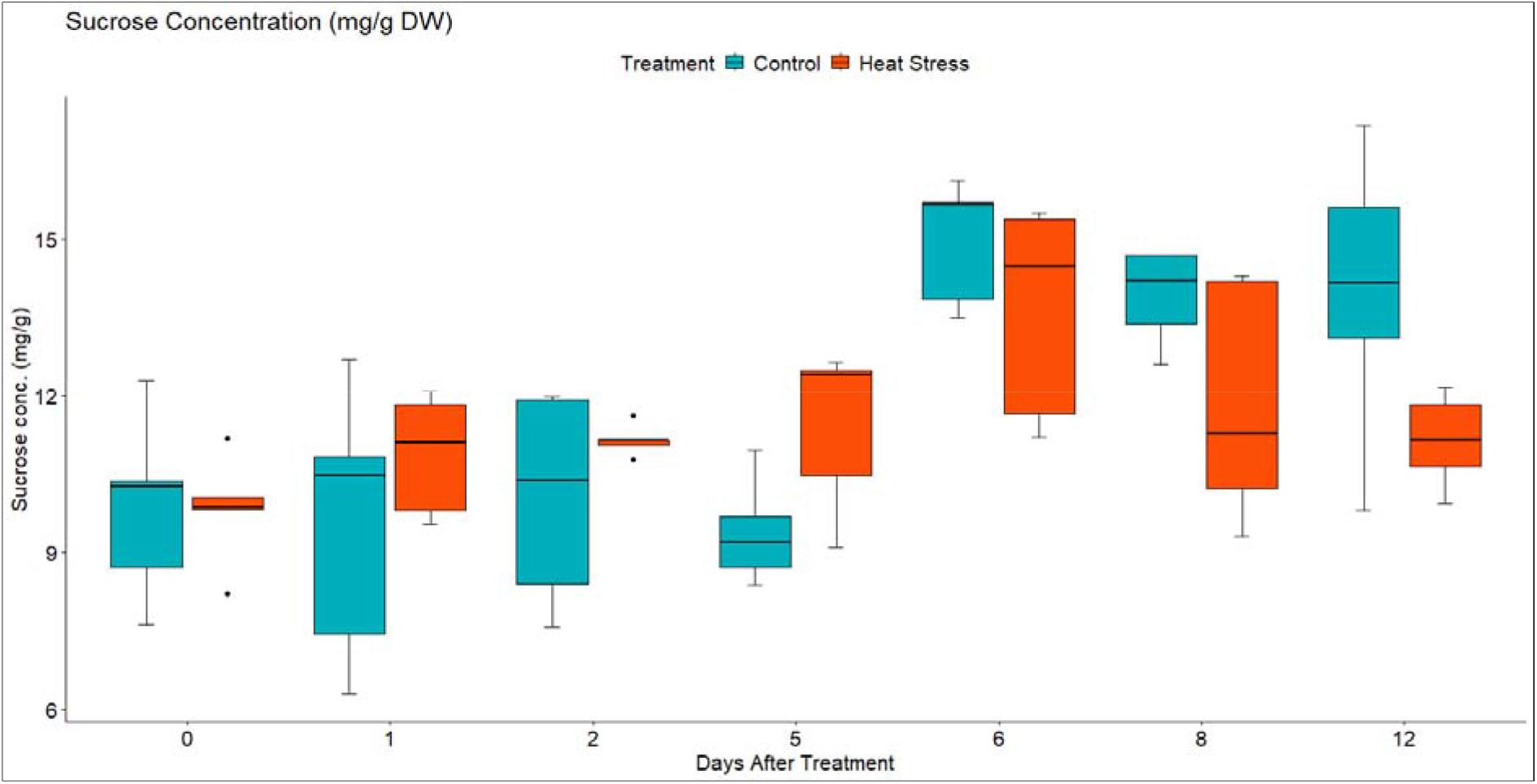
Leaf Sucrose concentration in mg/g of dry weight (DW) in Brassica napus plants subjected to heat treatment (30 °C/24 °C day/night) at timepoints 0-5 (red) or maintained under control temperatures (20 °C/14 °C day/night) at all timepoints (blue). Concentration was quantified using HPLC. Data were analysed using two-way mixed ANOVA pairwise comparisons. Each box plot represents data from 5 replicates, showing the median (horizontal bar), 25^th^ and 75^th^ percentiles (the box) and the min and max values (whiskers), with outliers as dots.

## 4. DISCUSSION

To investigate the effect of prolonged high temperatures (heatwaves) on the growth and adaptability of *B. napus,* especially during its yield-determining reproductive stages, heat treatment experiment was designed in a way that mimic field temperature fluctuations. In the present study, *B. napus* plants were exposed to a gradual increase in temperature for a six-day period and samples were collected at different timepoints during treatment and recovery. The results showed a prevailing effect on the plants both physiologically and metabolically. Heat treatment negatively impacted seed weight and total plant biomass (Supplementary Fig. S4), consistent with similar negative impacts documented in *B. napus* and in other plants such as, Arabidopsis and wheat (Aksouh-Harradj et al., 2006; Zhang et al., 2017; Bheemanahalli et al., 2019). This suggests that heat stress during flowering irreversibly damages vital processes leading to reduced productivity. NIRS showed significant effects of heat treatment on fatty acid concentrations in seeds in response to heat (Supplementary Fig. S5). Several studies also observed heat-induced alteration in lipid composition (Namazkar et al., 2016; Zhou et al., 2018; Zoong Lwe et al., 2021). In *B. napus*, high night temperature has led to overexpression of genes involved in fatty acid catabolism (Zhou et al., 2018). Furthermore, different heat tolerant and susceptible genotypes exhibited different heat-induced changes in their lipid content (Narayanan et al., 2016; Narayanan et al., 2020). Because lipids and proteins are the major constituents of biological membranes, maintaining cellular homeostasis depends mainly on the dynamic nature of lipid composition (Zheng et al., 2011). This suggests an essential role of cellular membrane lipid remodelling under high temperature stress. Additionally, under periods of reduced Carbon availability in plants, the conversion of lipids to organic acids provides an additional source of energy, thus contributing to the plants’ acclimation process (Yuenyong et al., 2019).

At the metabolomic level, heat treatment significantly altered the level of GLS and sugars and led to the differential expression of many genes involved in their synthesis, metabolism and transport. Our study also demonstrated an increase of Glucose, Fructose and Sucrose concentration in leaf in during heat stress, with level slowly reaching, similar or slightly less levels to their control counterpart upon removal and recovery from the stressor. GLS and sugars are among the major secondary metabolites naturally occurring in Brassica species (Ayaz et al., 2006; Rao et al., 2021). Their primary site of synthesis occurs mainly in the leaves, from where they are transported to other parts of the plant (Pacheco-Sangerman et al., 2022; Ren et al., 2022). While reports on the sugars content of leaves are common (Ren et al., 2022; Zafar et al., 2022; Dellero et al., 2024), GLS have been thoroughly investigated in seeds (Wang et al., 2019; Tang et al., 2023; Jhingan et al., 2023), with fewer studies focusing on their presence in leaves (Rhee et al., 2020).

In this study, the levels of GLS and sugars were extracted and quantified from *B. napus* leaves, and their roles in heat stress response were investigated.

### 4.1 Heat stress resulted in down regulation of many genes involved in multiple functional and metabolic processes at 2 DAT

In this study, comparative transcriptome analysis between heat treatment and control identified 9933 significantly differentially expressed transcripts (|log2(foldchange)|≥0.5, FDR ≤ 0.05) (Supplementary Fig. S8-A) with a balanced proportion of up and down regulated genes in contrast HS0vsC0 (0 DAT), and more down regulated than up regulated genes in the rest of the contrasts were identified. This suggests that the impact of heat treatment on the plant’s transcriptome after 24 hours of heat stress, resulted in down regulation of many genes involved in multiple functional and metabolic processes such as Glycolysis, PPP, Citrate Cycle and Oxidative Phosphorylation. In contrast HS12vsC12 (7 DOR), although the proportion of down regulated genes was higher than the up regulated ones, the total number of DEG decreased dramatically as the plant started to restore its metabolic functions after 7 days of recovery.

### 4.2 DEGs showed enrichment of pathways related to respiratory metabolism

The analysis of KEGG enriched pathways revealed the main underlying biochemical pathways altered in *B. napus* in response to heat stress (Supplementary Fig. S16-S19). In accordance with other reports (Estravis-Barcala et al., 2021; Li et al., 2022), DEGs showed an enrichment of pathways related to respiratory metabolism namely Glycolysis, Citrate Cycle, Oxidative Phosphorylation and Pentose Phosphate Pathway (PPP). At 2 DAT, the impact of heat stress on Glycolysis was manifested by the down regulation of the key enzymes Phosphoglucomutase, Aldose 1-Epimerase, Hexokinase, Glucose-6-Phosphate Isomerase, and 6-Phosphofructokinase (Supplementary Fig. S21). Moreover, most of the Citrate Cycle genes were also down regulated at 2 DAT (Supplementary Fig. S22). In the Electron Transport Chain (ETC), gene expression analysis showed that most of the involved genes were down regulated under heat treatment, except for Succinate Dehydrogenase (SDH) which was up regulated (Supplementary Fig. S23). In Mitochondrial metabolism, SDH also known as complex II, plays a central role in the Citrate Cycle and ETC (Huang & Millar, 2013). Additionally, SDH is thought to impact different components of the plant stress response, including stomatal conductance and ROS scavenging (Huang et al., 2019).

Although respiratory pathways have been found to have potential roles in adaptive response to abiotic stress, mainly through reducing the production of ROS (Van Dongen et al., 2011), under extreme conditions such as severe or prolonged heat stress, respiratory enzymes are deactivated, and proteins are denatured leading to culminating of ROS and a total breakdown of mitochondrial respiration (Scafaro et al., 2021). This negatively impacts Oxygen and Carbon fluxes and eventually leads to more severe yield penalty (Rasmusson et al., 2020). In line with this, results from heat stress experiment on seagrass showed that high midday temperature stress of 40°C was associated with significant decrease in biomass (George et al., 2018). Likewise, Imp and co-workers (2019) found that under high night temperature, wheat tolerant cultivar exhibited smaller reductions in biomass and lower rates of both net photosynthesis and respiration compared to other wheat cultivars (Imp et al., 2019).

In the PPP, the oxidative phase (OPPP) was mainly affected at 1 DAT, and this was indicated by the differential expression of Glucose-6-Phosphate Dehydrogenase (G6PDH) and 6-Phosphogluconate Dehydrogenase (6PGDH) enzymes (Supplementary Fig. S24). G6PDH and 6PGDH are the key enzymes in the OPPP. They catalyse the first and third steps of the pathway, respectively (Long et al., 2016), and have been reported to be associated with the response to various abiotic stresses in plants (Gong et al., 2012; Tian et al., 2021). Esposito (2016) reported the involvement of the OPPP in the early response to abiotic stress factors, forming a true metabolic sensor to oxidative stress. At 2 DAT, most of the oxidative and non-oxidative phase enzymes were down regulated, including Transketolase (TK), Transaldolase and Phosphofructokinase enzymes (Supplementary Fig. S25). TK has a central role in primary metabolism, where the products of the involved reactions produce the precursors for nucleic acids biosynthesis, aromatic amino acids, and vitamins (Bi et al., 2013). Bi et al (2013) showed that the cucumber TK gene (*CsTK*) was sensitive to temperature and light (Bi et al., 2013; Bi et al., 2019). Likewise, the Transaldolase gene was found to be involved in the regulation of expression of genes involved in the ABA signalling pathway and the enzymes responsive to ROS (Rong et al., 2021). The results of this study add further evidence to the involvement of the PPP enzymes in the heat stress response in *B. napus*.

### 4.3 Heat stress altered GLS metabolism during treatment

At the transcriptomic level, heat treatment differentially affected the expression of GLS synthesis and regulatory genes (Figs. 3 & 4). Several genes encoding different UDP Glycosyltransferases and Glutathione S-Transferases appeared down regulated, while others encoding GLS related transcription factors were up regulated, suggesting that these genes might have participated at different stages of the stress response.

The influence of heat stress on individual GLS was further evaluated through metabolomic analysis of two aliphatic GLS (Progoitrin (PRO) and Gluconapin (GNA)), and one indolic GLS (Glucobrassicin (GBS)). The role of these GLS in response to different abiotic stress factors has been previously reported (Jasper et al., 2020; Ljubej et al., 2021). In the current study, the concentration of PRO and GNA significantly increased in response to heat stress before it started to decline at 5 DAT (Figs. 6 & 7). This decline could have resulted from membrane damage caused by prolonged exposure to heat, leading to GLS degradation. Besides, the removal of heat stress after 5 DAT could also be another factor for the decline in GLS content. In addition to its primary role in response to plant pathogen interaction (Hopkins et al., 2009; Touw et al., 2020), GLS accumulation in response to heat stress has also been previously reported (Valente Pereira et al., 2002; Martínez-Ballesta et al., 2013; Jasper et al., 2020). When plants are stressed, growth is reduced, and Carbon utilization is predominantly diverted towards the production of secondary metabolites. As part of the plant defence mechanism, GLS play important role as osmoprotective compounds (Martínez-Ballesta et al., 2013), where their concentration increases in response to stress, providing protection against oxidative damage. At 30/15°C (day/night) temperature regime, *Brassica oleracea* seedlings had significantly higher GLS concentration than plants cultivated at lower temperatures (22/15°C and 18/12°C) (Valente Pereira et al., 2002). Similarly, an Arabidopsis GLS mutant experienced a remarkable decline in growth and development and was found to be less heat tolerant than wild-type plants upon exposure to high temperature (Ludwig-Müller et al., 2000).

In contrast, indolic glucosinolate GBS showed a downward trend where concentration decreased gradually in response to heat (Fig. 8). Similar results have been found where GBS concentration was reduced in response to heat treatment (Bohinc & Trdan, 2012). Also, during short term high temperature stress, the transcript level of the indolic glucosinolates synthetic genes were predominantly down regulated (Rao et al., 2021). Several studies reported that indole GLS are much more sensitive to heat treatment (Bones & Rossiter, 2006; Bohinc & Trdan, 2012) and can demonstrate more thermal degradation than aliphatic GLS at lower temperatures (Orlemans et al., 2006).

The results of our study indicate that the concentration of GLS in the leaves of *B. napus* is influenced by high temperature stress, which was previously reported in other Brassica species (Rask et al., 2000; Bohinc & Trdan, 2012). It also shows that different groups of GLS (aliphatic versus indolic) differ in their response to heat stress. As reported by Ljubej et al., (2021), the different accumulation trends of these groups could correspond to different protective mechanisms played by individual GLS (Ljubej et al., 2021). Additionally, the variety of enzymes involved in GLS synthesis and regulation could be another factor for the difference in heat response (Schonhof et al., 2007).

### 4.4 Heat stress altered the expression of Sulphur assimilation and transport genes

In the current study, the transcript levels of Sulphur assimilation and transport genes were significantly down regulated especially at 2 DAT (Fig. 5). This included several Adenylyl Sulphate Kinases and Reductases, ATP Sulphurylase (ATPS) and Sulphate transporter genes. These genes are important for the activation, catalysis and transport of Sulphur compounds (Capaldi et al., 2015). Moreover, enrichment pathway analysis showed that Cysteine and Methionine metabolism pathway was among the top significantly enriched pathways at 1 & 2 DAT (Supplementary Figs. 16-17). In this pathway, 11 transcripts mapping to S-Adenosylmethionine Synthetase and 11 transcripts mapping to Homocysteine S-Methyltransferases were down regulated (Supplementary Fig. S20). These enzymes are among the key enzymes that control the Methionine pool (Capaldi et al., 2015). Given that GLS compounds are rich in Sulphur, assimilation of Sulphur is closely associated with GLS biosynthesis. Cysteine which is the terminal metabolite in Sulphur assimilation, acts as a Sulphur donor for Methionine, a precursor for aliphatic GLS. This renders Cysteine an important intersection point between Sulphur assimilation and GLS biosynthesis (Rao et al., 2021). Such proximity between Sulphur and these amino acids makes Cysteine and Methionine metabolism pathway directly related to GLS biosynthesis.

### 4.5 Heat stress altered sugars metabolism during treatment and recovery periods

In the present study, heat treatment altered carbohydrate concentration as the amounts of Glucose, Fructose and Sucrose increased with heat treatment (Figs. 11-13). At the transcriptional level, the starch hydrolysing enzyme Alpha-Amylase was induced. Conversely, the Starch synthesizing enzymes such as Phosphoglucomutase (PGM), AGPase and Hexokinase were inhibited (Supplementary Figs. S21 & S27). Likewise, heat induced the up regulation of Sucrose synthesizing enzymes and the down regulation of Sucrose hydrolysing enzymes such as Cell Wall Invertase and Sucrose Synthase (Supplementary Tables S5 & S6). Together with the inhibition of many sugars transporters such as SWEET and ERD6-like (Figs. 9 & 10) suggest that heat induced Starch hydrolysis and promoted Sucrose synthesis and accumulation in the leaves. Several reports showed that different environmental stress affect sugars metabolism and transport (Xalxo et al., 2020; Mathan et al., 2020). For example, drought and salinity stresses increased Sucrose content in leaf and root tissues of rice (Mathan et al., 2020). Similarly, a significant accumulation of total sugars and Proline was observed in different genotypes of heat stressed seedlings of moth bean (Harsh et al., 2016). Marias et al. (2017) also reported an accumulation of Glucose and Fructose under heat stress in the leaves of *Coffea arabica* (Marias et al., 2017). In maize ear leaf, the ^13^C export rate was decreased by high temperature treatment, resulting in growth enhancement of vegetative plant parts and reduction in grain yield (Suwa et al., 2010). During heat stress, leaves are the first tissues to sense and experience damage by heat. Consequently, leaves use sugars to efficiently scavenge ROS and alleviate oxidative damage. In high respiration environment, it has been hypothesized that sugars acts as energy supply to ensure survival (Burke, 2007), as their role in osmotic readjustments helps to maintain cell-water balance and membrane integrity. Thereby, maintaining leaf sugars content through reducing source to sink export is considered a tolerance strategy (Kaushal et al., 2013). Additionally, the decrease in sink demand due to growth limitation, could also contribute to the accumulation of sugars in the source leaves under stressful conditions (Hummel et al., 2010; Lemoine et al., 2013).

## 5. CONCLUSION

Through a comprehensive transcriptomic and metabolomic analysis, this study brings evidence for the effect of high temperature stress and in particular heatwaves on different functional and metabolic pathways, highlighting its role in the plant stress defence system. More specifically, our results indicated that heat treatment:

1. Inhibited important metabolic pathways such as respiratory metabolism namely Glycolysis, Pentose Phosphate Pathway, Citrate Cycle and Oxidative Phosphorylation especially after 48 hours of heat treatment.
2. Reduced yield and plant biomass; and altered seed composition.
3. Altered sugars and glucosinolates levels in leaves.
4. Induced the expression of 9933 genes which were differentially regulated during heat treatment and recovery. Most of the top up and down regulated genes involved in key biological processes such as heat shock proteins, cellular processes regulation, transcription factors, cell wall remodelling, sugars and secondary metabolites transport and metabolism.

Finally, the gradual increase in temperature adapted in the heat experiment helped to create an environment that mimics field conditions. Thus, this study represents an important step towards developing an understanding of the heat stress response and tolerance mechanisms, a knowledge that could be transferred to other plants.

## Authors Contributions

All authors contributed to the experimental design of the study. MK undertook the research, carried out the lab work, conducted the bioinformatics and data analysis, and drafted the manuscript. FM supervised the bioinformatics analysis. MA supervised the metabolomic lab work and data analysis. JPH supervised the molecular genetics lab work. All authors contributed to and revised the manuscript, given the opportunity to comment, and approve the final version of the manuscript.

## Acknowledgment

We thank Dr Luke Bell from the University of Reading (School of Agriculture, Policy & Development) for providing the reference standards for glucosinolates and Monika Jodkowska from Cranfield University (Plant Science Laboratory) for developing the method for the LC-MS experiment.

## Data Availability Statement

Raw sequencing for the RNA-Seq analysis have been deposited at the NCBI’s Sequence Read Archive, under the submission number SUB14296957. All other datasets supporting the results of this article are included within the article and supplementary materials.

