## Supplementary Materials for "Prolonged heat stress in *Brassica napus* during flowering negatively impacts yield and alters glucosinolate and sugars metabolism"

### <u>Supplementary Figures and Tables:</u>

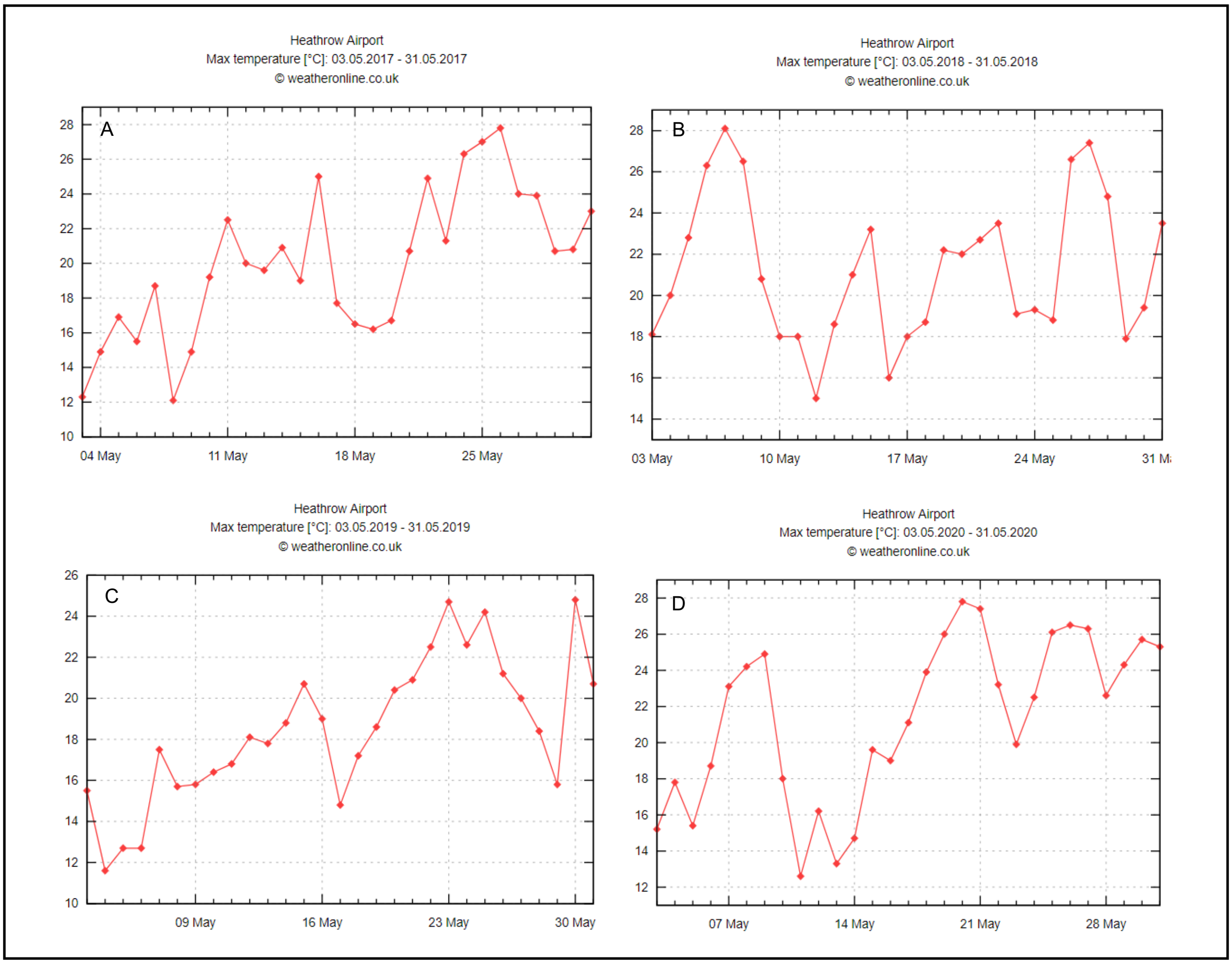

Figure S1. Maximum daily temperature (°C) recorded at Heathrow airport during May 2017 (A), May 2018 (B), May 2019 (C) and May 2020 (D). Records show an increase in the frequency of temperature surges during May.

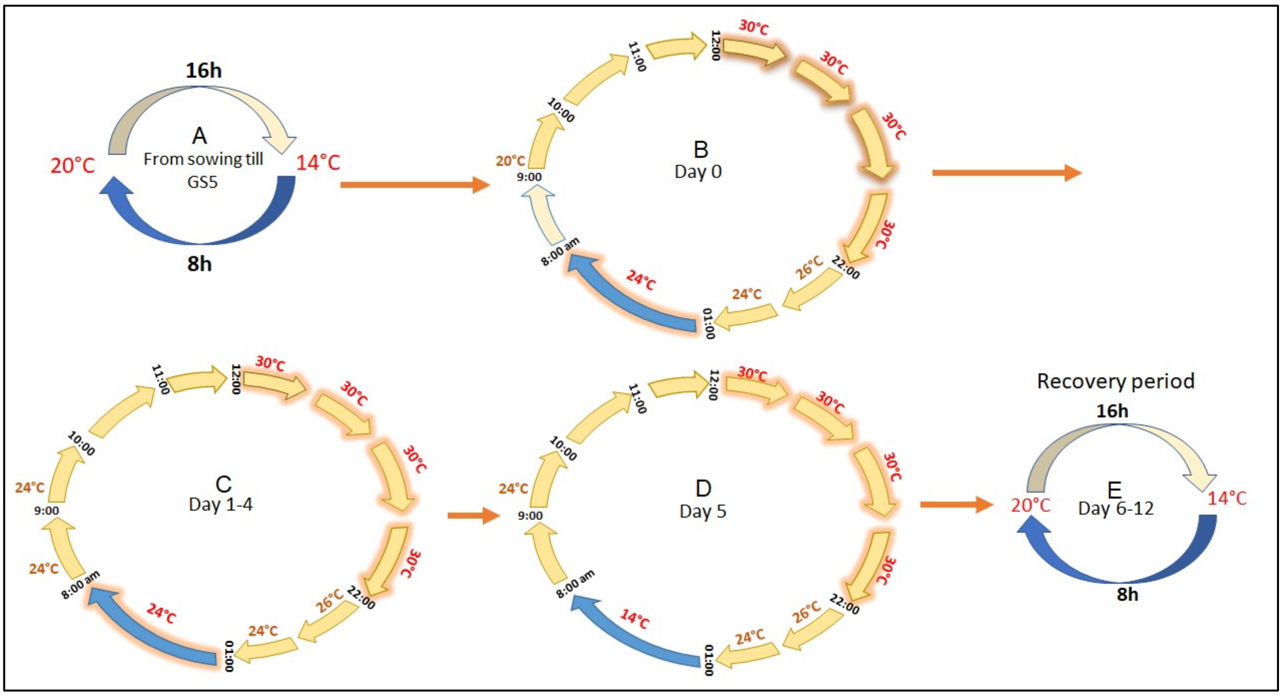

Figure S2. Heat treatment experimental design. To mimic a heatwave, the temperature in the heat treatment cabinet was increased gradually from 20 °C to 30 °C between 9:00 h and 12:00 h. The temperature was held at 30 °C until 22:00 h (B), before it was dropped gradually and maintained at 24 °C until 8:00 h of the next day (B). This cycle of gradual increase (day) and decrease (night) of temperature was held for the next 5 days (C). On the 6$^{th}$ day of treatment, the temperature was gradually decreased to 20 °C/14 °C day/night cycle (D) and held for a recovery period of 7 days (E).

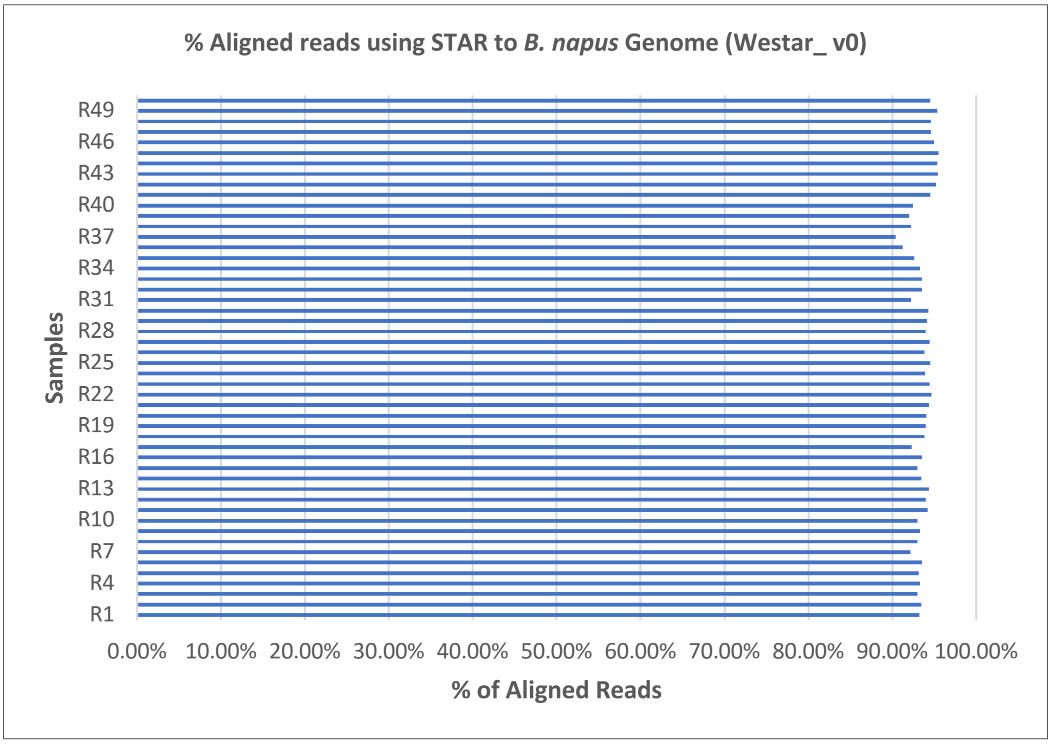

Figure S3. Percentage of aligned reads per sample. Around 90%- 95.49% total reads per sample were mapped to the *B. napus* cv. Westar v0 reference genome.

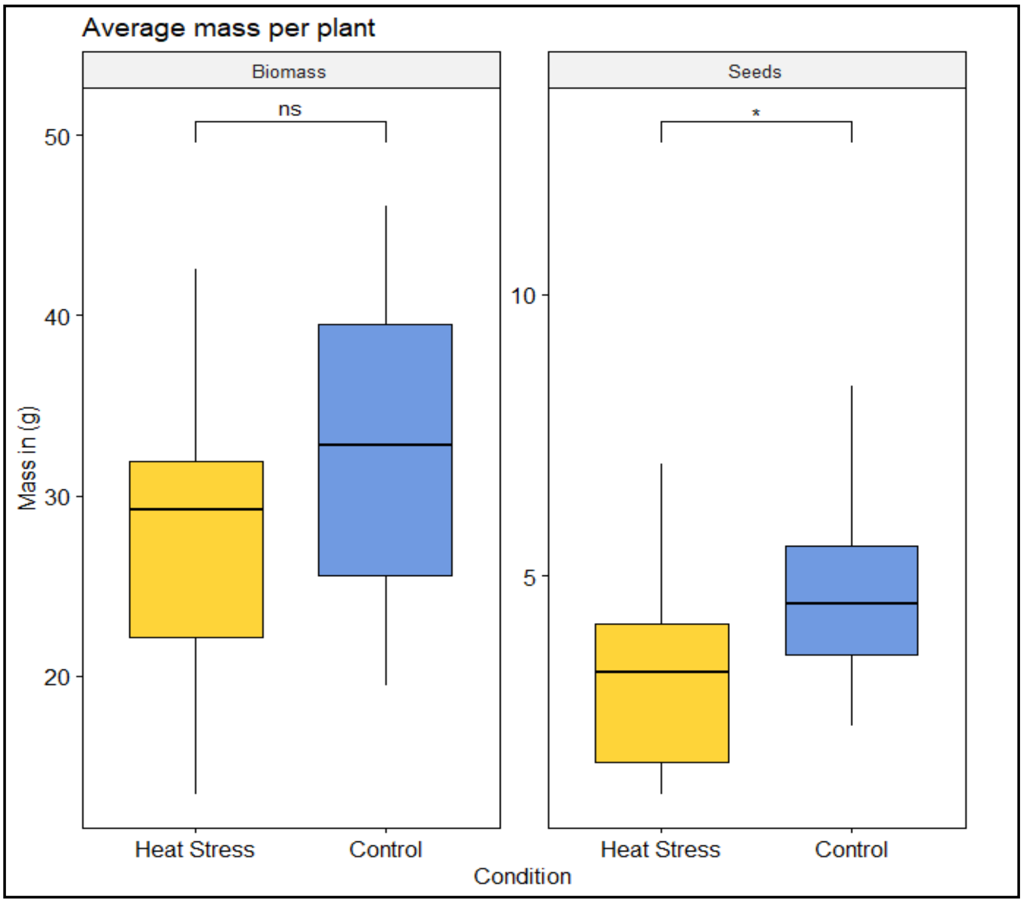

Figure S4. Above ground total biomass and seed yield per plant at maturity. Heat-treated plants (yellow) exhibited less total biomass and seed weight relative to control plants (blue).

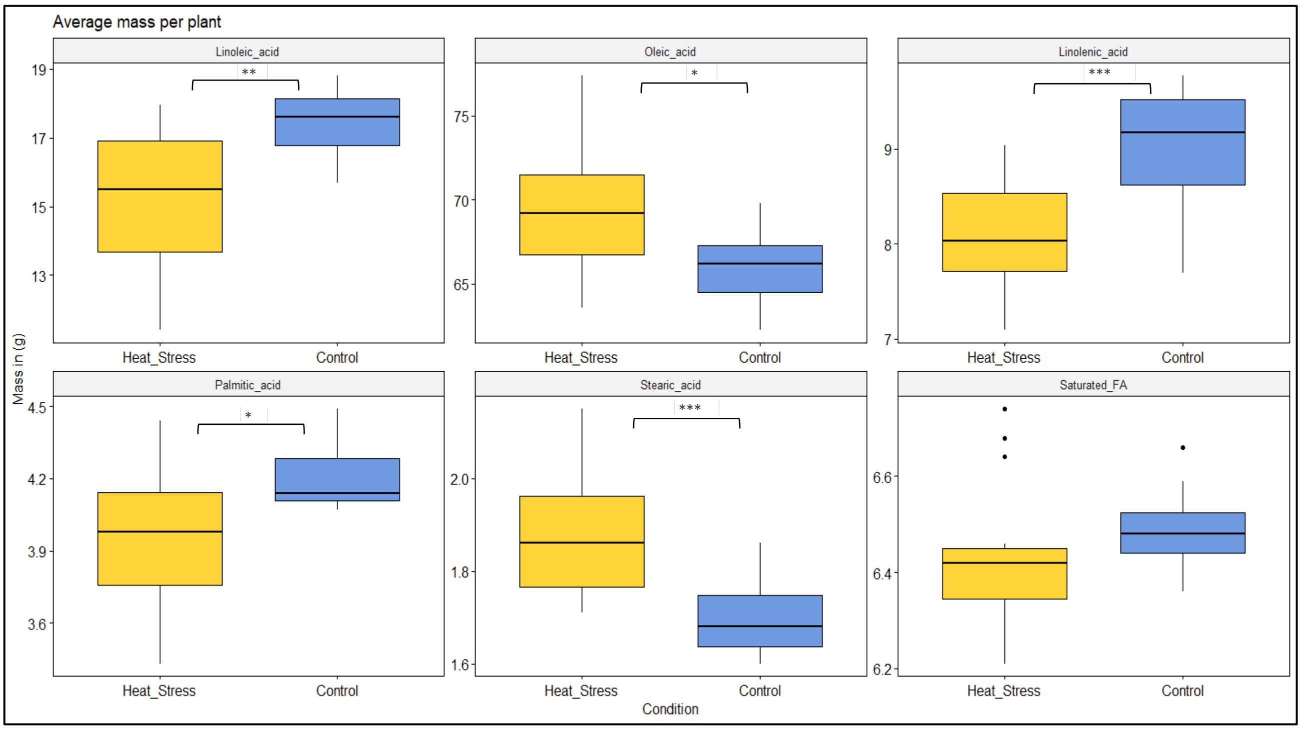

Figure S5. Fatty acid content in seeds per plant. Seeds from heat-treated plants (yellow) exhibited significant decrease in Linoleic Acid, Linolenic Acid and Palmitic Acid concentrations, but an increase in Oleic Acid and Stearic Acid concentrations compared to seeds from control plants (blue).

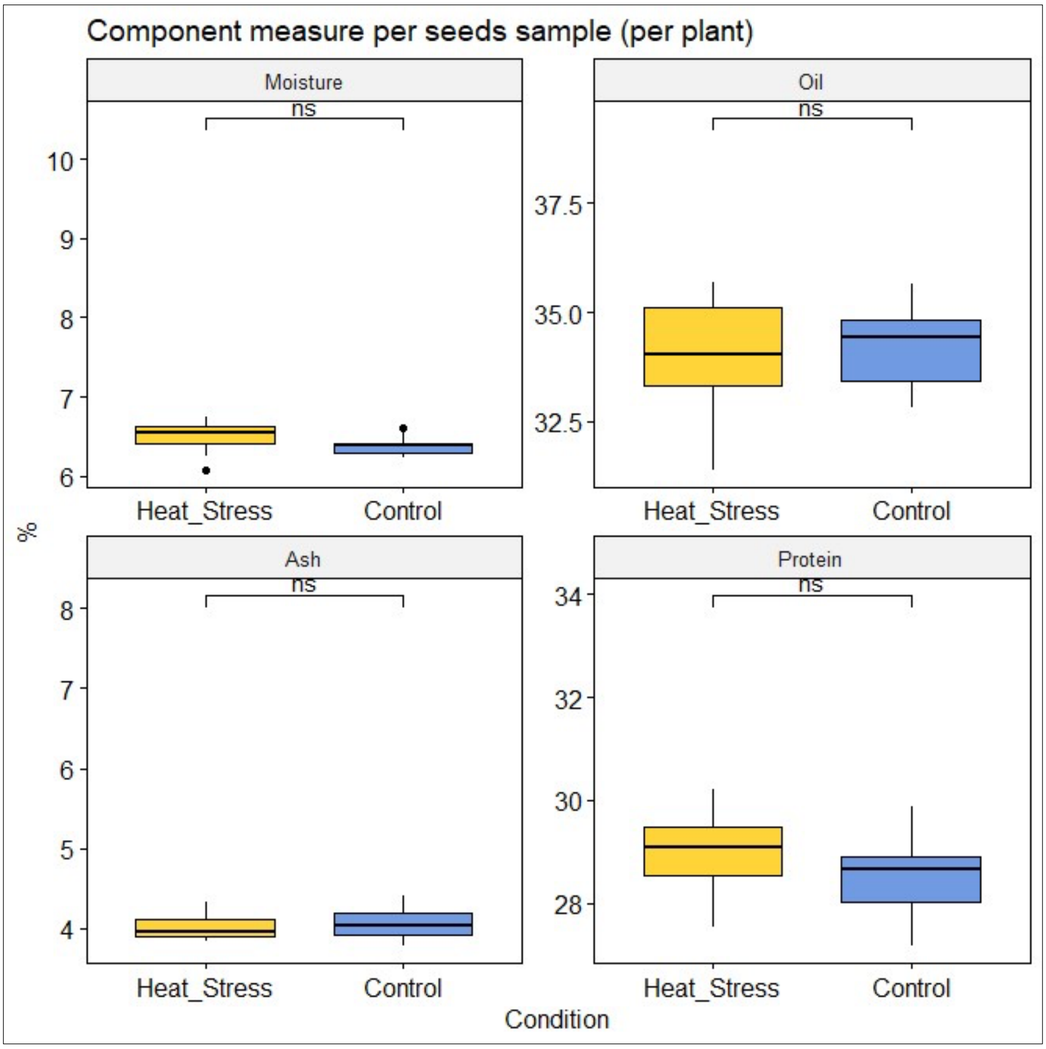

Figure S6. Moisture, oil, ash and protein content in seeds per plant.

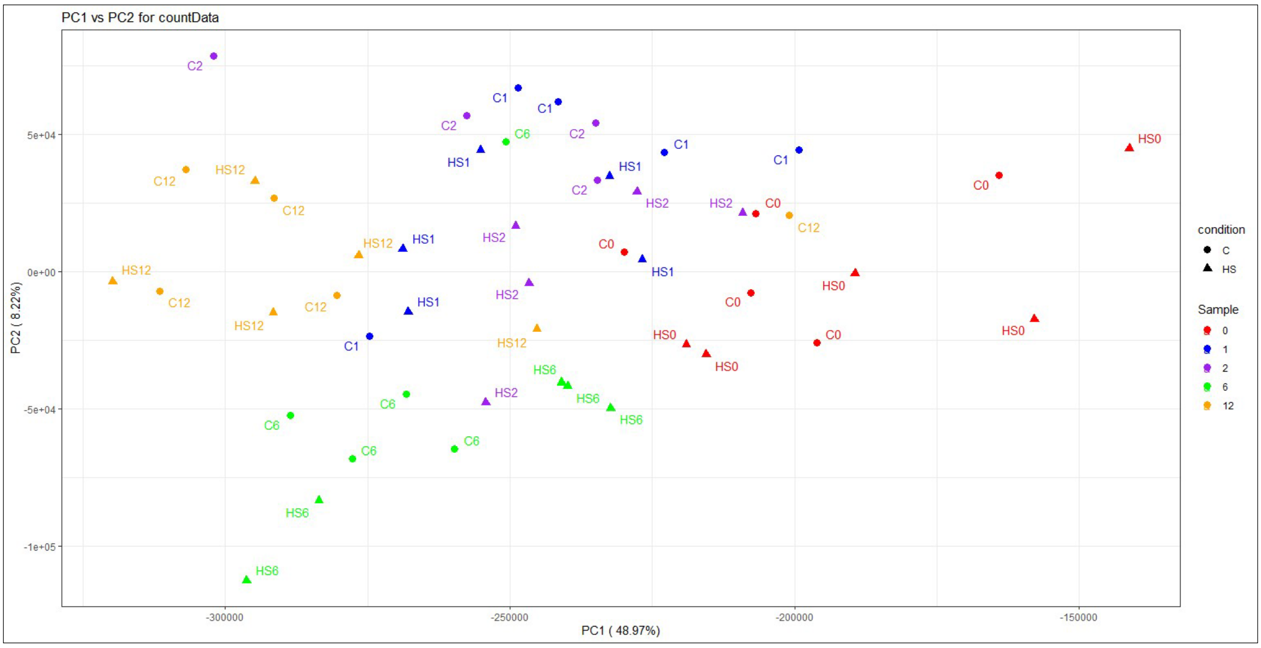

Figure S7. Principle component analysis of the control and heat treatment samples: Samples distribution in PC1 and PC2.

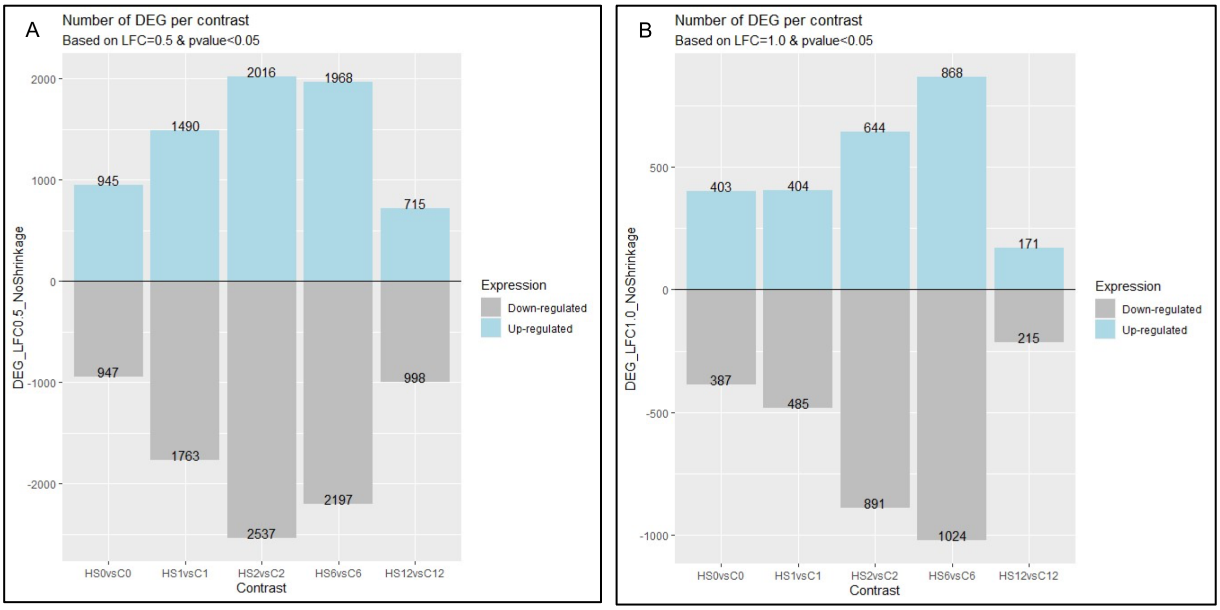

Figure S8. Pairwise comparison of the number of DEG between heat treatment and control at each timepoint based on LFC =0.5 (A) & LFC = 1.0 (B) and p-value ≤0.05. HS0vsC0, HS1vsC1, HS2vsC2, HS6vsC6, and HS12vsC12 contrasts correspond to 0, 1, 2 days after treatment (DAT) & 7 days of recovery (DOR) respectively.

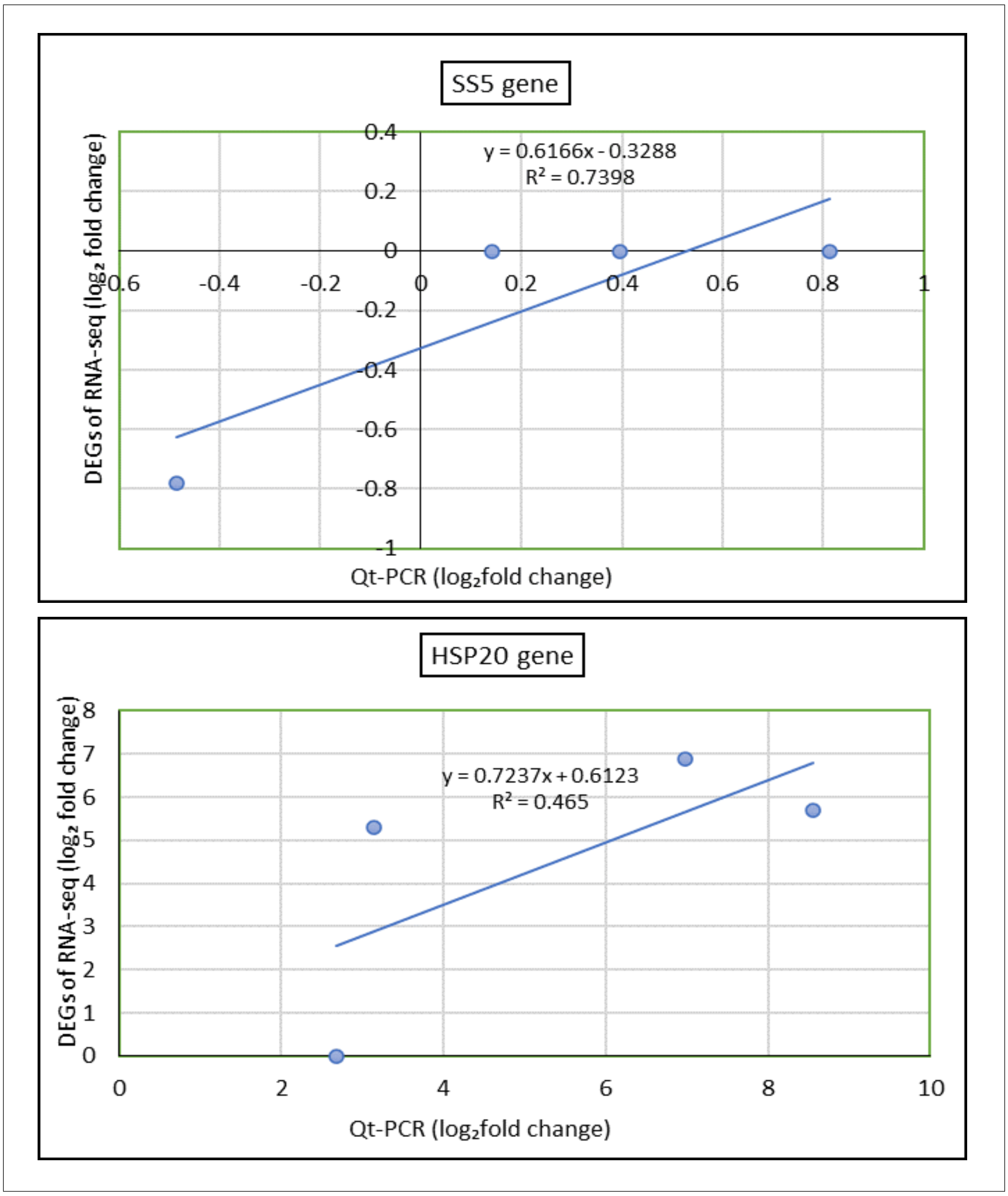

Figure S9. Correlation analysis of the results obtained from RNA-seq and qt-PCR: HSP20 and SS5 exhibited similar expression profiles between RT-qPCR and RNA-seq data with correlation coefficients of r=0.86, and r=0.68 respectively.

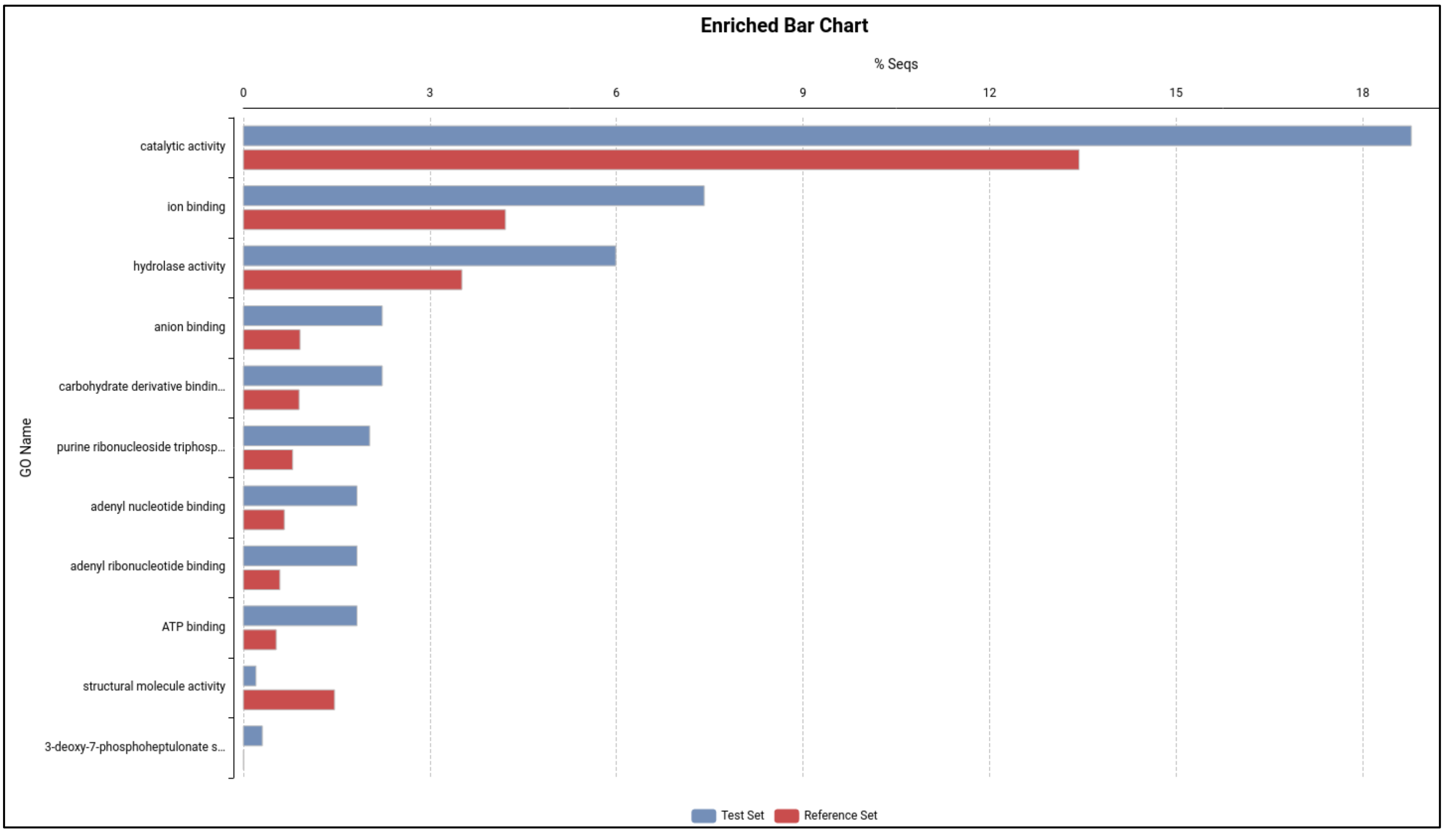

Figure S10. Enriched GO terms at 0 DAT, as identified by GO enrichment analysis.

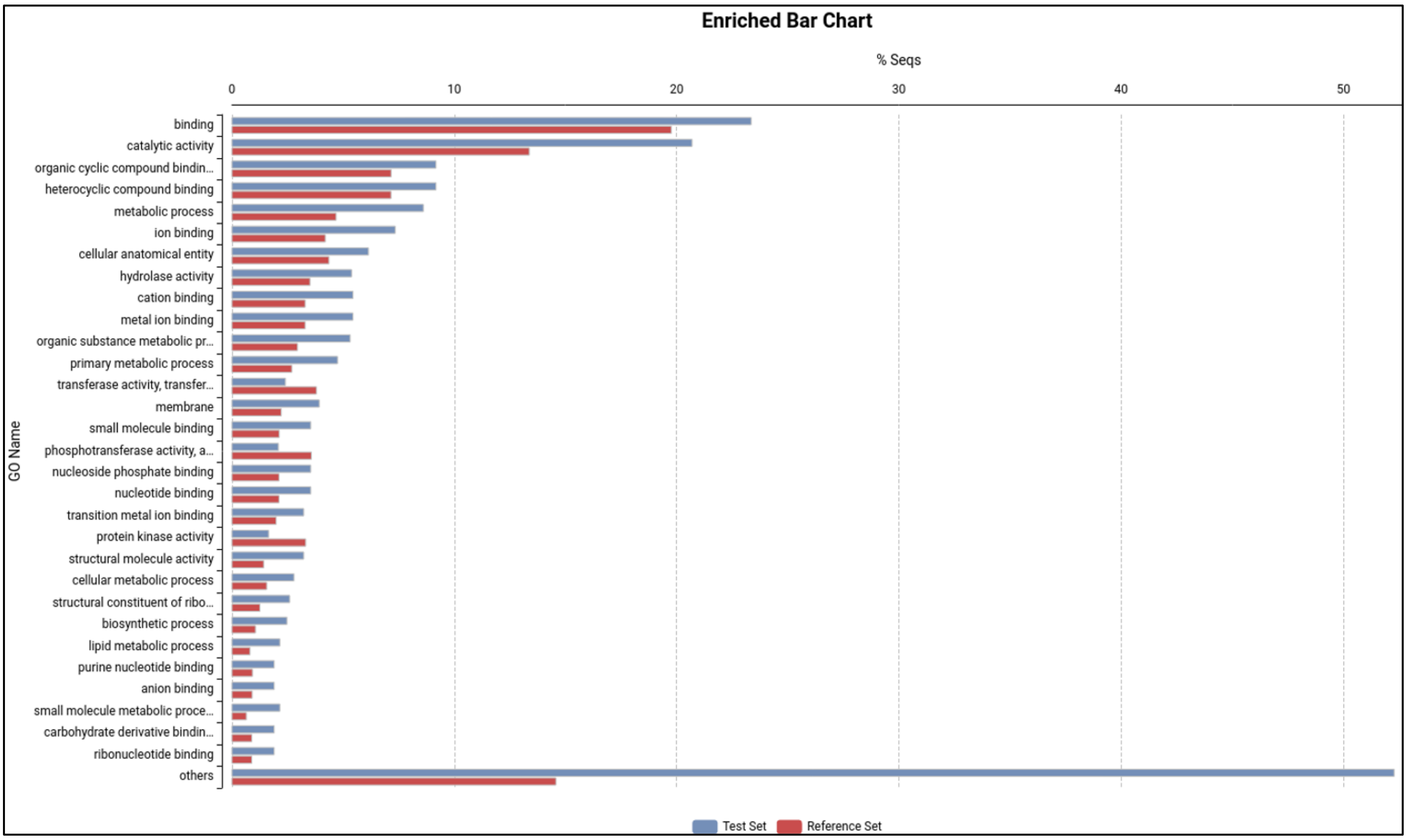

Figure S11. Enriched GO terms at 1 DAT, as identified by GO enrichment analysis.

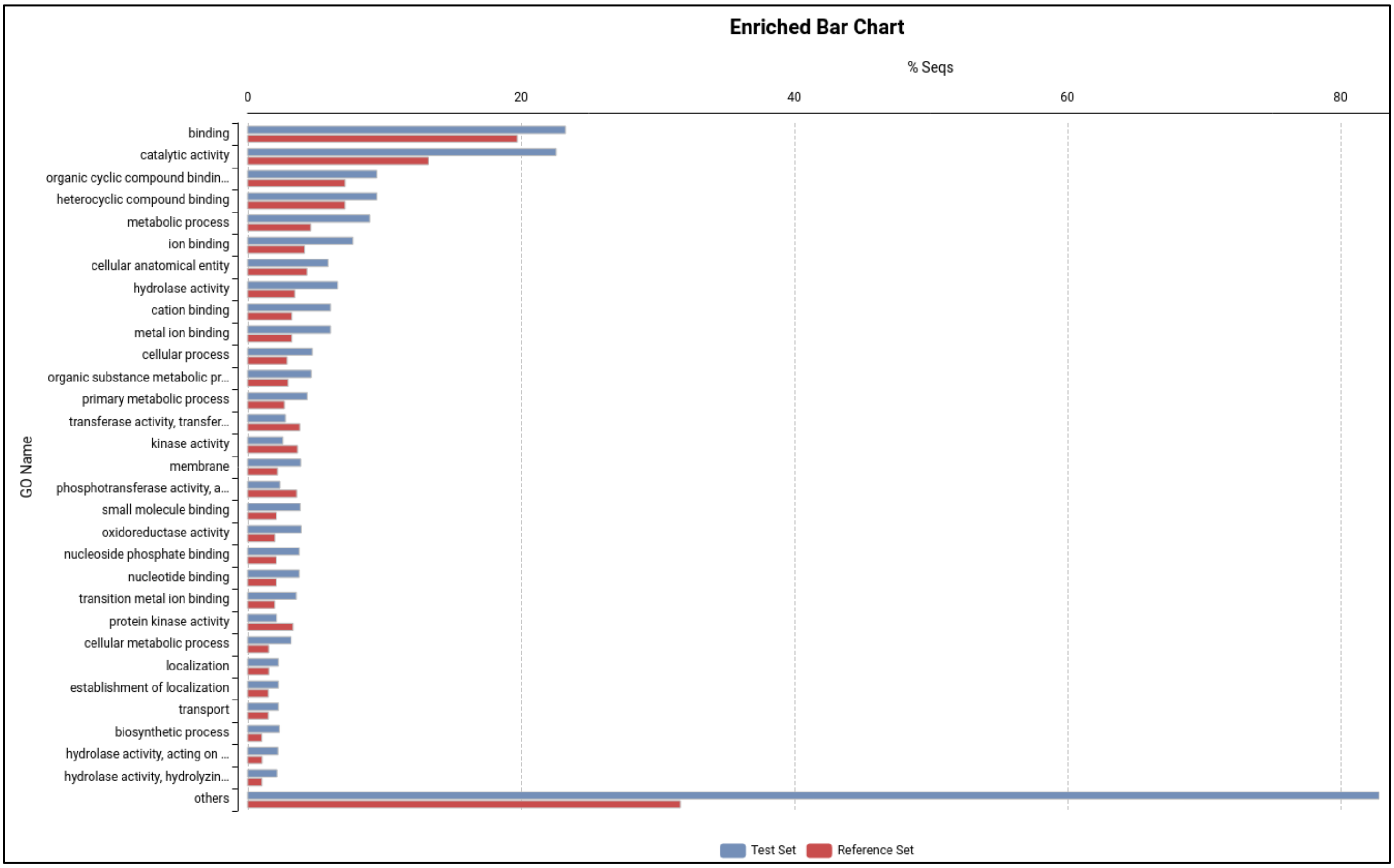

Figure S12. Enriched GO terms at 2 DAT, as identified by GO enrichment analysis.

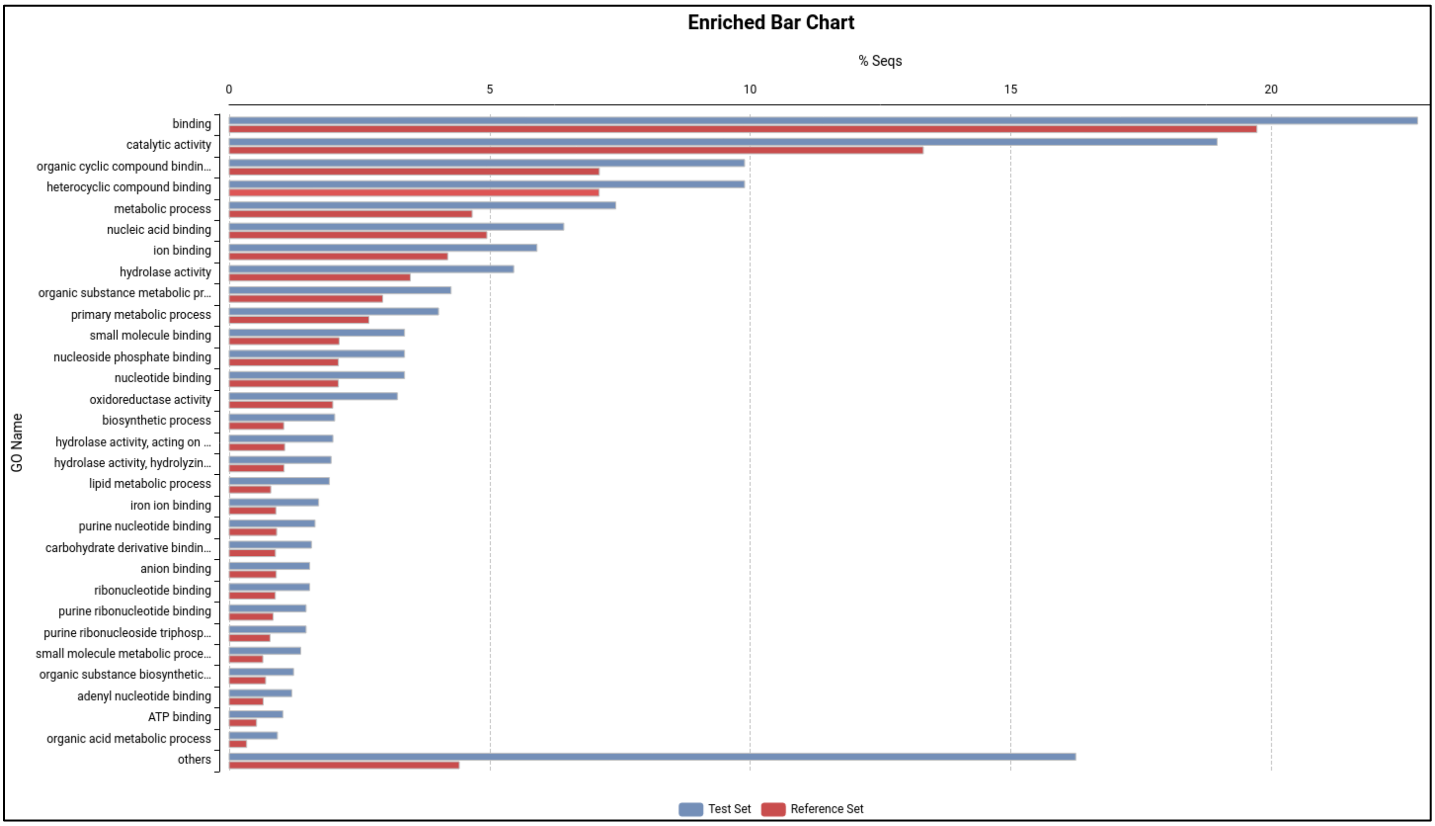

Figure S13. Enriched GO terms at 1 DOR, as identified by GO enrichment analysis.

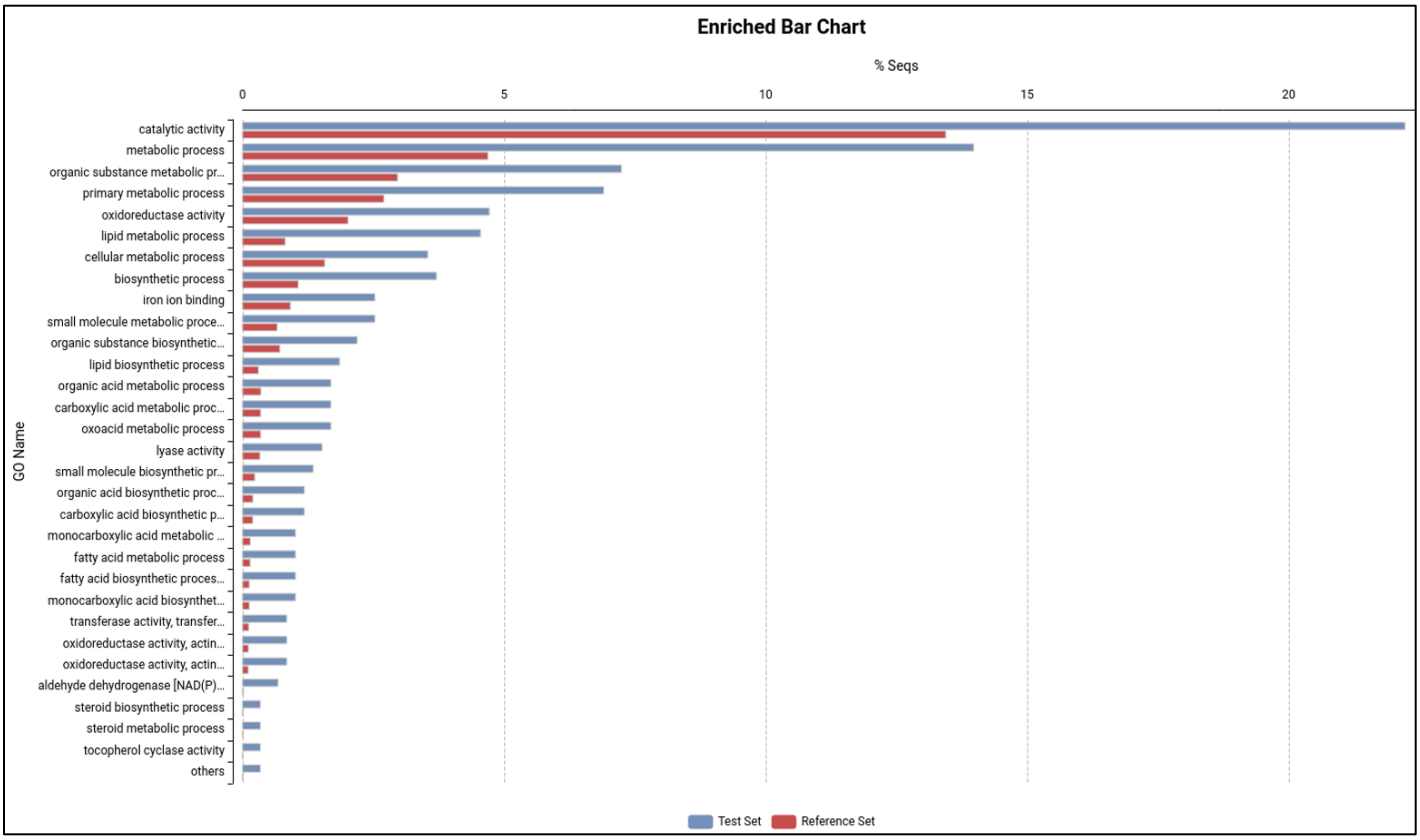

Figure S14. Enriched GO terms at 7 DOR, as identified by GO enrichment analysis.

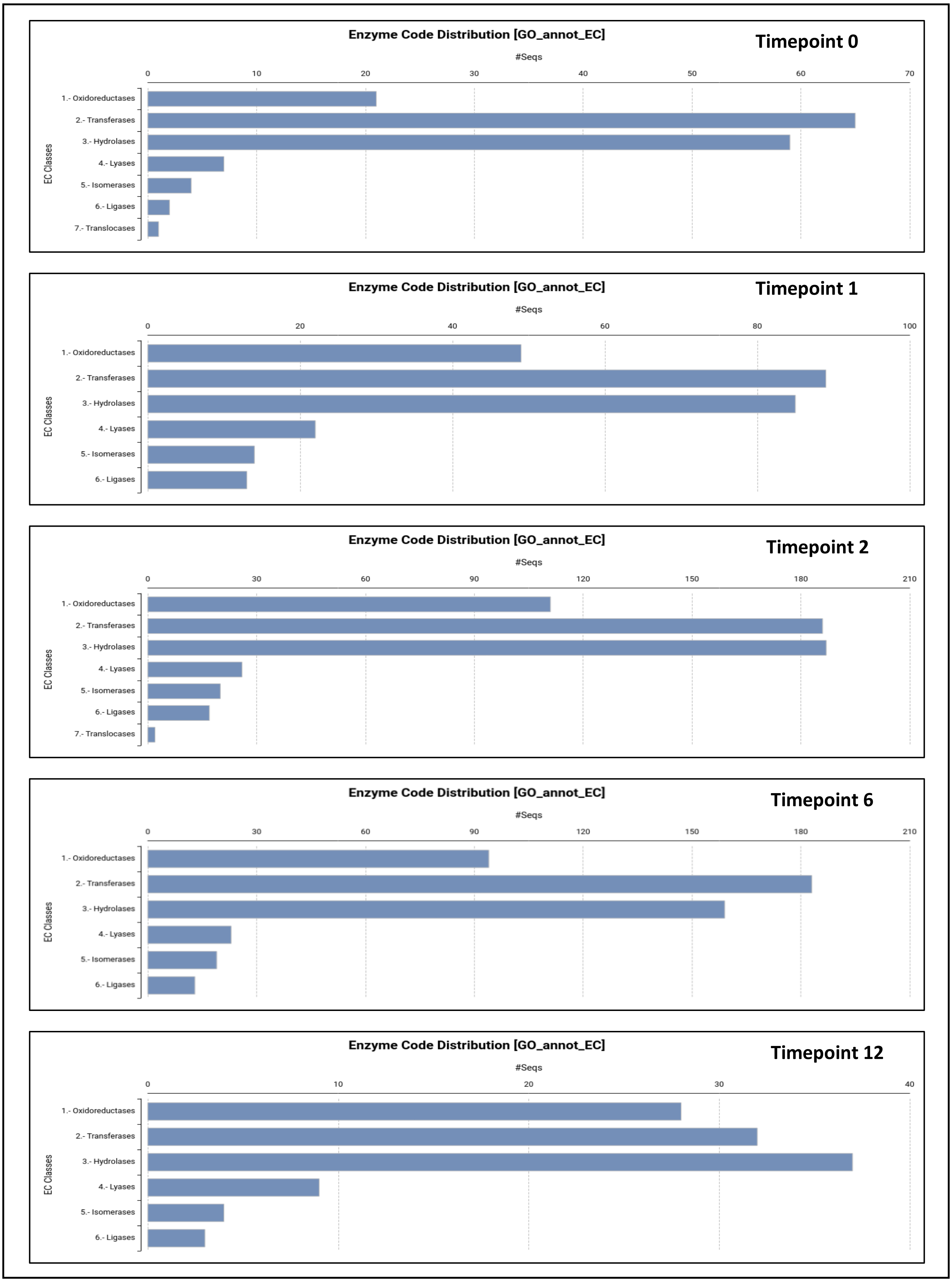

Figure S15. Enzyme code distribution at different timepoints as identified by GO enrichment analysis.

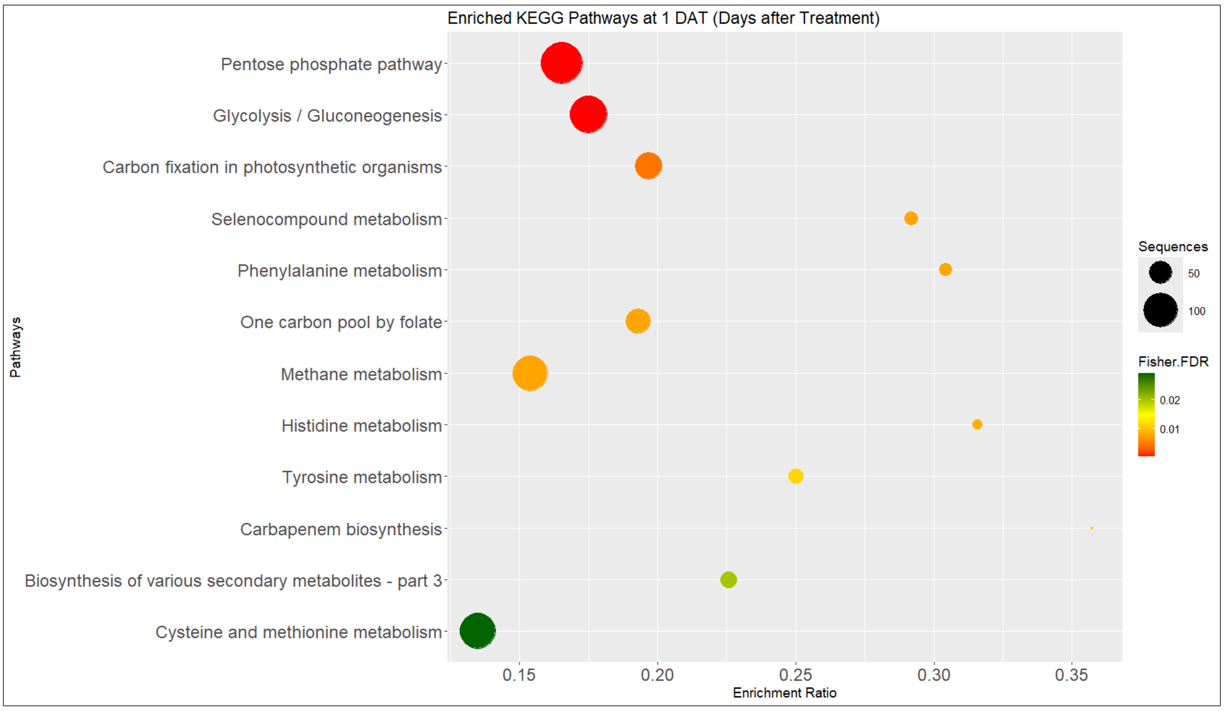

Figure S16. The top KEGG pathways and their corresponding number of genes at 1 DAT at p-value ≤ 0.05. The x-axis represents the enrichment factor. The y-axis represents KEGG pathways. The circle represents the number of genes mapped to pathways and the red to green bar indicates the significance.

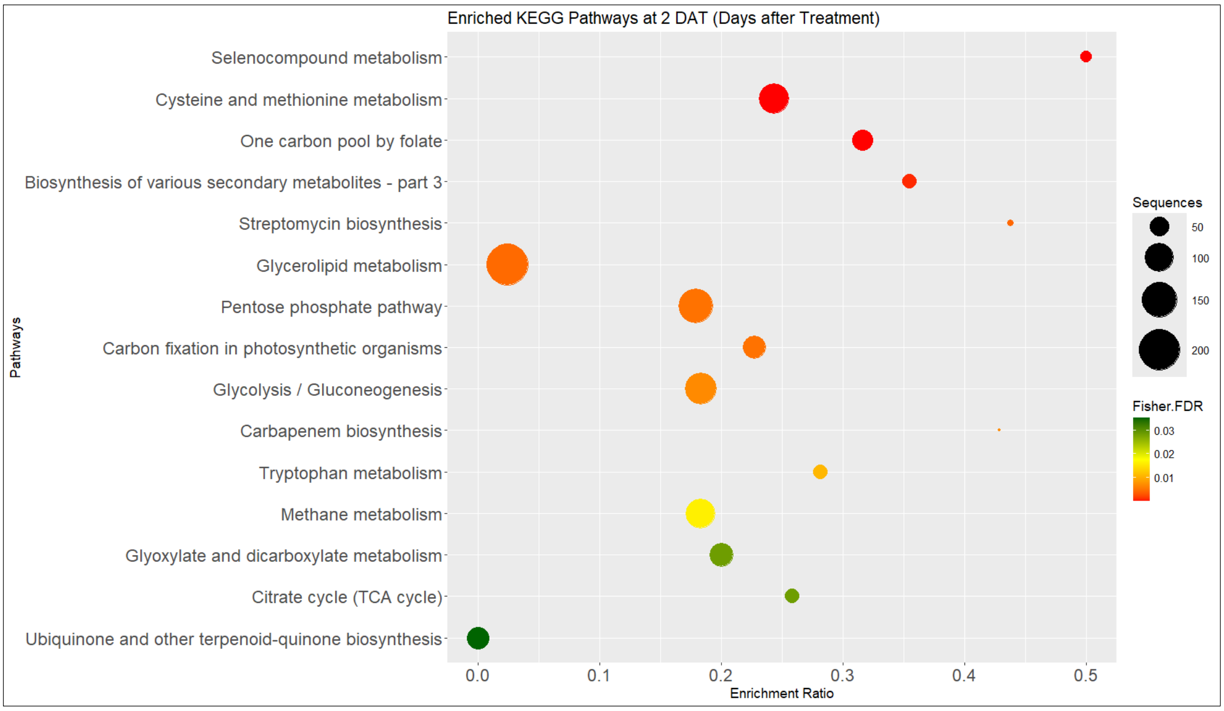

Figure S17. The top KEGG pathways and their corresponding number of genes at 2 DAT at p-value ≤ 0.05. The x-axis represents the enrichment factor. The y-axis represents KEGG pathways. The circle represents the number of genes mapped to pathways and the red to green bar indicates the significance.

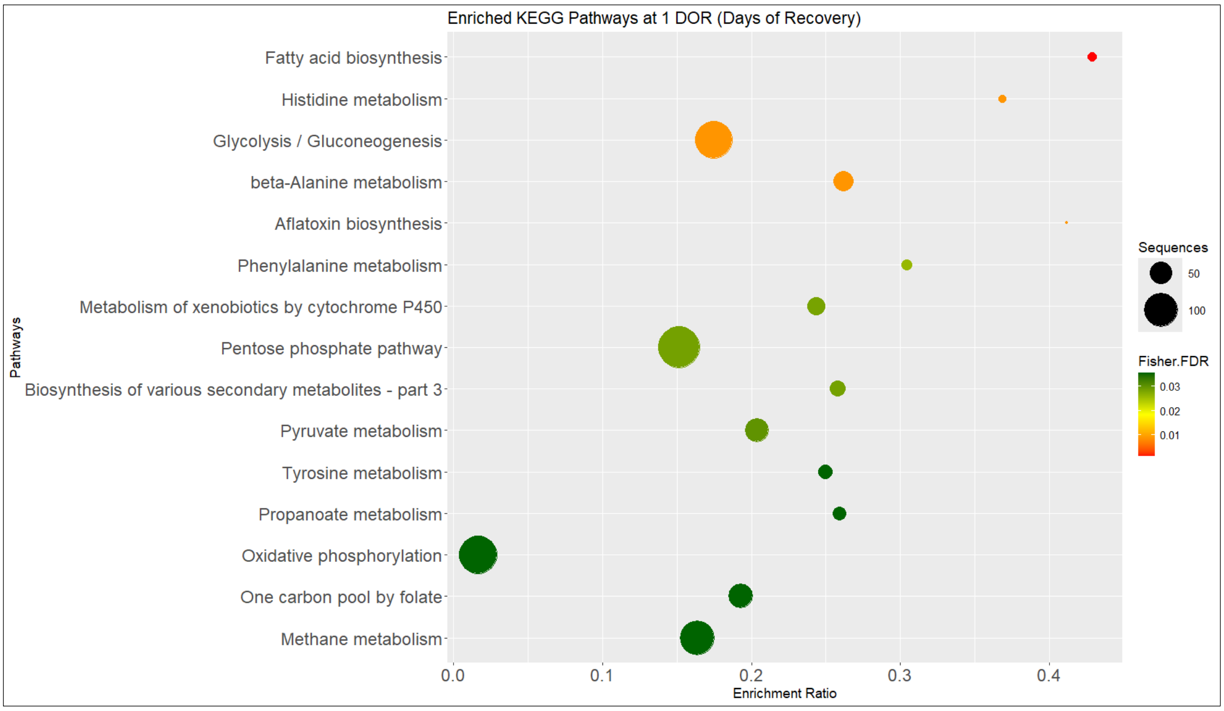

Figure S18. The top KEGG pathways and their corresponding number of genes at 1 DOR at p-value ≤ 0.05. The x-axis represents the enrichment factor. The y-axis represents KEGG pathways. The circle represents the number of genes mapped to pathways and the red to green bar indicates the significance.

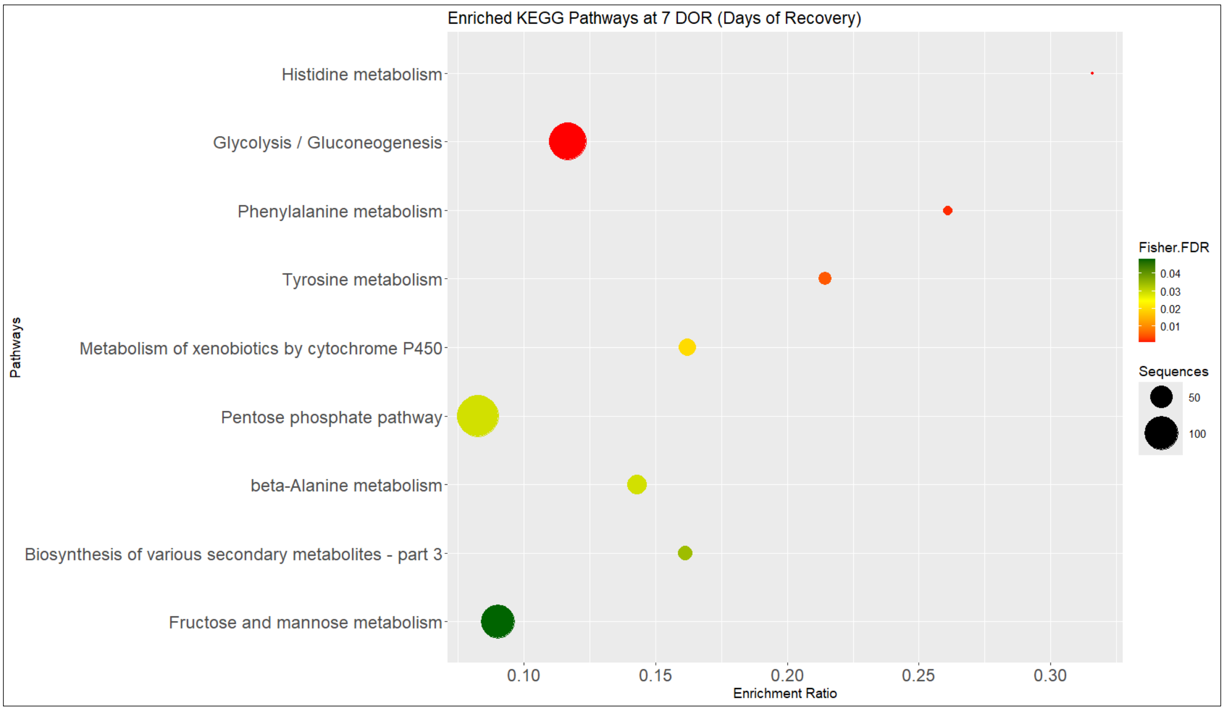

Figure S19. The top KEGG pathways and their corresponding number of genes at 7 DOR at p-value ≤ 0.05. The x-axis represents the enrichment factor. The y-axis represents KEGG pathways. The circle represents the number of genes mapped to pathways and the red to green bar indicates the significance.

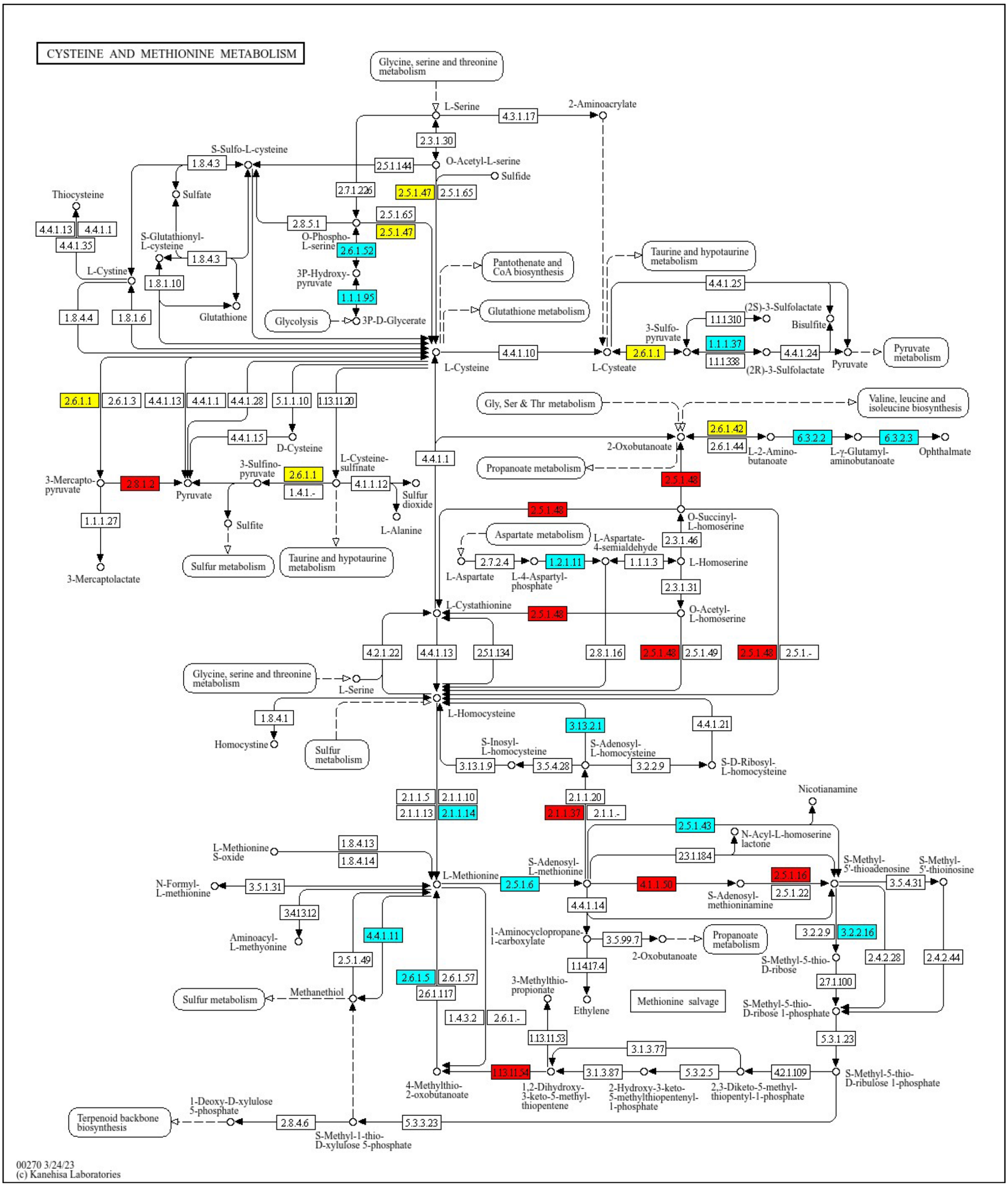

Figure S20. KEGG pathway for Cysteine and Methionine metabolism at 2 DAT.

*Blue denotes down regulated genes, red denotes up regulated genes and yellow denotes some isoforms were up regulated and some isoforms were down regulated.

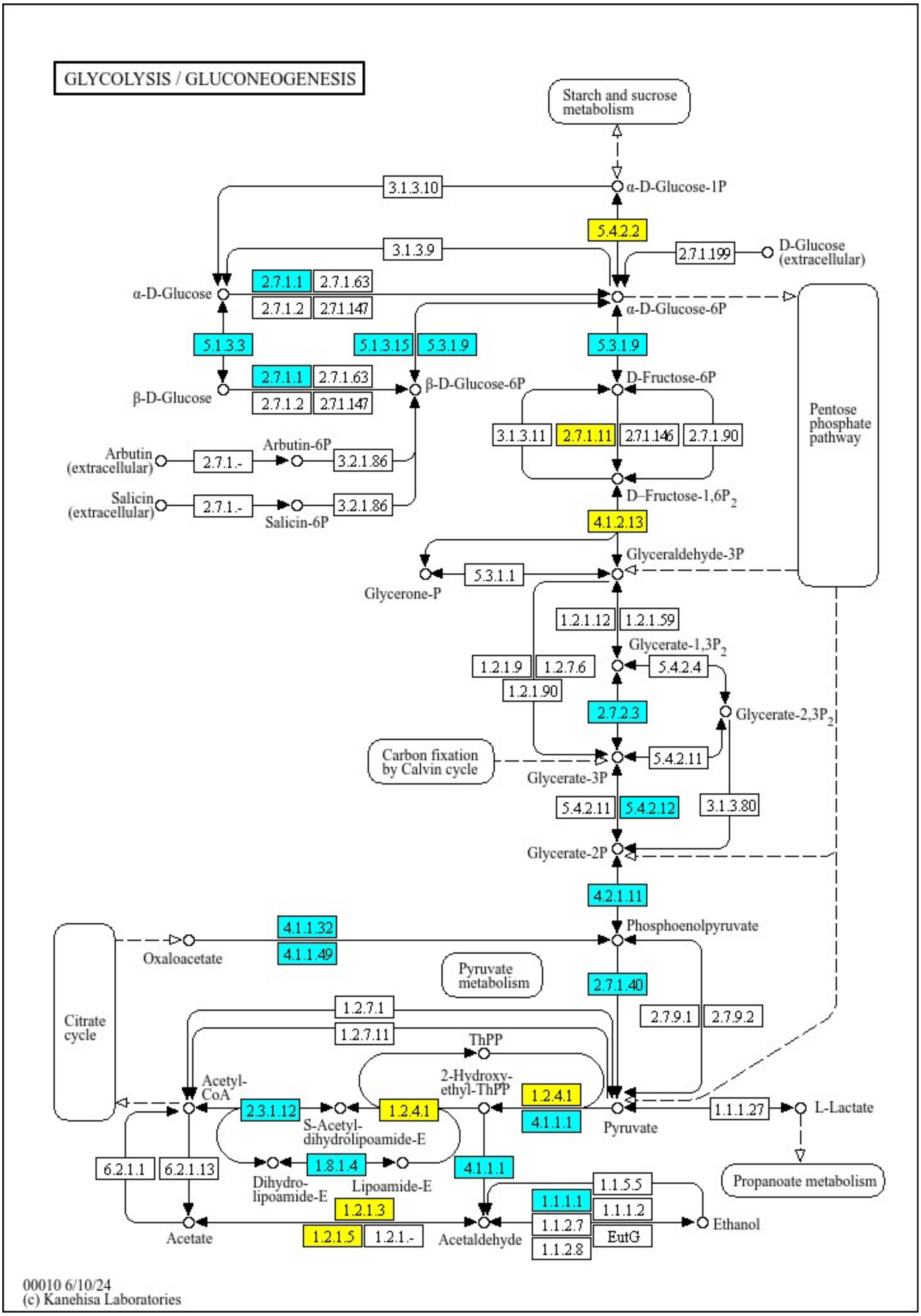

Figure S21. KEGG pathway for Glycolysis at 2 DAT. *Blue denotes down regulated genes and yellow denotes some isoforms were up regulated and some isoforms were down regulated.

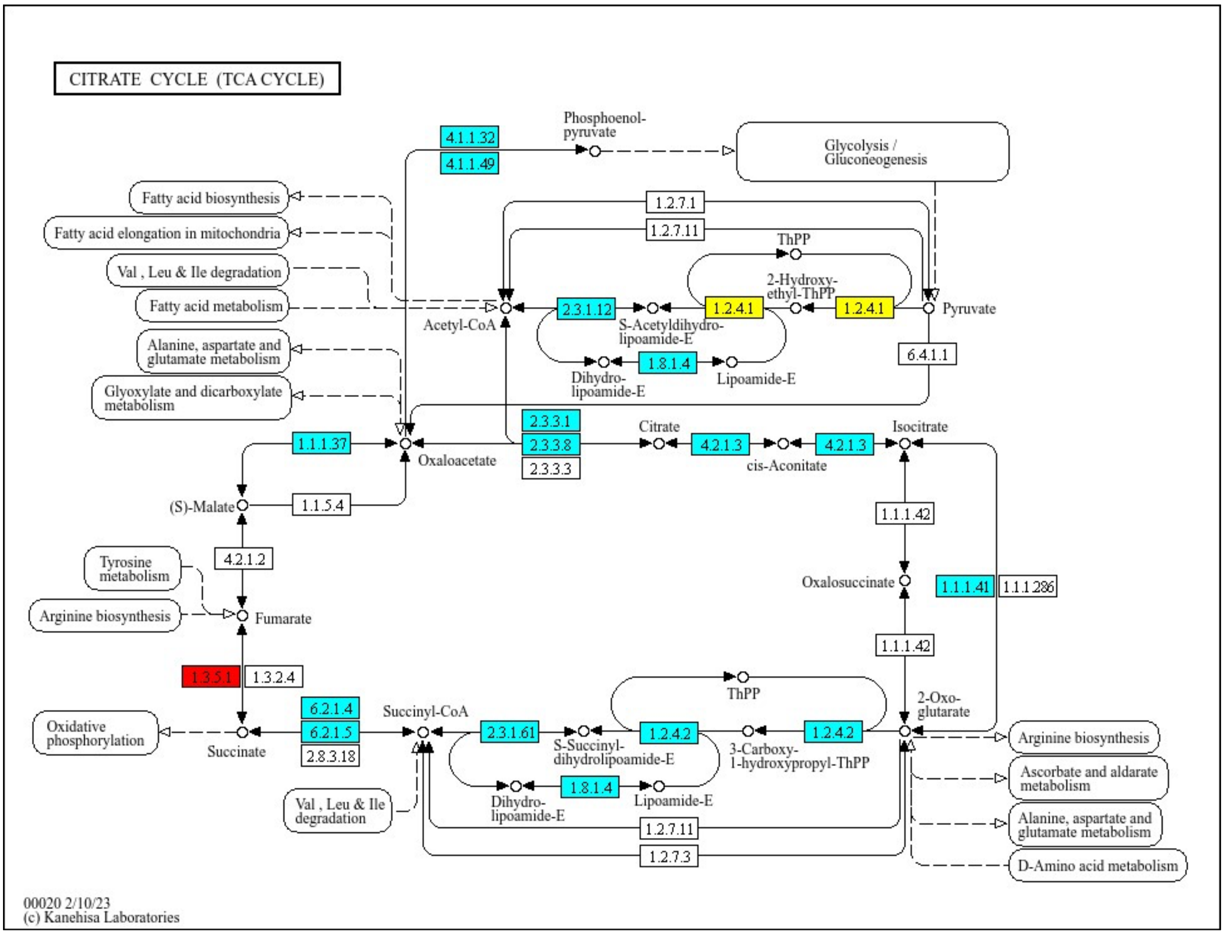

Figure S22. KEGG pathway for Citrate Cycle at 2 DAT. *Blue denotes down regulated genes, red denotes up regulated genes and yellow denotes some isoforms were up regulated and some isoforms were down regulated.

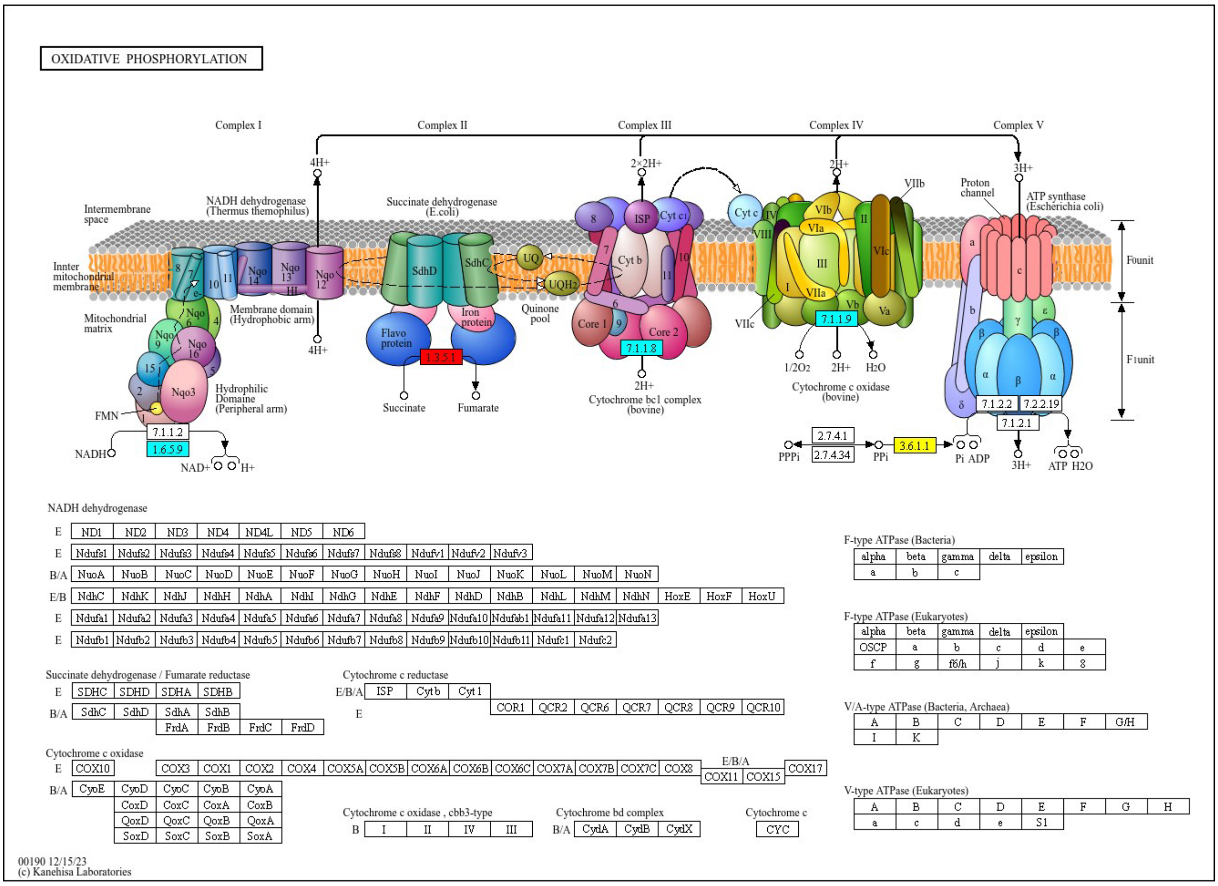

Figure S23. KEGG pathway for Oxidative Phosphorylation at 2 DAT. *Blue denotes down regulated genes, red denotes up regulated genes and yellow denotes some isoforms were up-regulated and some isoforms were down-regulated.

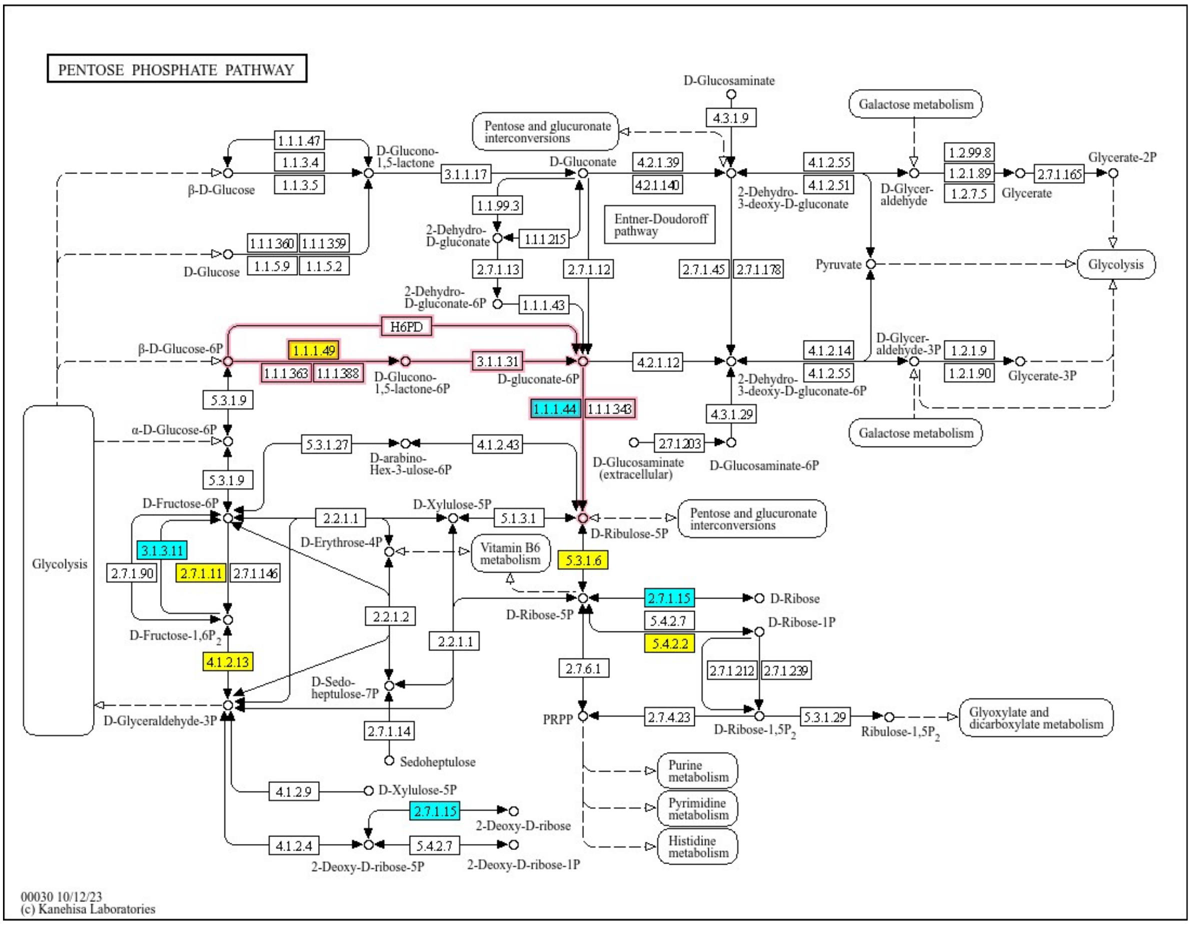

Figure S24. KEGG pathway for Pentose Phosphate Pathway (Oxidative branch) at 1 DAT. *Blue denotes down regulated genes, and yellow denotes some isoforms were up regulated and some isoforms were down regulated.

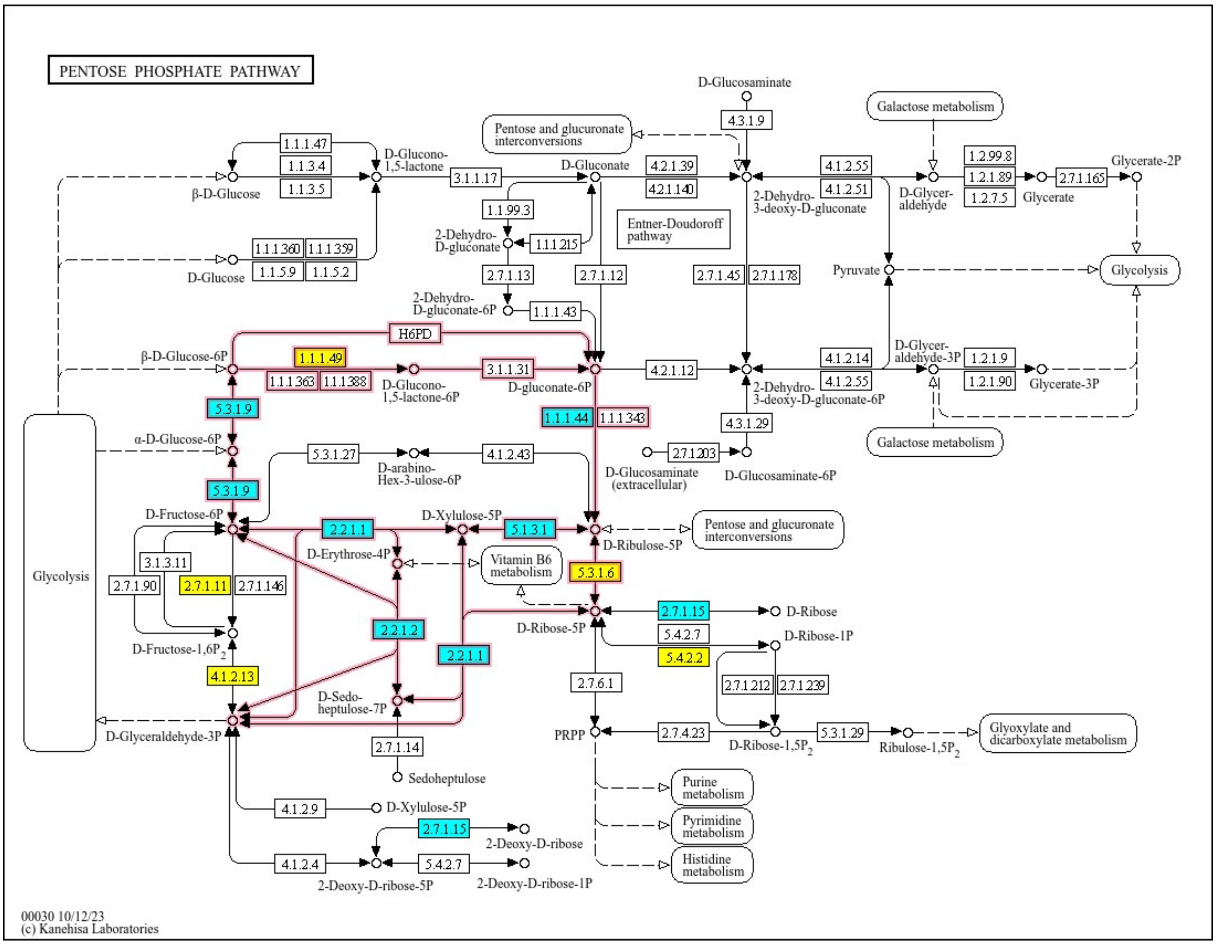

Figure S25. KEGG pathway for Pentose Phosphate Pathway (Oxidative and Non- Oxidative branch) at 2 DAT. *Blue denotes down regulated genes and yellow denotes some isoforms were up regulated and some isoforms were down regulated.

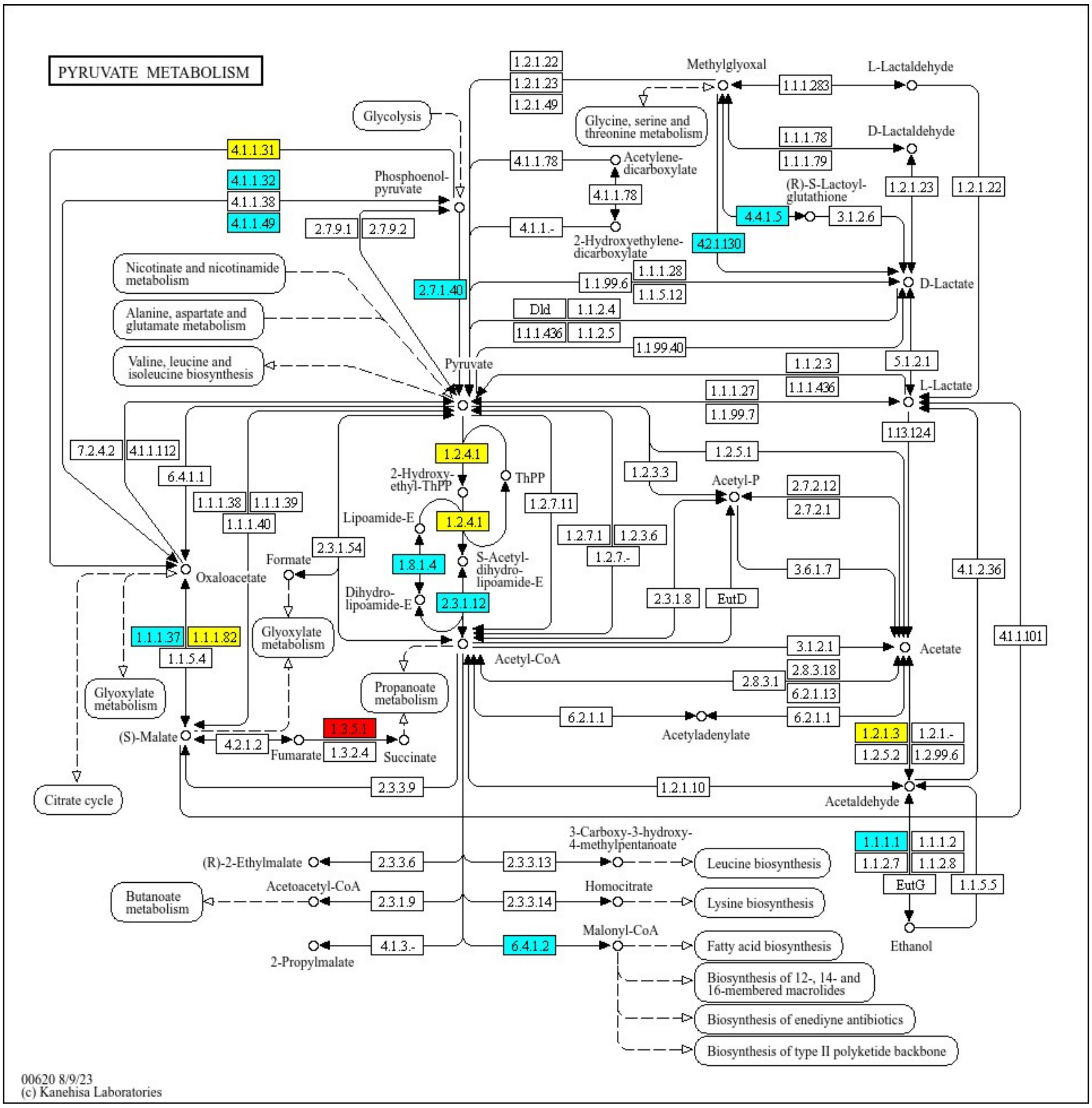

Figure S26. KEGG pathway for Pyruvate Pathway at 2 DAT. *Blue denotes down regulated genes, red denotes up regulated genes and yellow denotes some isoforms were up regulated and some isoforms were down regulated.

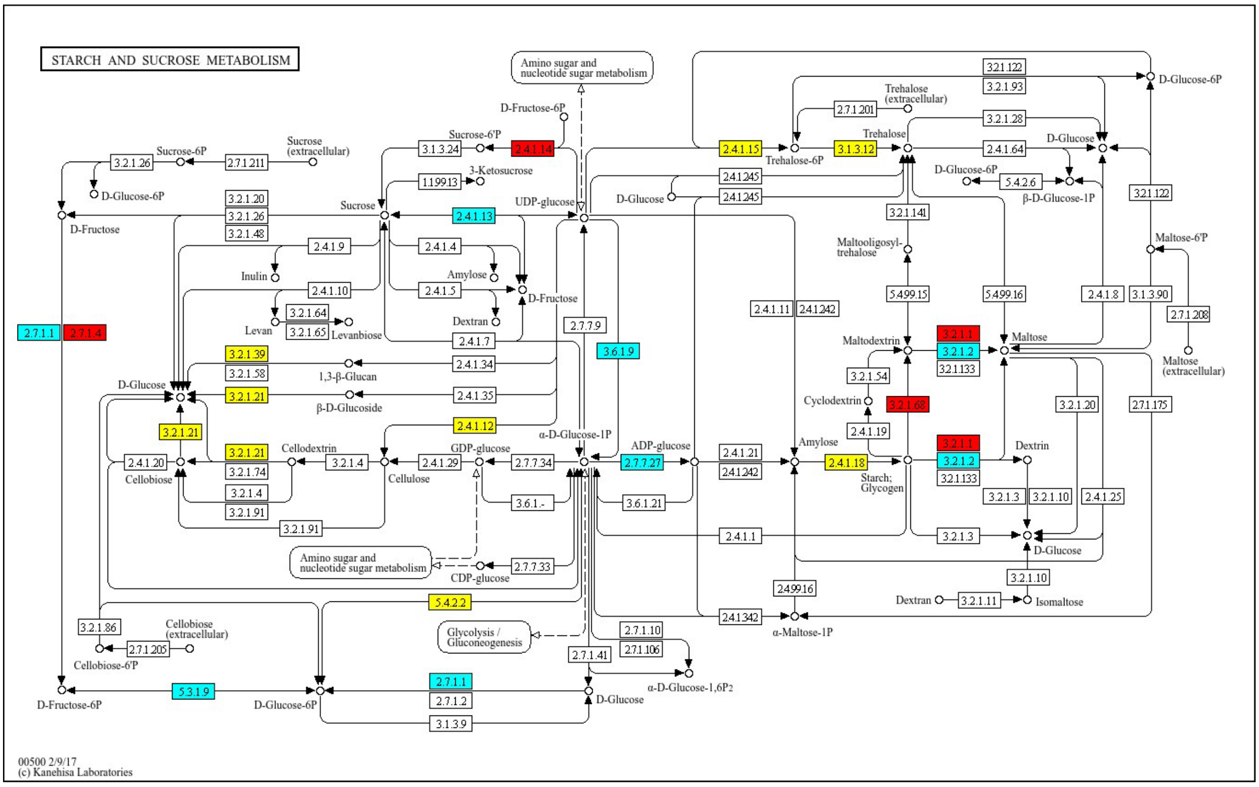

Figure S27. KEGG pathway for Starch and Sucrose metabolism at 2 DAT. *Blue denotes down regulated genes, and red denotes up regulated genes.

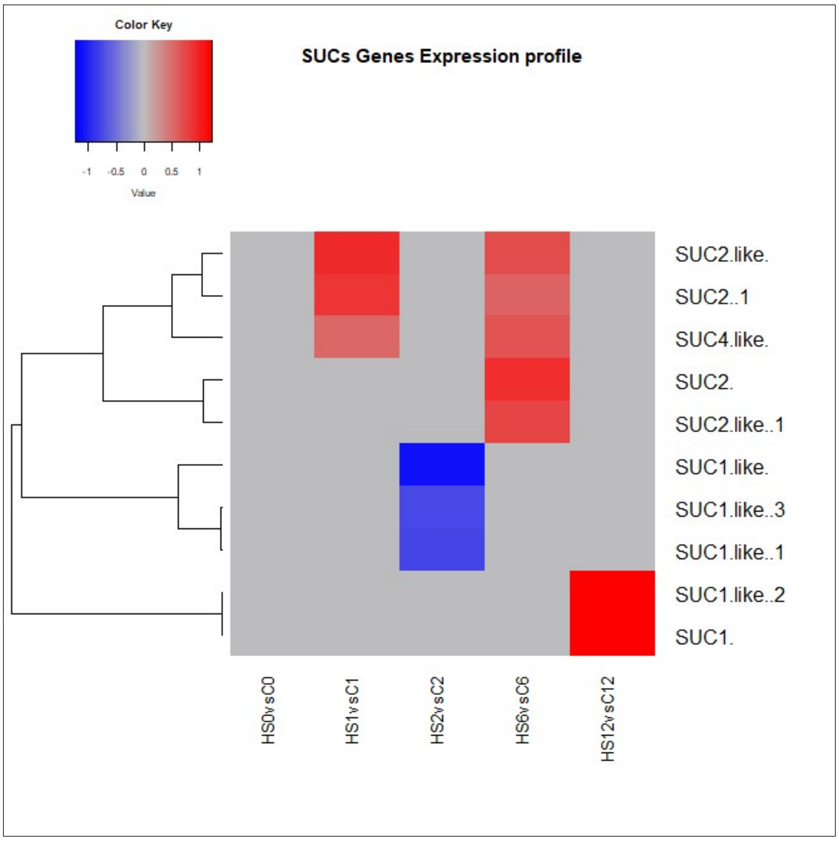

Figure S28. Heatmaps of differentially expressed Sucrose transporters genes (SUCs) at each pairwise comparison. Some transcripts encoding SUCs were found up regulated at 1 DAT and 1 & 7 DOR, while some were down regulated at 2 DAT.

Table S1. Data Quality Summary

| Sample ID | Treatment Day | Condition | Tissue | Library ID | Raw reads | Raw data | Effective(%) | Error(%) | Q20(%) | Q30(%) | GC(%) |
|---|---|---|---|---|---|---|---|---|---|---|---|
| **D0C1** | 0 | Control | Flower Buds | R11 | 42546892 | 6.4 | 98.24 | 0.03 | 96.74 | 91.33 | 43.59 |
| **D0C2** | 0 | Control | Flower Buds | R12 | 43146158 | 6.5 | 98.22 | 0.03 | 96.35 | 90.41 | 43.11 |
| **D0C3** | 0 | Control | Flower Buds | R13 | 48200214 | 7.2 | 98.23 | 0.03 | 96.63 | 91.05 | 43.06 |
| **D0C4** | 0 | Control | Flower Buds | R14 | 47204032 | 7.1 | 98.31 | 0.03 | 95.99 | 89.72 | 43.62 |
| **D0C5** | 0 | Control | Flower Buds | R15 | 46625810 | 7 | 98.33 | 0.03 | 96.13 | 89.91 | 43.29 |
| **D0HS1** | 0 | Heat stress | Flower Buds | R16 | 48823460 | 7.3 | 98.42 | 0.03 | 96.11 | 89.91 | 43.61 |
| **D0HS2** | 0 | Heat stress | Flower Buds | R17 | 42361588 | 6.4 | 98.21 | 0.03 | 95.72 | 89.14 | 43.52 |
| **D0HS3** | 0 | Heat stress | Flower Buds | R18 | 47472104 | 7.1 | 98.3 | 0.03 | 96.54 | 90.88 | 43.36 |
| **D0HS4** | 0 | Heat stress | Flower Buds | R19 | 40668522 | 6.1 | 98.3 | 0.03 | 96.35 | 90.47 | 43.35 |
| **D0HS5** | 0 | Heat stress | Flower Buds | R20 | 44058576 | 6.6 | 98.2 | 0.03 | 96.1 | 89.98 | 43.34 |
| **D1C1** | 1 | Control | Flower Buds | R21 | 44407298 | 6.7 | 98.53 | 0.03 | 96.46 | 90.65 | 44.78 |
| **D1C2** | 1 | Control | Flower Buds | R22 | 42572118 | 6.4 | 98.55 | 0.03 | 96.4 | 90.57 | 45.12 |
| **D1C3** | 1 | Control | Flower Buds | R23 | 45707162 | 6.9 | 98.54 | 0.03 | 96.65 | 91.14 | 44.99 |
| **D1C4** | 1 | Control | Flower Buds | R24 | 39779066 | 6 | 98.55 | 0.03 | 96.25 | 90.28 | 44.6 |
| **D1C5** | 1 | Control | Flower Buds | R25 | 48614394 | 7.3 | 98.21 | 0.03 | 96.3 | 90.47 | 44.96 |
| **D1HS1** | 1 | Heat stress | Flower Buds | R26 | 45844218 | 6.9 | 98.37 | 0.03 | 96.43 | 90.59 | 43.9 |
| **D1HS2** | 1 | Heat stress | Flower Buds | R27 | 47542486 | 7.1 | 98.21 | 0.03 | 96.07 | 89.82 | 44.54 |
| **D1HS3** | 1 | Heat stress | Flower Buds | R28 | 47371258 | 7.1 | 98.46 | 0.03 | 96.52 | 90.88 | 44.61 |
| **D1HS4** | 1 | Heat stress | Flower Buds | R29 | 46407612 | 7 | 98.58 | 0.03 | 96.58 | 91.05 | 44.59 |
| **D1HS5** | 1 | Heat stress | Flower Buds | R30 | 42705194 | 6.4 | 98.7 | 0.03 | 95.71 | 89.03 | 44.59 |
| **D2C1** | 2 | Control | Flower Buds | R31 | 48709190 | 7.3 | 98.45 | 0.03 | 96.39 | 90.54 | 45.57 |
| **D2C2** | 2 | Control | Flower Buds | R32 | 50232808 | 7.5 | 98.68 | 0.03 | 96.24 | 90.25 | 45.61 |
| **D2C3** | 2 | Control | Flower Buds | R33 | 42394356 | 6.4 | 98.72 | 0.03 | 96.86 | 91.66 | 45.25 |
| **D2C5** | 2 | Control | Flower Buds | R35 | 49019624 | 7.4 | 98.26 | 0.03 | 96.62 | 90.98 | 45.2 |
| **D2HS1** | 2 | Heat stress | Flower Buds | R36 | 43939042 | 6.6 | 98.46 | 0.03 | 96.65 | 91.08 | 45.12 |
| **D2HS2** | 2 | Heat stress | Flower Buds | R37 | 45232502 | 6.8 | 98.6 | 0.03 | 96.83 | 91.45 | 44.77 |
| **D2HS3** | 2 | Heat stress | Flower Buds | R38 | 41291490 | 6.2 | 98.43 | 0.03 | 96.6 | 90.94 | 44.93 |
| **D2HS4** | 2 | Heat stress | Flower Buds | R39 | 44141240 | 6.6 | 98.52 | 0.03 | 96.57 | 90.87 | 44.94 |
| **D2HS5** | 2 | Heat stress | Flower Buds | R40 | 40421822 | 6.1 | 98.54 | 0.03 | 96.48 | 90.68 | 44.36 |
| **D6C1** | 6 | Control | Flower Buds | R41 | 45618730 | 6.8 | 98.3 | 0.03 | 96.09 | 89.94 | 43.28 |
| **D6C2** | 6 | Control | Flower Buds | R42 | 47279800 | 7.1 | 98.36 | 0.03 | 96.88 | 91.63 | 43.5 |
| **D6C3** | 6 | Control | Flower Buds | R43 | 49797014 | 7.5 | 98.42 | 0.03 | 96.2 | 90.13 | 43.04 |
| **D6C4** | 6 | Control | Flower Buds | R44 | 48275232 | 7.2 | 98.45 | 0.03 | 95.79 | 89.21 | 43.72 |
| **D6C5** | 6 | Control | Flower Buds | R45 | 41275426 | 6.2 | 98.51 | 0.03 | 95.91 | 89.47 | 43.53 |
| **D6HS1** | 6 | Heat stress | Flower Buds | R46 | 45090682 | 6.8 | 98.26 | 0.03 | 96.04 | 89.81 | 43.4 |
| **D6HS2** | 6 | Heat stress | Flower Buds | R47 | 46214376 | 6.9 | 97.73 | 0.03 | 96.51 | 90.89 | 43.35 |
| **D6HS3** | 6 | Heat stress | Flower Buds | R48 | 44791070 | 6.7 | 98.28 | 0.03 | 96.38 | 90.48 | 43.85 |
| **D6HS4** | 6 | Heat stress | Flower Buds | R49 | 43154676 | 6.5 | 98.09 | 0.03 | 95.9 | 89.59 | 43.06 |
| **D6HS5** | 6 | Heat stress | Flower Buds | R50 | 48451812 | 7.3 | 98.31 | 0.03 | 96.35 | 90.49 | 43.31 |
| **D12C1** | 12 | Control | Flower Buds | R51 | 42706732 | 6.4 | 98.61 | 0.03 | 96.19 | 90.2 | 45.62 |
| **D12C2** | 12 | Control | Flower Buds | R52 | 45100386 | 6.8 | 98.52 | 0.03 | 97.7 | 93.57 | 45.32 |
| **D12C3** | 12 | Control | Flower Buds | R53 | 49761388 | 7.5 | 98.61 | 0.03 | 97.43 | 92.94 | 45.52 |
| **D12C4** | 12 | Control | Flower Buds | R54 | 55353976 | 8.3 | 98.67 | 0.03 | 97.45 | 92.98 | 45.8 |
| **D12C5** | 12 | Control | Flower Buds | R55 | 43229896 | 6.5 | 98.7 | 0.03 | 97.49 | 93 | 45.51 |
| **D12HS1** | 12 | Heat stress | Flower Buds | R56 | 39276294 | 5.9 | 98.68 | 0.03 | 97.39 | 92.84 | 44.61 |
| **D12HS1** | 12 | Heat stress | Flower Buds | R57 | 43958400 | 6.6 | 98.8 | 0.03 | 97.61 | 93.31 | 44.4 |
| **D12HS2** | 12 | Heat stress | Flower Buds | R58 | 49089438 | 7.4 | 98.67 | 0.03 | 97.43 | 92.92 | 44.69 |
| **D12HS3** | 12 | Heat stress | Flower Buds | R59 | 43623028 | 6.5 | 98.71 | 0.03 | 97.54 | 93.2 | 44.56 |
| **D12HS4** | 12 | Heat stress | Flower Buds | R60 | 42641344 | 6.4 | 98.62 | 0.03 | 97.47 | 93.04 | 44.27 |

Table S2. PCR timepoints tested per gene

| Timepoints / Genes | 0 (baseline) | 1 DAT (24 hrs of HS*) | 2 DAT (48 hrs of HS) | 5 DAT (5 days of HS) | 1 DOR (1 day of recovery) | 7 DOR (7 days of recovery) |
|---|---|---|---|---|---|---|
| **ACTIN** | ✓ | ✓ | ✓ | ✓ | ✓ | ✓ |
| **SS5** | ✓ | X | ✓ | ✓ | ✓ | ✓ |
| **HSP20** | ✓ | X | ✓ | ✓ | ✓ | ✓ |

* HS: Heat Stress

Table S3. List of primer sequences of 3 selected genes and reference gene used for the qt-PCR analysis

| Primer name | Forward primer sequence ((5' to 3') | Reverse primer sequence ((5' to 3') |
|---|---|---|
| **BnaSS5** | GGAGAAAACGCAGGGAAAG | TCAGGCAAACCAAGAACATC |
| **BnaHSP20** | CAAGGAGTATCAGCCAGGTG | TAGAGGGCATCGTCTTTCTC |
| **BnaACTIN** | TGGGTTTGCTGGTGACGAT | TGCCTAGGACGACCAACAATACT |

Table S4.Correlation analysis of the results obtained from RNA-seq and qt-PCR for SS5, HSP20 & BRCA1 genes at different timepoints.

| Log2 qt-PCR & RNA-seq values | | | | |
|---|---|---|---|---|
| | **SS5** | | **HSP20** | |
| **Timepoints** | **Qt-PCR** | **RNA-seq** | **Qt-PCR** | **RNA-seq** |
| HS0vsC0 | 0.141107 | 0 | 6.981295 | 6.9 |
| HS1vsC1 | | 0 | | 6.7 |
| HS2vsC2 | -0.48492 | -1.1 | 8.550429 | 5.7 |
| HS5vsC5 | 0.163089 | | 5.673896 | |
| HS6vsC6 | 0.814462 | 0 | 3.146389 | 5.3 |
| HS12vsC12 | 0.397126 | 0 | 2.671107 | 0 |
| R | 0.860104 | | 0.681909 | |
| R2 | 0.73978 | | 0.464999 | |

Table S5. DE of Sucrose synthesizing enzyme (Sucrose Phosphate Synthase)

| Gene | HS0vsC0 | HS1vsC1 | HS2vsC2 | HS6vsC6 | HS12vsC12 | Seq. Description | Enz. Code |
|---|---|---|---|---|---|---|---|
| BnaA06G0367000WE | 0 | 0.69 | 0 | 0.9 | 0.7 | Probable sucrose-phosphate synthase 2 | EC:2.4.1.14 |
| BnaC02G0093600WE | 0.5 | 0.76 | 0.77 | 0 | 0 | Sucrose-phosphate synthase 1-like | EC:2.4.1.14 |
| BnaC07G0352200WE | 0.8 | 1.1 | 1.0 | 1.4 | 0.94 | Probable sucrose-phosphate synthase 2 | EC:2.4.1.14 |
| BnaA02G0084400WE | 0 | -0.5 | 0 | 0 | 0 | sucrose-phosphate synthase 1-like | EC:2.4.1.14 |
| BnaA10G0171700WE | -0.52 | 0 | 0 | 0 | 0 | sucrose-phosphate synthase 1 | EC:2.4.1.14 |

Table S6. DE of Sucrose hydrolysing enzymes (Cell Wall Invertase and Sucrose Synthase)

| Gene | HS0vsC0 | HS1vsC1 | HS2vsC2 | HS6vsC6 | HS12vsC12 | Seq. Description | Enz. Code |
|---|---|---|---|---|---|---|---|
| BnaA03G0187500WE | -1.1 | 0 | -1.2 | 0 | 0 | CWINV4-1; beta-fructofuranosidase | EC:3.2.1.26 |
| BnaC01G0375500WE | 0 | 0 | 0 | 1.7 | 0 | CWINV1; beta-fructofuranosidase | EC:3.2.1.26 |
| BnaC03G0139900WE | 0 | 0 | -1.7 | 0 | 0 | CWINV4-1; beta-fructofuranosidase | EC:3.2.1.26 |
| BnaC05G0441400WE | 0.9 | 0 | 0 | 1.5 | 0 | CWINV5-2; beta-fructofuranosidase | EC:3.2.1.26 |
| BnaA05G0141800WE | 0 | 0 | -1.19 | 0 | 0 | Sucrose synthase 5 | EC:2.4.1.13 |
| BnaA07G0333600WE | 0 | 0 | -1.45 | 0 | 0 | Sucrose synthase 6 | EC:2.4.1.13 |
| BnaA09G0016500WE | 0 | 0 | 0 | -1.04 | 0 | Sucrose synthase 3 | EC:2.4.1.13 |
| BnaC06G0143400WE | 0 | 0 | -1.16 | 0 | 0 | Sucrose synthase 5 | EC:2.4.1.13 |
| BnaA10G0167800WE | 0 | 0 | 0 | 0 | -0.8 | Sucrose synthase -1 like | EC:2.4.1.13 |

Table S7. LogFC of top 30 up- regulated genes at 1 DAT as obtained by DESeq2 pipeline

| Gene | HS0vsC0 | HS1vsC1 | HS2vsC2 | HS6vsC6 | HS12vsC12 | Gene Annotation |
|---|---|---|---|---|---|---|
| BnaA08G0138700WE | 25.50 | 23.04 | 0.00 | 23.20 | 0.00 | -- |
| Bnascaffold2770G0000100WE | 0.00 | 19.40 | 22.96 | -22.29 | 0.00 | probable ATP-dependent DNA helicase CHR12; K11647 SWI/SNF-related matrix-associated actin-dependent regulator of chromatin subfamily A member 2/4 [EC:3.6.4.-] (A) |
| BnaC03G0236200WE | 10.03 | 10.84 | 11.03 | 5.90 | 0.00 | HSP22; 22.0 kDa heat shock protein; K13993 HSP20 family protein (A |
| BnaA06G0312000WE | 7.11 | 7.79 | 6.90 | 0.00 | 0.00 | 23.6 kDa heat shock protein mitochondrial-like; K13993 HSP20 family protein (A) |
| BnaA01G0068500WE | 7.69 | 7.31 | 6.13 | 6.23 | 0.00 | 23.6 kDa heat shock protein mitochondrial-like; K13993 HSP20 family protein (A) |
| BnaC06G0410600WE | 5.13 | 6.55 | 7.80 | 4.72 | 1.58 | Developmentally- regulated G-protein 2; K06944 uncharacterized protein (A) |
| BnaC02G0043400WE | 7.05 | 5.57 | 6.31 | 4.49 | 0.00 | 17.6 kDa class II heat shock protein; K13993 HSP20 family protein (A) |
| BnaC05G0190700WE | 4.43 | 5.55 | 5.20 | 3.40 | 0.00 | p23; uncharacterized protein OsI_027940-like; K15730 cytosolic prostaglandin-E synthase [EC:5.3.99.3] (A) |
| BnaA06G0177800WE | 6.47 | 5.33 | 4.96 | 5.62 | 0.00 | BcHSP17.6; 17.4 kDa class I heat shock protein; K13993 HSP20 family protein (A |
| BnaA09G0429400WE | 4.48 | 5.32 | 5.66 | 4.47 | 0.00 | p23; uncharacterized protein OsI_027940-like; K15730 cytosolic prostaglandin-E synthase [EC:5.3.99.3] (A) |
| BnaA10G0226400WE | 4.76 | 5.14 | 4.92 | 3.19 | 0.00 | 17.6 kDa class II heat shock protein-like; K13993 HSP20 family protein (A) |
| BnaA08G0073800WE | 4.06 | 4.74 | 4.64 | 3.27 | 2.13 | zinc finger protein ZPR1-like; K06874 zinc finger protein (A) |
| Bnascaffold3043G0000200WE | 4.54 | 4.42 | 4.78 | 3.69 | 0.00 | 17.6 kDa class II heat shock protein; K13993 HSP20 family protein (A) |
| BnaC09G0508300WE | 4.01 | 4.34 | 3.71 | 2.14 | 0.00 | elongation factor 1-beta 1-like; K03232 elongation factor 1-beta (A) |
| BnaC08G0070600WE | 3.19 | 4.28 | 3.79 | 2.48 | 1.29 | zinc finger protein ZPR1-like; K06874 zinc finger protein (A) |
| BnaC06G0424500WE | 3.26 | 3.73 | 3.33 | 2.50 | 0.00 | chaperone protein ClpB1; K03695 ATP-dependent Clp protease ATP-binding subunit ClpB (A) |
| BnaA06G0328200WE | 2.62 | 3.58 | 3.15 | 1.54 | 0.00 | -- |
| BnaA03G0160300WE | 0.00 | 3.43 | 2.66 | 0.00 | 0.00 | potassium transporter 1; K03549 KUP system potassium uptake protein (A) |
| BnaC06G0380500WE | 0.00 | 3.28 | 5.35 | 3.87 | 5.65 | hypothetical protein; K17803 methyltransferase OMS1 mitochondrial [EC:2.1.1.-] (A |
| BnaC01G0386900WE | 3.48 | 3.18 | 3.32 | 2.85 | 0.00 | probable mediator of RNA polymerase II transcription subunit 37c; K03283 heat shock 70kDa protein 1/8 (A) |
| BnaA07G0329100WE | 3.52 | 3.11 | 3.11 | 3.09 | 0.00 | developmentally- regulated G-protein 2; K06944 uncharacterized protein (A) |
| BnaA07G0341000WE | 2.98 | 2.99 | 2.76 | 2.65 | 0.00 | chaperone protein ClpB1; K03695 ATP-dependent Clp protease ATP-binding subunit ClpB (A) |
| BnaC04G0385500WE | 2.89 | 2.92 | 3.33 | 2.29 | 0.00 | 40S ribosomal protein S9-2; K02997 small subunit ribosomal protein S9e (A) |
| BnaA06G0148400WE | 0.00 | 2.86 | 1.76 | 0.00 | 0.00 | -- |
| BnaA01G0210500WE | 2.32 | 2.84 | 2.12 | 2.75 | 1.26 | pre-mRNA-splicing factor SLU7-like; K12819 pre-mRNA-processing factor SLU7 (A) |
| BnaA02G0000800WE | 0.00 | 2.83 | 10.65 | 0.00 | 11.21 | pyridoxal biosynthesis protein PDX1.3; K06215 pyridoxal 5'-phosphate synthase pdxS subunit [EC:4.3.3.6] (A) |
| BnaA05G0117900WE | 2.47 | 2.79 | 2.82 | 1.66 | 0.00 | heat shock 70 kDa protein 8-like; K09489 heat shock 70kDa protein 4 (A) |
| BnaA03G0022900WE | 1.94 | 2.70 | 1.93 | 2.25 | 0.00 | transcription factor bHLH100-like; K18486 heart-and neural crest derivatives-expressed protein 2 (A) |
| BnaA07G0306600WE | 1.94 | 2.70 | 1.93 | 2.25 | 0.00 | transcription factor bHLH100-like; K18486 heart-and neural crest derivatives-expressed protein 2 (A) |
| BnaA05G0366100WE | 0.00 | 2.68 | 4.28 | 0.00 | 0.00 | chaperonin 60 subunit beta 2 chloroplastic; K04077 chaperonin GroEL (A) |

Table S8. LogFC of top 30 down- regulated genes at 1 DAT as obtained by DESeq2 pipeline

| Gene | HS0vsC0 | HS1vsC1 | HS2vsC2 | HS6vsC6 | HS12vsC12 | Gene Annotation |
|---|---|---|---|---|---|---|
| BnaA01G0299900WE | 0.00 | -25.04 | -24.77 | 0.00 | 0.00 | probable ATP-dependent DNA helicase CHR12; K11647 SWI/SNF-related matrix-associated actin-dependent regulator of chromatin subfamily A member 2/4 [EC:3.6.4.-] (A) |
| BnaA02G0390800WE | 0.00 | -24.10 | 0.00 | 0.00 | 0.00 | 50S ribosomal protein L22-like; K02890 large subunit ribosomal protein L22 (A) |
| Bnascaffold390G0000100WE | 0.00 | -22.84 | 23.50 | 0.00 | 0.00 | BTB/POZ and MATH domain-containing protein 2-like; K10523 speckle-type POZ protein (A) |
| BnaA09G0548300WE | 0.00 | -21.05 | 0.00 | 0.00 | 0.00 | serine/threonine-protein phosphatase 2A 65 kDa regulatory subunit A beta isoform-like; K03456 serine/threonine-protein phosphatase 2A regulatorysubunit A (A) |
| Bnascaffold2041G0000500WE | 0.00 | -10.51 | 0.00 | 0.00 | -9.64 | solute carrier family 2 facilitated glucose transporter member 8-like; K08145 MFS transporter |
| BnaC03G0531800WE | 0.00 | -10.02 | 0.00 | 0.00 | 0.00 | asparagine synthetase [glutamine-hydrolyzing]; K01953 asparagine synthase (glutamine-hydrolysing) [EC:6.3.5.4] (A) |
| BnaA05G0410300WE | 0.00 | -7.40 | 8.07 | 0.00 | 0.00 | cell division control protein 48-like; K08900 mitochondrial chaperone BCS1 (A) |
| BnaC03G0161100WE | 0.00 | -5.64 | -7.24 | -6.53 | 0.00 | bidirectional sugar transporter SWEET9; K15382 solute carrier family 50 (sugar transporter) (A) |
| BnaA08G0122000WE | 0.00 | -5.09 | 0.00 | 0.00 | 0.00 | endo-1 4-beta-xylanase C-like; K01181 endo-1 |
| BnaC06G0349400WE | 0.00 | -4.83 | 0.00 | 0.00 | 0.00 | Peptidase putative (EC:3.4.21.25); K18443 golgi-specific brefeldin A-resistance guanine nucleotide exchange factor 1 (A) |
| BnaA05G0285000WE | 0.00 | -4.25 | -3.68 | -4.25 | 0.00 | homocysteine S-methyltransferase 3; K00547 homocysteine S-methyltransferase [EC:2.1.1.10] (A) |
| BnaC02G0269700WE | 0.00 | -3.98 | 0.00 | 0.00 | 0.00 | strictosidine synthase 1-like; K01757 strictosidine synthase [EC:4.3.3.2] (A) |
| BnaC09G0033200WE | -3.53 | -3.80 | -6.65 | -5.10 | 0.00 | -- |
| BnaC03G0440400WE | 0.00 | -3.43 | 0.00 | 0.00 | 0.00 | geranylgeranyl diphosphate reductase chloroplastic; K10960 geranylgeranyl reductase [EC:1.3.1.83] (A) |
| BnaA08G0177000WE | 0.00 | -3.41 | 0.00 | 0.00 | 0.00 | Glutamyl-tRNA(Gln) amidotransferase subunit A; K02433 aspartyl-tRNA(Asn)/glutamyl-tRNA (Gln) amidotransferase subunit A [EC:6.3.5.6 6.3.5.7] (A) |
| BnaA03G0270900WE | 0.00 | -3.27 | -3.37 | -3.13 | 0.00 | uncharacterized LOC103858630; K14488 SAUR family protein (A) |
| BnaC05G0336600WE | -2.85 | -3.01 | -3.35 | -2.72 | 0.00 | homocysteine S-methyltransferase 3-like; K00547 homocysteine S-methyltransferase [EC:2.1.1.10] (A) |
| BnaC02G0072900WE | 0.00 | -2.95 | -3.61 | -3.90 | 0.00 | homeobox-leucine zipper protein HDG9-like; K09338 homeobox-leucine zipper protein (A) |
| BnaA03G0328400WE | -3.61 | -2.91 | -3.15 | -3.01 | 0.00 | alanine--glyoxylate aminotransferase 2 homolog 3 mitochondrial; K00827 alanine-glyoxylate transaminase / (R)-3-amino-2-methylpropionate-pyruvate transaminase [EC:2.6.1.44 2.6.1.40] (A) |
| BnaA05G0069600WE | -2.45 | -2.84 | -2.23 | -1.88 | 0.00 | ribonucleoprotein At2g37220 chloroplastic-like; K11294 nucleolin (A) |
| BnaC01G0410600WE | 0.00 | -2.82 | -4.14 | -3.26 | 0.00 | cytosolic sulfotransferase 11-like; K01016 estrone sulfotransferase [EC:2.8.2.4] (A) |
| BnaC04G0129500WE | -2.72 | -2.79 | 0.00 | -2.46 | 0.00 | peroxidase 19; K00430 peroxidase [EC:1.11.1.7] (A) |
| BnaA06G0144700WE | -2.08 | -2.73 | -2.42 | -2.29 | 0.00 | auxin:hydrogen symporter putative; K07088 uncharacterized protein (A) |
| BnaC07G0304700WE | -1.57 | -2.71 | -3.10 | -1.92 | 0.00 | bidirectional sugar transporter SWEET4-like; K15382 solute carrier family 50 (sugar transporter) (A) |
| BnaC03G0244200WE | 0.00 | -2.71 | -2.35 | -3.11 | 0.00 | auxin-responsive protein SAUR36-like; K14488 SAUR family protein (A) |
| BnaA03G0313000WE | 0.00 | -2.70 | 0.00 | 0.00 | 0.00 | -- |
| BnaA02G0006300WE | -3.95 | -2.69 | -4.98 | -3.01 | 0.00 | -- |
| BnaC04G0003100WE | -3.13 | -2.66 | 0.00 | 0.00 | 0.00 | probable pectinesterase/pectinesterase inhibitor 20; K01051 pectinesterase [EC:3.1.1.11] (A) |
| BnaA04G0096200WE | 0.00 | -2.56 | -2.12 | 0.00 | 0.00 | proline dehydrogenase 2 mitochondrial; K00318 proline dehydrogenase [EC:1.5.-.-] (A) |
| BnaC02G0359700WE | -2.67 | -2.53 | -3.28 | 0.00 | 0.00 | BAHD acyltransferase DCR; K19747 BAHD acyltransferase [EC:2.3.1.-] (A) |
